# A *de novo* protein structure prediction by iterative partition sampling, topology adjustment, and residue-level distance deviation optimization

**DOI:** 10.1101/2021.05.12.443769

**Authors:** Jun Liu, Kai-Long Zhao, Guang-Xing He, Liu-Jing Wang, Xiao-Gen Zhou, Gui-Jun Zhang

## Abstract

**Motivation:** With the great progress of deep learning-based inter-residue contact/distance prediction, the discrete space formed by fragment assembly cannot satisfy the distance constraint well. Thus, the optimal solution of the continuous space may not be achieved. Designing an effective closed-loop continuous dihedral angle optimization strategy that complements the discrete fragment assembly is crucial to improve the performance of the distance-assisted fragment assembly method.

**Results:** In this article, we proposed a de novo protein structure prediction method called IPTDFold based on closed-loop iterative partition sampling, topology adjustment and residue-level distance deviation optimization. First, local dihedral angle crossover and mutation operators are designed to explore the conformational space extensively and achieve information exchange between the conformations in the population. Then, the dihedral angle rotation model of loop region with partial inter-residue distance constraints is constructed, and the rotation angle satisfying the constraints is obtained by differential evolution algorithm, so as to adjust the spatial position relationship between the secondary structures. Lastly, the residue distance deviation is evaluated according to the difference between the conformation and the predicted distance, and the dihedral angle of the residue is optimized with biased probability. The final model is generated by iterating the above three steps. IPTDFold is tested on 462 benchmark proteins, 24 FM targets of CASP13, and 20 FM targets of CASP14. Results show that IPTDFold is significantly superior to the distance-assisted fragment assembly method Rosetta_D (Rosetta with distance). In particular, the prediction accuracy of IPTDFold does not decrease as the length of the protein increases. When using the same *FastRelax* protocol, the prediction accuracy of IPTDFold is significantly superior to that of trRosetta without orientation constraints, and is equivalent to that of the full version of trRosetta.

**Availability:** The source code and executable are freely available at https://github.com/iobio-zjut/IPTDFold.

## 1 Introduction

With the introduction of deep residual neural networks into inter-residue contact prediction by Xu (Wang *et al*., 2017), the application of deep learning in inter-residue contact/distance prediction has achieved great success in recent CASPs (Wang *et al*., 2018; Xu and Wang, 2019; Kandathil *et al*., 2019; Li *et al*., 2019; Mao *et al*., 2020;Yang *et al*., 2020), thus promoting a breakthrough in protein structure prediction (Zheng *et al*., 2019; Xu, 2019; Greener *et al*., 2019; Yang *et al*., 2020; Senior *et al*., 2020; Moult *et al*., 2018). In general, de novo protein structure prediction using predicted distance is mainly divided into two categories: distance-assisted fragment assembly and geometric-constrained energy minimization (AlQuraishi, 2019; Kuhlman and Bradley, 2019). CNS (Brunger *et al*., 2007) and gradient descent are two commonly used geometric-constrained energy minimization methods. EVfold (Marks *et al*., 2011), CONFOLD (Adhikari *et al*., 2015; Adhikariand Cheng, 2018), RaptorX (Xu, 2019; Xu and Wang, 2019), and DMPfold (Greener *et al*., 2019) feed the predicted contact/distance and other constraints into the CNS to generate a model. AlphaFold (Senior*et al*., 2020) and trRosetta (Yang *et al*., 2020) convert the distance and/or orientation distribution into a protein-specific statistical potential function, and generate the model through gradient descent. Gradient descent optimization is effective at finding the nearest local minimum in the energy landscape, but generally, it will not locate the global minimum (Kuhlman and Bradley, 2019). Therefore, multiple minimization protocols need to be initiated to generate the lowest-potential model. Moreover, there are still challenges in accurately predicting protein contact distance when there are few homologous sequences, folding proteins from noisy contact distance (Hou *et al*., 2019).

The results of recent CASPs show that the distance-assisted fragment assembly is still one of the most competitive protein structure prediction methods (Zheng *et al*., 2019; Kryshtafovych *et al*., 2019; Zhang *et al*., 2020). In the distance-assisted fragment assembly methods, contact/distance is used as an energy term or constraint combined with a classical energy function to guide folding. Generally, candidate conformations are generated by fragment assembly and other sampling strategies, and then evaluated by the energy function and contact/distance. Afterward, the conformation is updated according to Metropolis criterion (Metropolis *et al*., 1953). FRAGFOLD adds the predicted contact to the existing energy function (Jones *et al*., 2005) and assembles a mixture of super secondary structural fragments and short-fixed length fragments through simulated annealing to generate a 3D structure (Kosciolek and Jones, 2014). PconsFold applies contacts into the Rosetta AbinitioRelax folding protocol to generate structural models through fragment assembly (Michel *et al*., 2014). CoDiFold combines the contacts from two different servers and fragment-derived distance profile into the Rosetta coarse-grained energy function, and predicts the structure model through fragment assembly and evolutionary algorithms (Peng *et al*., 2020). In CASP13, C-QUARK designed a composite energy function by combining the contact potential, the fragment-derived distance profile restraints and the inherent knowledge-based energy function. The composite energy function is used to guide the assembly of the fragments into full structural models by replica-exchange Monte Carlo simulations (Xu and Zhang, 2012; Zheng *et al*., 2019). In the recent CASP14, D-QUARK, which integrates the predicted distance, is proposed (Zhang *et al*., 2020). Fragment assembly converts the continuous dihedral space into discrete experimental fragment combinatorial optimization to effectively reduce the search space. However, some potential conformational spaces cannot be sampled due to the limitations of the fragment library, especially its inability to sufficiently sample the flexible loop area (Heoetal.,2017; Liuetal.,2020). In addition, because fragment insertion moves tend to perturb long-range contacts, fragment assembly does not work well for complex, non-local topological proteins in the absence of predicted or experimentally determined residue contact information (Kuhlman and Bradley, 2019).

Therefore, how to effectively use predicted distance and design a new folding strategy is crucial to improve the prediction accuracy of distance-assisted fragment assembly method. In this article, we proposed a protein structure prediction algorithm IPTDFold, which contains three modules: partition sampling based on random fragment insertion and local structure crossover and mutation; topology adjustment by partial distance constrained loop-specific dihedral angle sampling; and residue-level distance deviation optimization based on the difference of the conformation and the predicted distance. The structure model is predicted by iterating the three modules. The predicted distance is constructed as a potential and combined with the energy function to guide folding. Experimental results show that IPTDFold significantly improves the prediction accuracy of the distance-assisted fragment assembly method and reaches the accuracy level of the state-of-the-art gradient descent energy minimization method.

## 2 Methods

The pipeline of IPTDFold is shown in **Figure 1**. In addition to the query sequence, the inter-residue distance and the fragment library are required as input to IPTDFold. Instead of running thousands of folding simulations independently to generate a large number of candidate models (Kuhlman and Bradley, 2019; Rohl *et al*., 2004). IPTDFold is designed based on a population optimization framework, which can realize information sharing between conformations in the population. The initial population is generated by randomly fragment assembly. Then, the three sampling modules (A), (B), and (C) in **Figure 1** are designed to iteratively optimize the conformation in the population. Finally, the lowest-potential conformation is selected from the final population as the prediction model.

**Fig.1.**
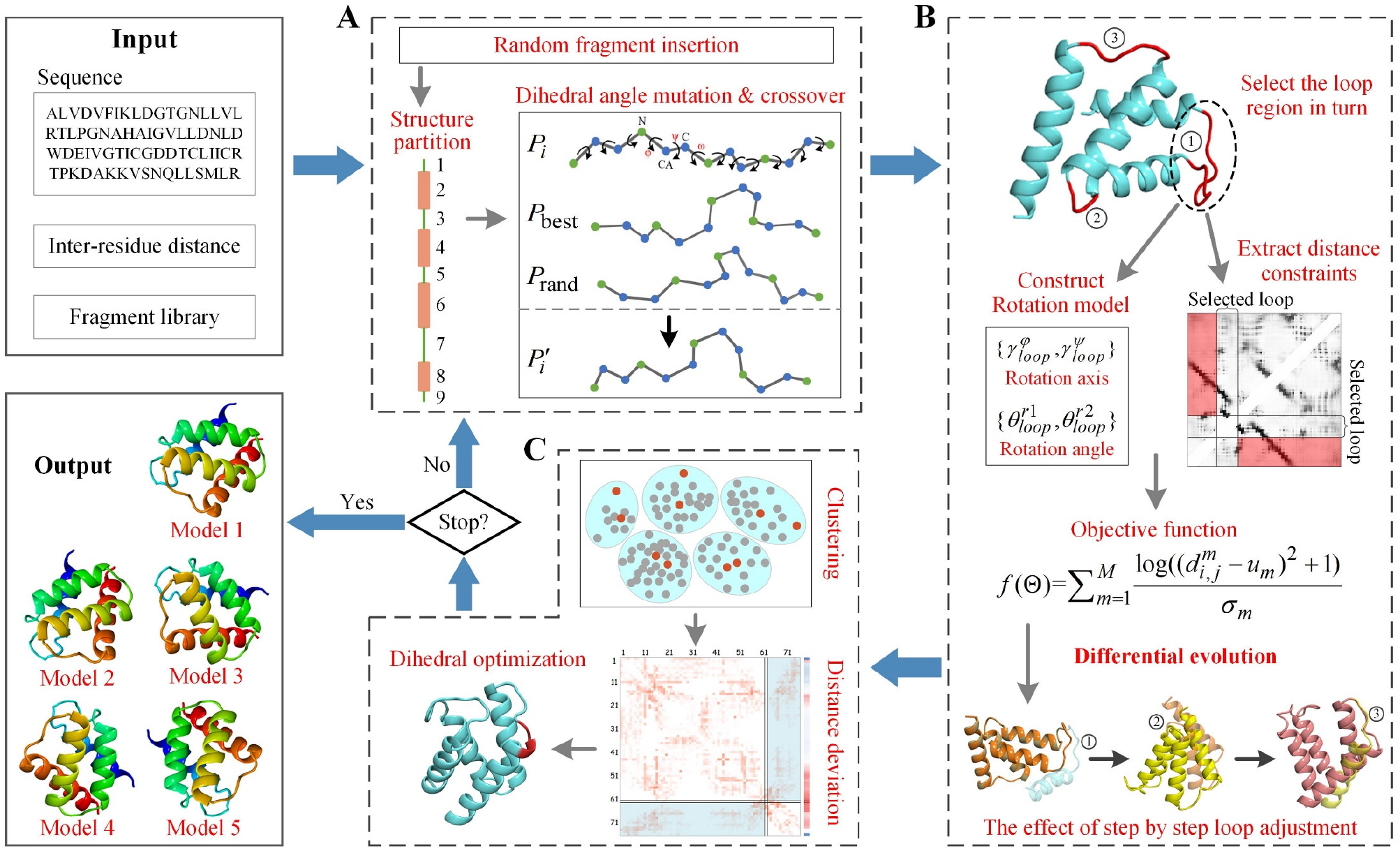
IPTDFold pipeline. (A) Partition sampling. For each conformation in the population, fragment insertion, structure partitioning, and local structure dihedral mutation and crossover are performed to explore the conformational space extensively. (B) Topology adjustment. For the loop region of the conformation, a local dihedral rotation model is first constructed. Then, the objective function is constructed by partial distance to guide the differential algorithm to find the optimal rotation angles, so as to adjust the spatial relationship of secondary structures. (C) Residue-level distance deviation optimization. The representative conformations are selected from the population by clustering. Then, the distance deviation of each residue is estimated according to the difference between the representative conformation and the predicted distance, which is used to perform probability-biased residue dihedral angle optimization.

### 2.1 Distance potential

The predicted inter-residue distance provides abundant spatial constraint information for protein folding. In this study, the inter-residue distance is constructed as a potential, which is defined as follows:

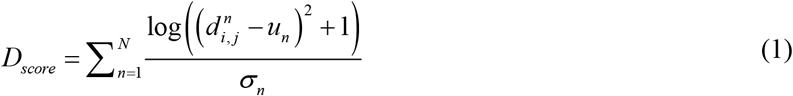

where *N* is the number of effective inter-residue distance; *i* and *j* are the residue indexes of the *n*-th distance; *d_i,j_* is the real distance between C_*β*_ (C_*α*_ for glycine) of residues *i* and *j* in the evaluated conformation; *u_n_* and *σ_n_* n are the mean and standard deviation obtained by Gaussian fitting of the *n*-th inter-residue distance distribution, respectively. *D_score_* and Rosetta score2 (Rohl *et al*., 2004) are combined with the same weight to form a composite potential function to guide folding.

### 2.2 Partition sampling

Although fragment assembly greatly reduces the conformational space, conformational sampling remains a challenging problem, and it easily causes insufficient or ineffective sampling in some regions (Kuhlman and Bradley, 2019). Crossover and mutation are the source of power for population evolution in nature. Crossover is the reorganization and inheritance of parental genes, while mutation will produce new genes. We introduced local structure crossover and mutation operators to increase sampling diversity. Fragment assembly is only the combinatorial optimization of the existing dihedral angles in the fragment library and it has difficulty crossing the energy barrier, while crossover and mutation can generate new dihedral angles and explore more potential conformations.

The flowchart of the partition sampling module is shown in Supplementary **Figure S1**. For the target conformation *P_i_*, random fragment insertion is first performed. Then, the DSSP algorithm (Kabsch and Jones, 1983) is used to calculate the secondary structure, which is used to divide the conformation into multiple local structures. Each local structure will undergo dihedral angle mutation and crossover operations in sequence. For each residue in a local structure, dihedral angle mutation and crossover are performed as following equation:

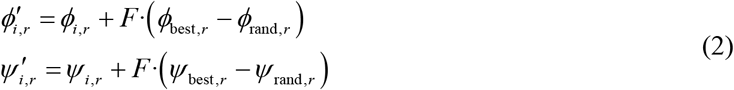

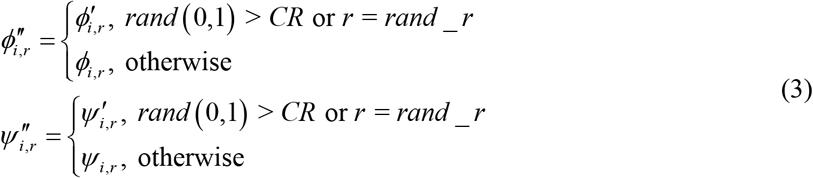

where *ϕ_i,r_* and *ψ_i,r_* are the *ϕ* and *ψ* dihedral angles of the *r*-th residue of *P_i_*; similarly, *ϕ*_best,*r*_ and *ψ*_best,*r*_ are the dihedral angles of the *r*-th residue of the lowest-potential conformation *P*_best_ in the population, *ϕ*_rand,*r*_ and *ψ*_rand,*r*_ are the dihedral angles of the *r*-th residue of the randomly selected conformation *P*_rand_ in the population; *F* is the scaling factor. 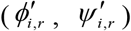 and 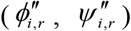 represent the dihedral angles obtained by mutation and crossover operations, respectively; *CR* is the crossover rate; and *rand_r* is the index of a random residue in the local structure. For each residue in the local structure, (*ϕ_i,r_*, *ψ_i,r_*) in the target conformation *P_i_* is replaced with 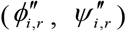 to generate a new conformation 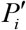. The composite potential function is used to calculate the potentials of *P_i_* and 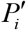, and the Metropolis criterion (Metropolis *et al*., 1953) is used to decide whether to replace *P_i_* with 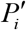. The above steps are applied to traverse all the local structures to complete the partition sampling.

### 2.3 Topology adjustment

The structure of the protein loop region connected with the regular *α*-helixes or *β*-sheets is flexible and irregular, and its small changes will produce dramatic effect on the entire topology (Liu *et al*., 2020). Many methods have been proposed to enhance the sampling of the loop Region (Spassovetal.,2008; Sotoetal.,2008; Arnautovaetal.,2011; Liang *et al*., 2014; Marks and Deane, 2019). The detailed inter-residue distance undoubtedly provides a powerful constraint for flexible loop sampling. According to the characteristics of the inter-residue distance, we designed a loop-specific dihedral angle optimization strategy to adjust the topology efficiently.

As shown in **Figure 1(B)**, for the target conformation, the secondary Structure is first calculated by DSSP (Kabsch and Jones,1983), and then the dihedral optimization is performed in turn for the loop regions connected with *α*-helix or *β*-sheet at both ends (such as the regions ①, ②, and ③ marked in red in **Figure1 (B)**). For the selected loop region, a loop-specific dihedral angle rotation model is constructed as shown in **Figure 2**. The set of dihedral angle rotation axes Γ of the loop region is defined as follows:

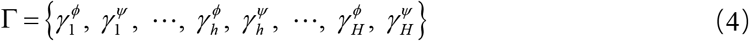

where 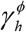 and 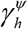 are the unit axes of the N-C_*α*_ and C_*α*_-C atomic bonds of the *h*-th residue in the selected loop region, corresponding to the dihedral angles *ϕ* and *ψ*, respectively. *H* is the number of residues in the selected loop region. The inter-residue distance with residues on both sides of the loop region are selected as the constraints. The coordinates of the C_*β*_ (C_*α*_ for glycine) atomics of the two residues of the m-th selected residue pair are expressed as 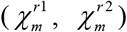.

**Fig.2.**
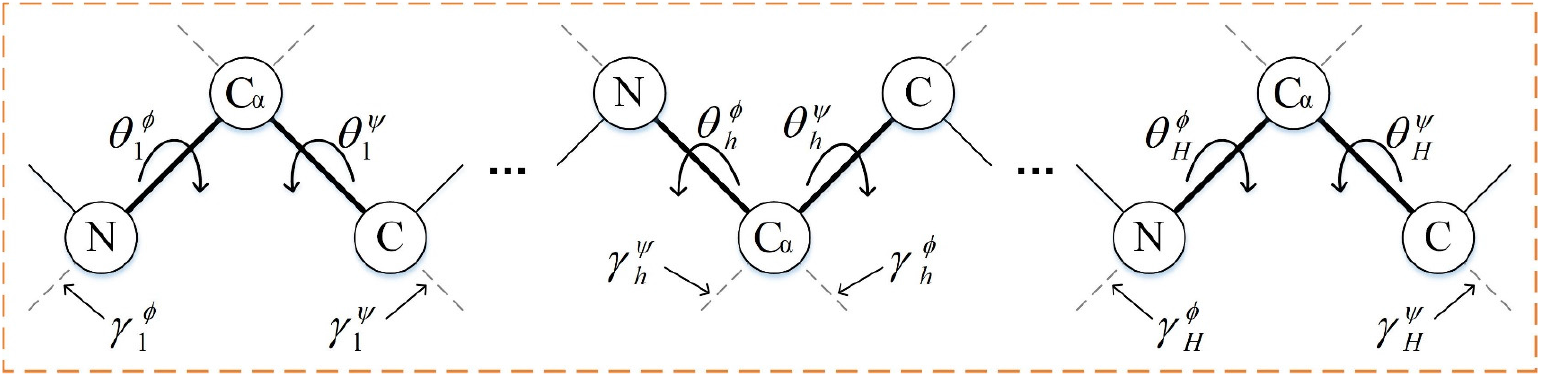
Schematic of the dihedral angle rotation model in the loop region. For each residue in the selected loop region, according to its φ and ψ dihedral angles, the rotation axis is extracted and the rotation angle is defined to construct a rotation model.

The aim of this section is to find a set of rotation angles 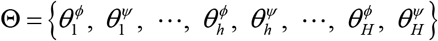 acting on the dihedral angle of the rotation axes set Γ, to generate a conformation that satisfies the distance constraint to the greatest extent. We use coordinate transformation and differential evolution algorithm (Hart *et al*., 1999) to generate such a set of rotation angles. For each rotation angle, a rotation matrix can be constructed. The detailed schematic of construction is shown in **Supplementary Figure S2**. The rotation matrix *T*(Θ) of the set of rotation angles Θ is defined as follows:

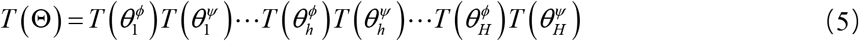

where 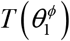 and 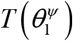 represent the rotation matrix formed by the rotation angles 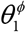 and 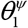 acting on the rotation axes 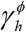 and 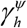, respectively. With the assumption that the residues on the left of the loop region remain fixed and the residues on the right are rotated, the coordinates of 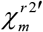 obtained by the rotation transformation of 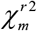 can be calculated:

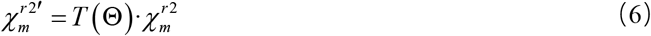

The objective function used in the differential evolution algorithm is defined as follows:

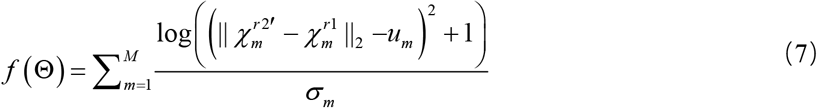

where *u_m_* and *σ_m_* are the mean and standard deviation of the *m*-th residue pair, and *M* is the number of residue pairs selected. The rotation angles Θ that best satisfies the distance constraint will be generated by the differential evolution algorithm (Zhang *et al*., 2016; Zhou *et al*., 2019), and the candidate conformations will be generated by adding the rotation angles to the corresponding dihedral angles of the loop region in the target conformation. If the composite potential of the candidate conformation is lower than that of the target conformation, the target conformation will be replaced by it.

### 2.4 Residue distance deviation optimization

The partition sampling module is a random sampling of the entire conformation, and the topology adjustment module is a loop-specific sampling. However, it is also important to know which residues have deviations that need to be adjusted and then correct them accordingly. In this section, the residue-level distance deviation is estimated through the difference of the conformational distance map and the predicted distance map, and then probability-biased residue dihedral angle optimization is performed to correct the residue distance deviation. First, DMscore, which is a structural similarity evaluation model we developed previously (Zhao *et al*., 2021), is used to calculate the similarity between conformations in the population. Then, the K-medoids algorithm (Kaufman and Rousseeuw, 2009) is used to cluster the conformations in the population, and the number of clusters is set to 5. The cluster center conformation and the lowest-potential conformation are selected from each cluster as the representative conformations. For each representative conformation, the distance deviation of each residue is calculated according to the distance map difference. The change in the dihedral angles of the residue will not cause a change in the internal structure on both sides, but it will affect the spatial position relationship between them. Therefore, the maximum gain that can be achieved by adjusting the residue can be estimated by calculating the distance deviation of residue pairs in which the two residues are respectively located on both sides of the current residue. As shown the distance difference map in Figure 3, for a given residue, only the residue pair in its upper-right or lower-left need to be considered. The distance deviation estimation for a given residue is quantified as follows:

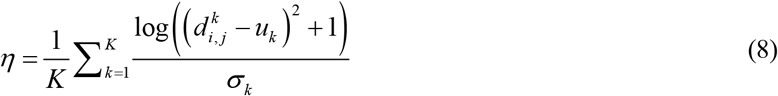

where *K* is the number of effective predicted inter-residue distance in the area to be considered; 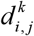 is the real distance between residue *i* and *j* of the *k*-th residue pair in the representative conformation; and *u_k_* and *σ_k_* are the mean and standard deviation of the distance distribution of the *k*-th residue pair, respectively. The probability is assigned to each residue according to the magnitude of residue distance deviation (the greater the deviation, the higher the probability), and then the residue is selected according to the probability for dihedral angle optimization. The residue dihedral angle optimization is achieved by using the partial distance constrained differential evolution algorithm similar to that of as the topology adjustment module. **Figure 3** shows an example of residue dihedral angle optimization guided by residue distance deviation. The difference between the distance map of representative conformation and the predicted distance map is first calculated, and then the distance deviation of each residue is calculated according to Equation 8 to obtain the residue distance deviation heat map, which is used to guide the residue dihedral angle optimization. It can be seen that the local structure of the conformation has been corrected to be closer to the experimental structure. Notably, the excellent genes of the representative conformations optimized in this module can be passed on to other conformations in the population through the mutation and crossover operator of the partition sampling module.

**Fig.3.**
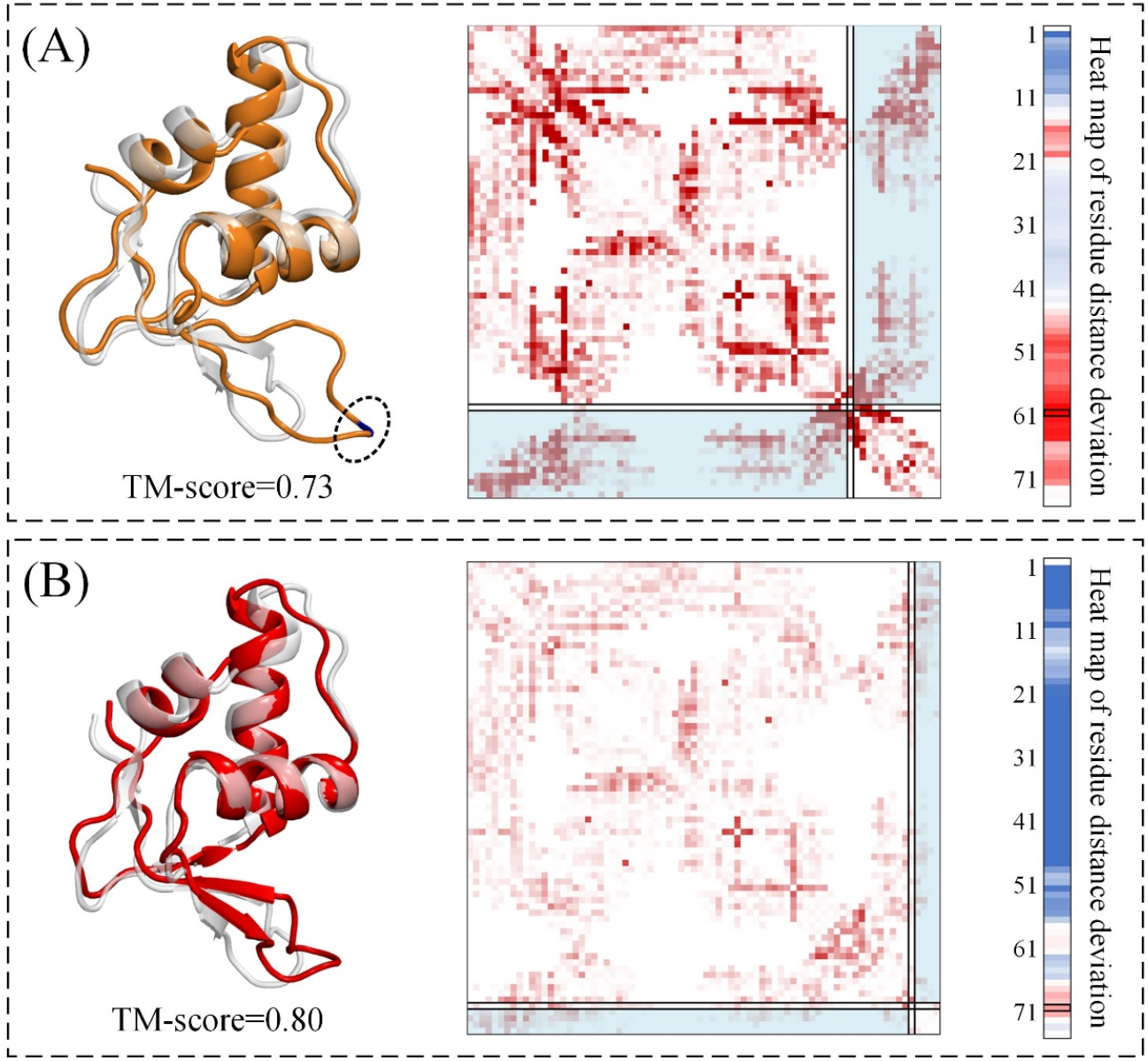
Example of residue distance deviation optimization. In (A), the left shows the alignment of the representative conformation structure (orange) and the experimental structure (light gray), the middle presents the difference between the distance map of representative conformation and the predicted distance map, and the right shows the residue distance deviation heat map (blue to red indicates the deviation from small to large). (B) is the corresponding ones after the dihedral angle optimization guided by the residue distance deviation.

## 3 Result and discussion

### 3.1 Dataset and evaluation metrics

IPTDFold is extensively tested on a non-redundant benchmark dataset of 462 proteins, 24 FM targets of CASP13, and 20 FM targets of CASP14. The benchmark test set was constructed from SCOPe2.07 (Fox *et al*.,2014) in several steps. First, CD-HIT (Fu *et al*., 2012) was used to cluster the SCOPe dataset with a 30% sequence identity cut-off, and the representative protein of each cluster constituted 11,198 non-redundant proteins. Then, a protein was discarded if its length is outside the range of 50 to 500 residues or contains multiple domains. Lastly, 462 proteins with a length ranging from 53 to 481 residues were randomly selected from the remaining 4,332 non-redundant proteins as the benchmark dataset. The detailed information of benchmark dataset and FM targets of CASP13 and CASP14 are shown in **Supplementary Tables S1, S4, and S5**, respectively. The root mean square deviation (RMSD) and TM-score (Zhang and Skolnick, 2004) are used to evaluate the predicted model’s accuracy.

### 3.2 Parameter settings

IPTDFold is a population-based structure prediction method, with the following parameters: population size *NP*=100, generation number *G*=100, dihedral mutation scaling factor *F*=0.5, dihedral crossover rate *CR*=0.5, and composite potential temperature scaling factor *KT*=2. The following are the parameters of the differential evolution algorithm for finding the dihedral rotation angles that satisfy the distance constraint: population size *NP′* =50, generation number *G′* =50, scaling factor *F′* =0.5, and crossover rate *CR′* =0.5. The number of iterations of residue dihedral angle optimization guided by the residue distance deviation is set to 500. The ratio of the iteration of partition sampling, topology adjustment, and residue-level distance deviation optimization is 10:1:2, that is, if partition sampling module runs for ten generations, topology adjustment module runs for one generation, and residue-level distance deviation optimization module runs for two generations. In this work, the fragment library was built by the Robetta fragment server with the “*Exclude Homologues*” option selected (http://robetta.bakerlab.org/fragmentsubmit.jsp), and the inter-residue distance was predicted by the trRosetta server with the “*Do not use templates*” option selected (http://yanglab.nankai.edu.cn/trRosetta/).

### 3.3 Comparison with Rosetta_D

IPTDFold is compared with the distance-assisted fragment assembly method Rosetta_D, which is a version of the Rosetta ClassicAbinitio protocol (Rohletal.,2004) that adds the inter-residue distance constraints. In the third and fourth stages of the ClassicAbinitio protocol, the same distance potential as IPTDFold is used to guide the fragment assembly together with the energy function. One thousand candidate models are generated by running 1,000 independent trajectories with the *increase_cycles* = 10, and then the candidate models are clustered by SPICKER (Zhang and Skolnick, 2004) and the centroid structure of the first cluster is selected as the final model. IPTDFold and Rosetta_D use the same fragment library and inter-residue distance potential. Thus, The comparison between them can fairly reflect the performance of the protein folding algorithms. The predicted results of IPTDFold and Rosetta_D on the benchmark dataset are summarized in **Table 1**, and the detailed results of each protein can be found in **Supplementary Table S2**.

**Table 1.** Results of IPTDFold and Rosetta_D on the benchmark dataset. #TM ≥ 0.5 and #TM ≥ 0.8 are the number of proteins with TM-score ≥ 0.5 and TM-score ≥ 0.8, respectively. The last column shows the result of the Wilcoxon signed-rank test calculated in accordance with TM-score.

| Method | RMSD | TM-score | #TM≥0.5 | #TM≥0.8 | *p*-value |
| --- | --- | --- | --- | --- | --- |
| IPTDFold | 6.12 | 0.715 | 430 | 125 | NA |
| Rosetta_D | 7.55 | 0.596 | 362 | 28 | 1.65E-70 |

The average RMSD and TM-score of IPTDFold are 6.12Å and 0.715, respectively. Compared with that of Rosetta_D, the average RMSD of IPTDFold is decreased by 18.9%, and the average TM-score is increased by 20.0%. IPTDFold correctly folds (i.e., TM-score≥0.5) 430 out of 462 proteins, accounting for 93.1% of the total, which is 14.7% more than Rosetta_D. IPTDFold predicted a model with TM-score≥0.8 on 125 proteins, which is 97 more than Rosetta_D. The Wilcoxon signed-rank test (Corder and Foreman, 2009) result in the last column of **Table 1** show that IPTDFold is significantly better than Rosetta_D. **Figure 4** further illustrates the comparison of the TM-score of IPTDFold and Rosetta_D on the benchmark dataset. IPTDFold achieves a higher TM-score on 423 test proteins, accounting for 91.6% of the total. Compared with the TM-score of Rosetta_D, the TM-score of IPTDFold increased by more than 0.1 on 234 proteins and by more than 0.2 on 94 proteins. **Figure 4(b)** shows the average TM-score of IPTDFold and Rosetta_D in different protein length bins, reflecting the relationship between prediction accuracy and protein length. IPTDFold obtains the highest average TM-score in all protein length bins. With the increase in the protein length, the average TM-score of IPTDFold shows a gentle upward trend and is relatively stable, whereas the average TM-score of Rosetta_D continues to decrease. Two possible reasons could explain why the prediction accuracy of Rosetta_D decreases with the increase in the protein length: (i) The fragment library discretizes the continuous dihedral space, which resulting in the possibility that the optimal dihedral angle cannot be matched during the folding process, and this influence will accumulate as the size of the protein increases. (ii) The protein conformational space expands geometrically as the length increases. It is difficult for random fragment assembly to find the near-native conformation in the broad and rugged conformational space. The possible reasons why the prediction accuracy of IPTDFold does not decrease with the increase of protein length are as follows: (i) Local dihedral angle mutation and crossover can break through the limitation of the fragment library, and explore more potential conformations through the information interaction of the conformations in the population. (ii) The partial distance-constrained dihedral angle optimization algorithm can efficiently generate the dihedral angle that satisfies the distance constraint to the greatest extent, and is also not restricted by fragment library. (iii) The loop-specific dihedral angle sampling and residue-level probability-biased dihedral angle optimization strategies can avoid insufficient or ineffective sampling to improve the sampling efficiency. The component analysis in Section 3.5 shows that the new moves and strategies we designed significantly improves the prediction accuracy of the algorithm.

**Fig.4.**
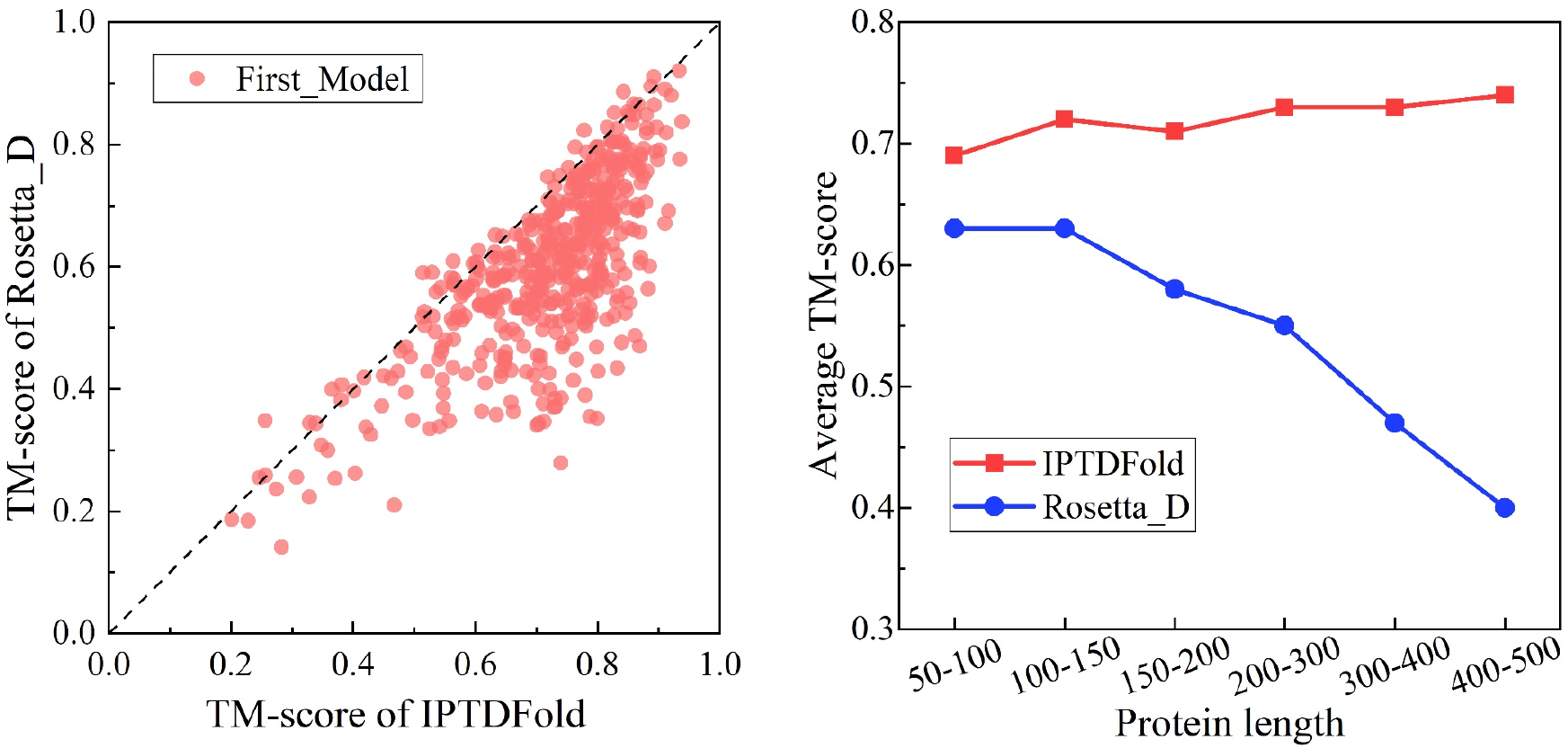
Comparison of TM-score between IPTDFold and Rosetta_D on the benchmark dataset. (a) Head-to-head comparison between TM-score of the first models predicted by IPTDFold and Rosetta_D. (b) Relationship between the average TM-score of the first model of IPTDFold and Rosetta_D and protein length.

### 3.4 Comparison with trRosetta

IPTDFold is also compared with the state-of-the-art gradient descent energy minimization method trRosetta on the benchmark dataset. In trRosetta, the predicted distance and orientation are used as restraints, together with the Rosetta energy function (Rohl *et al*., 2004), and coarse-grained models satisfying the restraints were generated by quasi-Newton minimization. The coarse-grained models were then subjected to Rosetta full-atom relaxation (*FastRelax*), including the distance and orientation restraints, to generate the lowest-energy full-atom model (Yang *et al*., 2020). IPTDFold did not use orientation and did not perform full-atomic refinement. For a fair comparison, we designed two sets of comparative experiments. One is a comparison between IPTDFold_relax1 (IPTDFold plus *FastRelax* with distance constraints) and trRosetta* (a version of trRosetta that excludes orientation restraints), and both of them use exactly the same *FastRelax* (Chaudhury *et al*., 2010). The other is a comparison of IPTDFold_relax2 (IPTDFold plus *FastRelax* with distance and orientation restraints, same as trRosetta) with the full version of trRosetta. The results of trRosetta* and trRosetta are obtained by running the official source code five times to generate five models, and the lowest-energy model is selected as the final model. The predicted results of the four methods on the benchmark dataset are **summarized in Table 2**, and the detailed results of each protein can be found in **Supplementary Table S2**. The comparison of TM-score of the first model is visually presented in **Figure 5**.

**Fig. 5.**
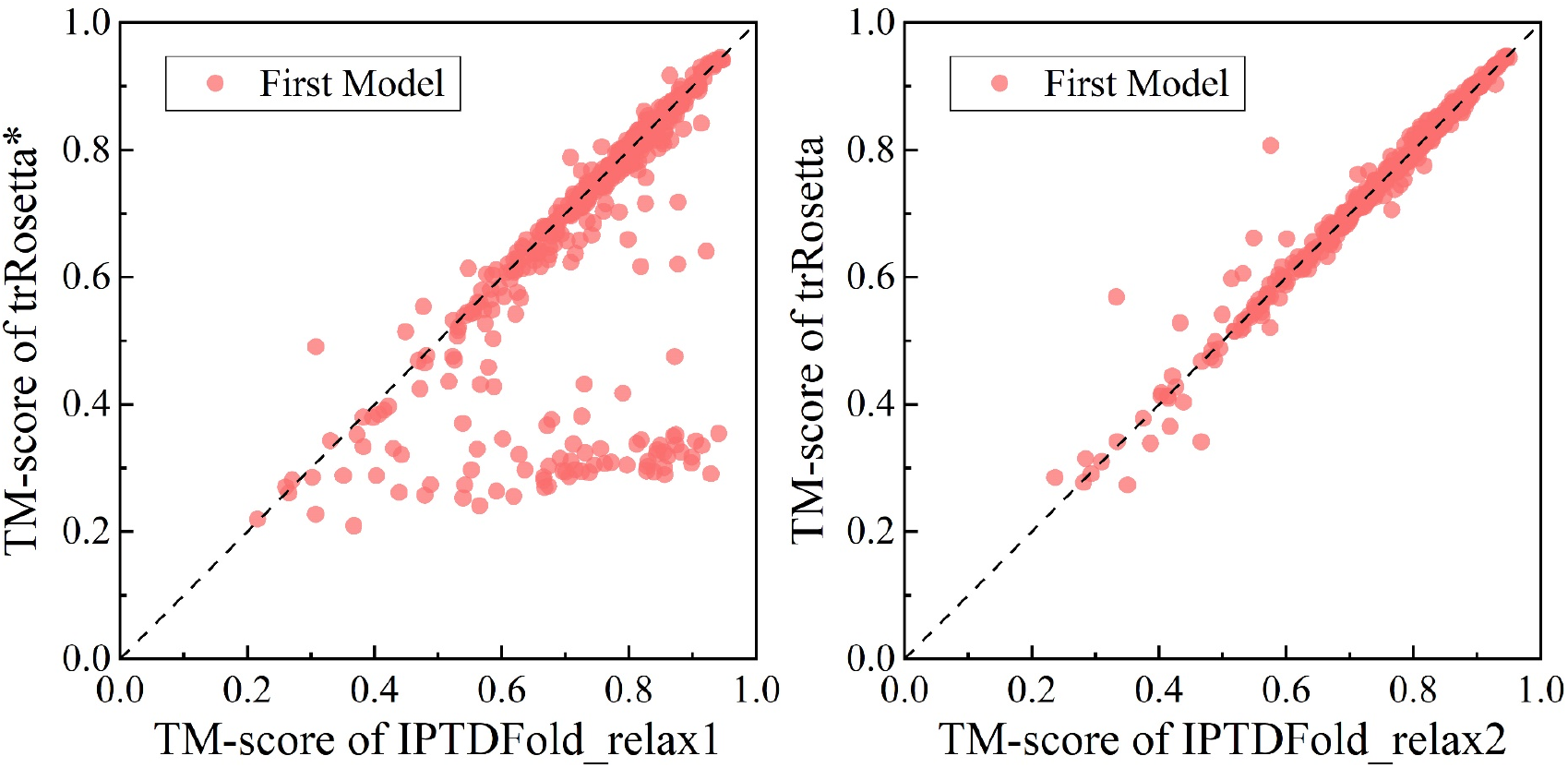
Comparison of TM-score of the first models on the benchmark dataset. (a) Head-to-head comparisons of TM-score between IPTDFold_relax1and trRosetta*. (b)Head-to-head comparisons of TM-score between IPTDFold_relax2 and trRosetta.

**Table 2.**
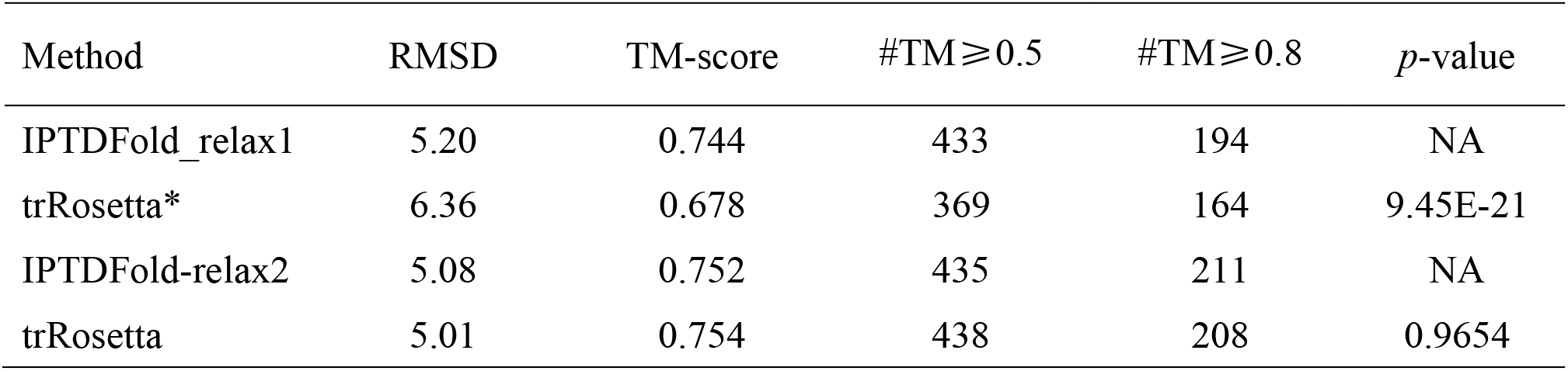
Results of IPTDFold_relax1, trRosetta*, IPTDFold_relax2, and trRosetta on the benchmark dataset. (*Note*:trRosetta* is a version of trRosetta that excludes orientation restraints. IPTDFold_relax1 is a version of IPTDFold with the same *FastRelax* protocol as trRosetta*. IPTDFold_relax2 is a version of IPTDFold with the same FastRelax protocol as trRosetta.)

| Method | RMSD | TM-score | #TM≥0.5 | #TM≥0.8 | *p*-value |
| --- | --- | --- | --- | --- | --- |
| IPTDFold_relax1 | 5.20 | 0.744 | 433 | 194 | NA |
| trRosetta* | 6.36 | 0.678 | 369 | 164 | 9.45E-21 |
| IPTDFold-relax2 | 5.08 | 0.752 | 435 | 211 | NA |
| trRosetta | 5.01 | 0.754 | 438 | 208 | 0.9654 |

The average RMSD and TM-score of IPTDFold_relax1 are 5.20Å and 0.744, respectively. Compared with that of trRosetta*, the average RMSD of IPTDFold_relax1 is decreased by 18.2%, and the average TM-score is increased by 9.73%. IPTDFold_relax1 correctly folds 433 test proteins, which is 64 more than that of trRosetta*. IPTDFold_relax1 achieved a lower RMSD on 272 proteins and higher TM-score on 312 proteins, accounting for 58.9% and 67.5% of the total, respectively. The Wilcoxon signed-rank test result shows that IPTDFold_relax1 is significantly superior to trRosetta*. **Figure 5(a)** shows that the TM-score of some protein models predicted by trRosetta* is abnormally low, and many of them are mirror structures. Interestingly, IPTDFold_relax1 (and IPTDFold) does not appear to generate abnormal mirror models, which may be because IPTDFold is potential-guided simulation folding. During the folding process, the quality of each folding step can be evaluated to ensure that the protein is folded in a reasonable direction. The average RMSD and TM-score of IPTDFold_relax2 are 5.08Å and 0.752, respectively. Compared with that of trRosetta, the average RMSD of IPTDFold_relax2 is 0.14% higher and the average TM-score is 0.27% lower. Compared with trRosetta, IPTDFold_relax2 has three fewer models with TM-score≥0.5, but has three more models with TM-score≥0.8. The Wilcoxon signed-rank test result shows no significant difference between IPTDFold_relax2 and trRosetta. It should be noted that the orientation constraint is added only to the *FastRelax* of IPTDFold_relax2, and is not used in the core algorithm of IPTDFold.

### 3.5 Component analysis

To examine the effects of the three sampling modules of IPTDFold, we set up two comparative versions IPTDFold-A and IPTDFold-AB for component analysis. IPTDFold-A means that only the partition sampling module is used, and IPTDFold-AB means that the partition sampling module and topology adjustment module are used. For a fair comparison, we set the population generation numbers of IPTDFold-A and IPTDFold-AB to 1.5 and 1.25 times those of IPTDFold, respectively, so that their computational costs are equal. The predicted results of IPTDFold, IPTDFold-A, and IPTDFold-AB on the benchmark test set are summarized in **Table 3**, and the detailed results on each protein can be found in **Supplementary Table S3**. **Figure 6** intuitively reflects the comparison of the TM-score of the first model.

**Table 3.**
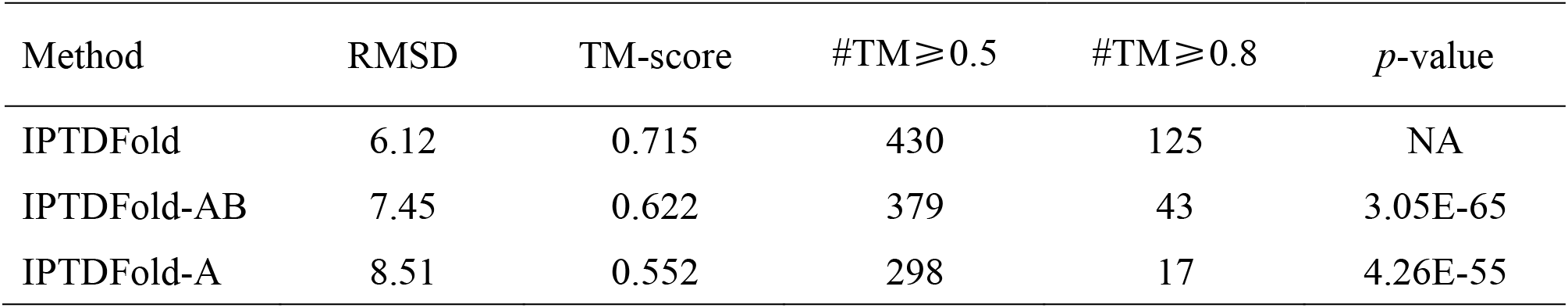
Results of IPTDFold, IPTDFold-AB, and IPTDFold-A on the benchmark dataset.

| Method | RMSD | TM-score | #TM≥0.5 | #TM≥0.8 | *p*-value |
| --- | --- | --- | --- | --- | --- |
| IPTDFold | 6.12 | 0.715 | 430 | 125 | NA |
| IPTDFold-AB | 7.45 | 0.622 | 379 | 43 | 3.05E-65 |
| IPTDFold-A | 8.51 | 0.552 | 298 | 17 | 4.26E-55 |

**Fig.6.**
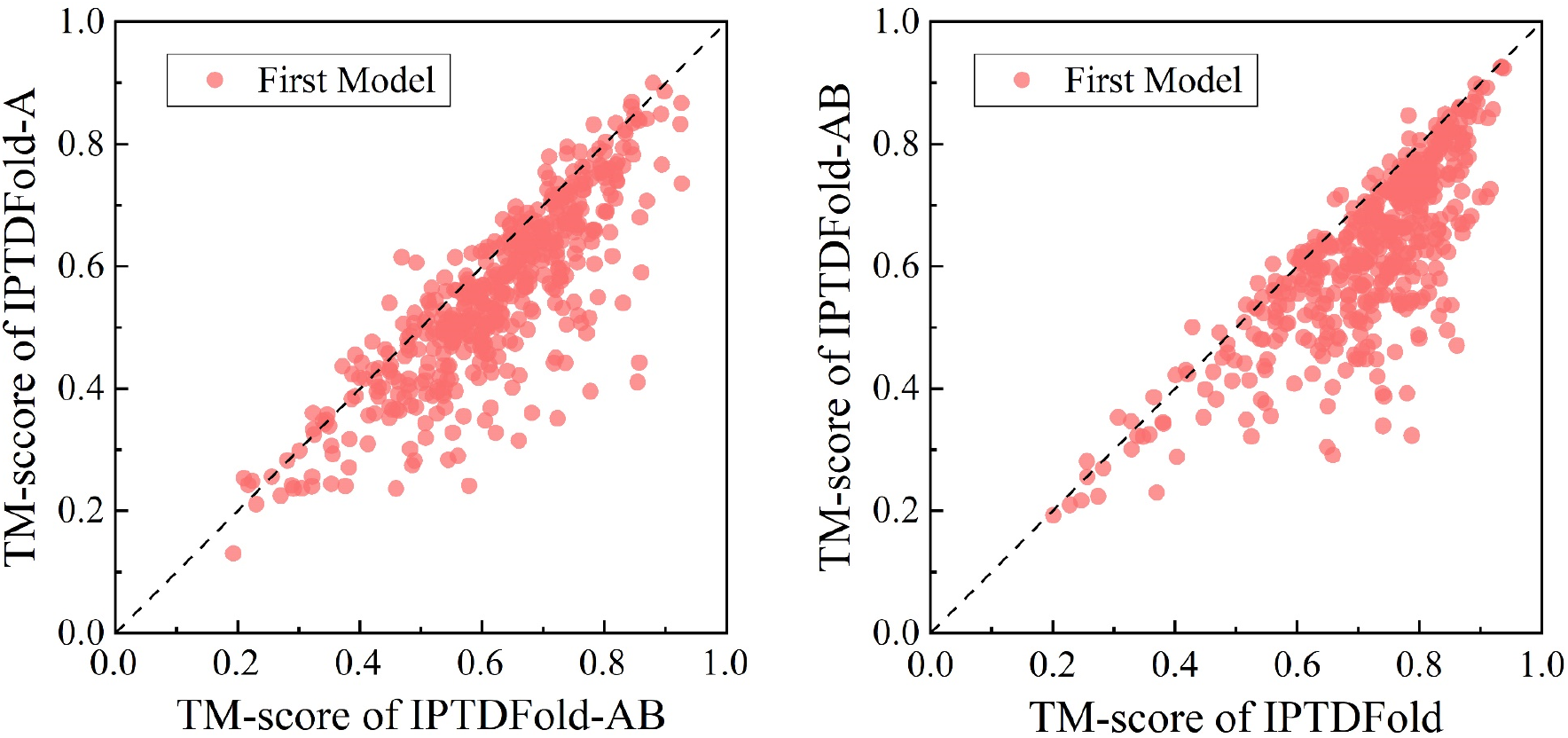
Comparison of TM-score between different IPTDFold versions on the benchmark dataset. (a) Head-to-head comparisons of TM-score between IPTDFold-AB and IPTDFold-A. (b) Head-to-head comparisons of TM-score between IPTDFold and IPTDFold-AB.

The average RMSD and TM-score of the first model generated by IPTDFold-A are 8.51Å and 0.552, respectively. The average RMSD of IPTDFold-AB is decreased by 12.5%, and the average TM-score is increased by 12.7% when the topology adjustment module is added. The average RMSD of IPTDFold is further reduced by 17.9%, and the average TM-score is further increased by 15.0% when the residue-level distance deviation module is added. IPTDFold-A correctly folds 298 proteins, and generates the first model with TM-score≥0.8 on 17proteins. On this basis, the number of correctly folded proteins increased by 81, and the number of proteins with the first model’s TM-score≥0.8 increased by 26 when the topology adjustment module is added. IPTDFold further increased the number of proteins to 430, and increased the number of proteins with first model’s TM-score≥0.8 to 125. IPTDFold-AB achieves lower RMSD and higher TM-score than IPTDFold-A on 329 and 381 proteins, respectively. The TM-score of IPTDFold-AB increased by more than 0.1 on 132 proteins, and increased by more than 0.2 on 33 proteins. Furthermore, IPTDFold achieves lower RMSD and higher TM-score than IPTDFold-AB on 361 and 424 proteins, respectively. The TM-score of IPTDFold increased by more than 0.1 on 174 proteins, and increased by more than 0.2 on 57 proteins. The Wilcoxon signed-rank test results show that the performance improvement of each step is significant. **Figure 7** reflects the contribution of different components of IPTDFold (including *FastRelax*), and the prediction accuracy of different methods. The clustered histogram of the TM-score of the first model predicted by IPTDFold, Rosetta_D, IPTDFold_relax1, trRosetta*, IPTDFold_relax2, and trRosetta is shown in **Supplementary Figure S3**.

**Fig.7.**
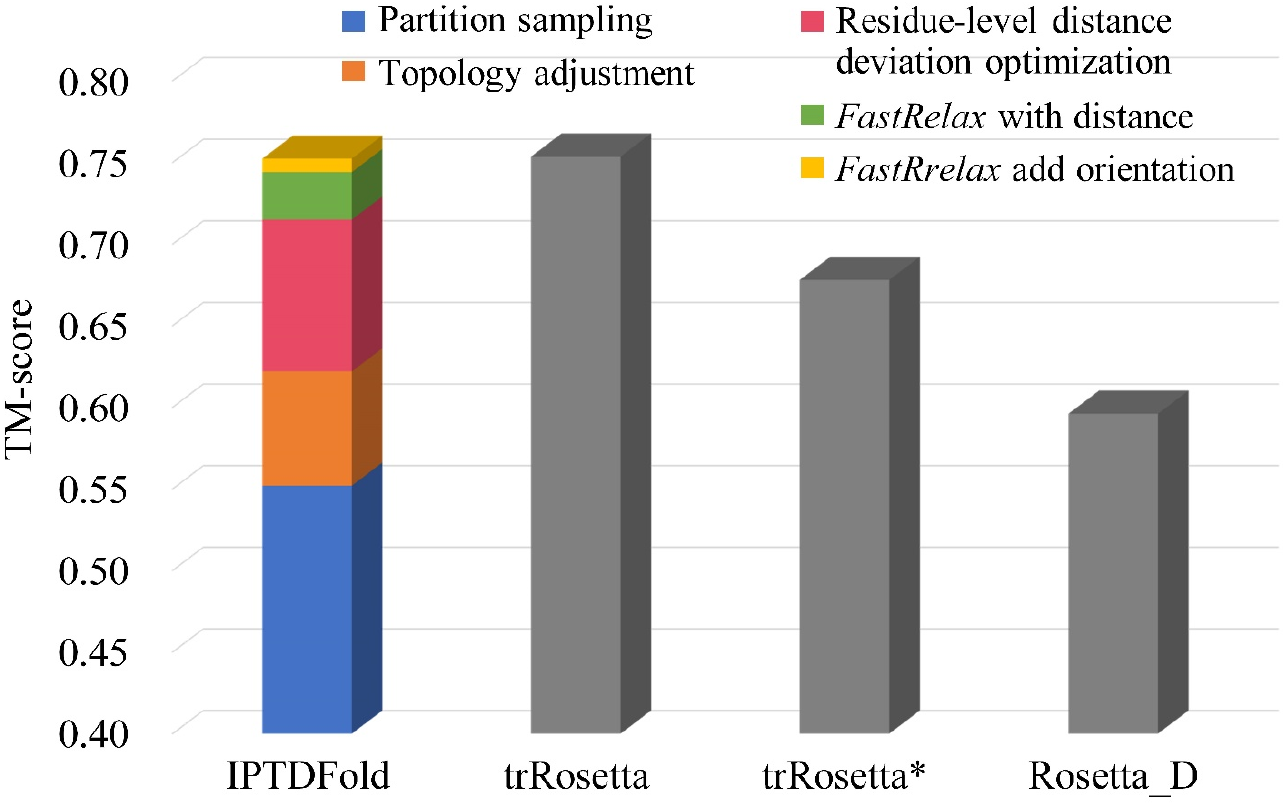
Average TM-score of all methods on the 462 benchmark test proteins. The colored stacked bar indicates the contributions of different components to our method.

### 3.6 Results on CASP targets

We also tested IPTDFold on 24 FM targets of CASP13 and 20 FM targets of CASP14. On the 24 targets of CASP13, IPTDFold and IPTDFold_relax1 was compared with four state-of-the-art methods of server groups in CASP13, i.e., QUARK (Zheng *et al*., 2019), RaptorX-Contact (Xu and Wang, 2019), BAKER-ROSETTASERVER (Park *et al*., 2019), and MULTICOM-CLUSTER (Houetal.,2019). On the 20 targets of CASP14, IPTDFold and IPTDFold_relax1 was compared with four state-of-the-art methods of server groups in CASP14, i.e., QUARK (Zhang *et al*., 2020), RaptorX (Xu *et al*., 2020), BAKER-ROSETTASERVER (Anishchenko *et al*., 2020), and MULTICOM-CLUSTER (Liu *et al*., 2020). The results of the compared methods were obtained from the CASP official website (http://predictioncenter.org). **Figure 8** shows the TM-score of the first model predicted by each method on each target, and the detailed results are listed in **Supplementary Tables S4 and S5**. For the 24 targets of CASP13, the average TM-score of IPTDFold is 0.55, which is 10.0% higher than that of QUARK (0.50), 12.2% higher than that of RaptorX-Contact (0.49), 30.9% higher than that of BAKER-ROSETTASERVER (0.42), and 48.6% higher than that of MULTICOM_CLUSTER (0.37). The average TM-score of IPTDFold_relax1 is 0.59, which is 18.0% higher than that of QUARK, 20.4% higher than that of RaptorX-Contact, 40.5% higher than that of BAKER-ROSETTASERVER, and 59.5% higher than that of MULTICOM_CLUSTER. IPTDFold and IPTDFold_relax1 correctly fold 15 and 20 of the 24 targets, respectively. For the 20 FM targets of CASP14, the average TM-score of IPTDFold is 0.41, which is the same as that of RaptorX (0.41) and BAKER-ROSETTASERVER (0.41), 24.1% lower tha that of QUARK (0.54), and 10.7% lower than that of MULTICOM_CLUSTER (0.46). The average TM-score of IPTDFold_relax1 is 0.47, which is 14.6% higher than that of RaptorX and BAKER-ROSETTASERVER, 2.2% higher than that of MULTICOM_CLUSTER, and 13.0% lower than that of QUARK. IPTDFold and IPTDFold_relax1 correctly fold 6 and 10 of the 20 targets, respectively. **Figure 9** shows the superimpositions of the target model predicted by IPTDFold and the experimental structure for four targets.

**Fig.8.**
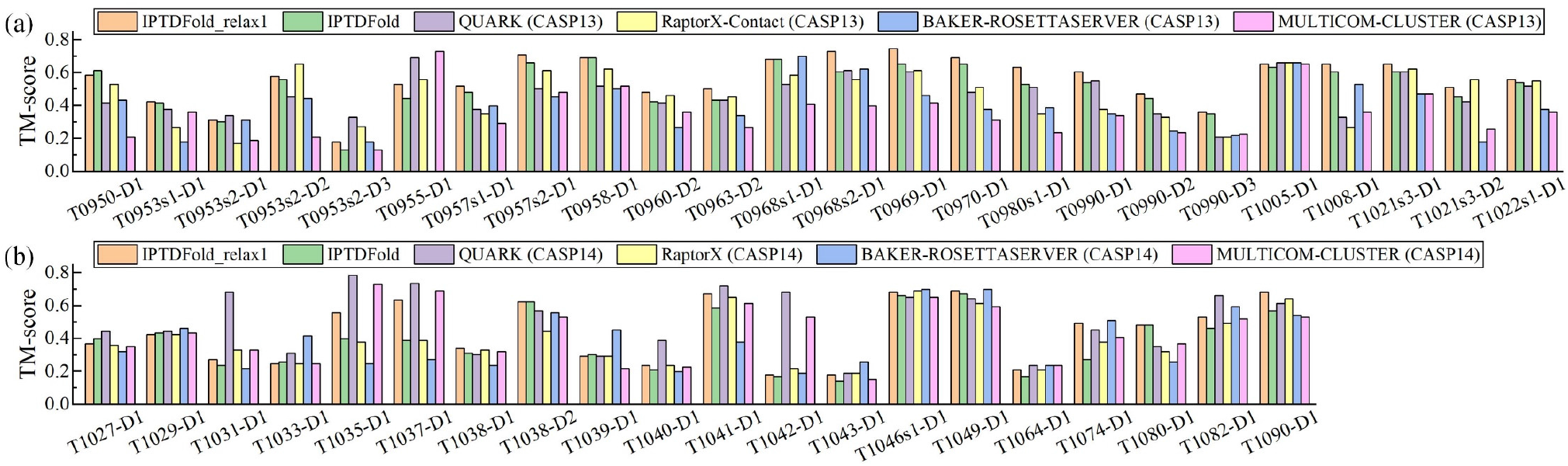
TM-score of the first model predicted by IPTDFold_relax1 (IPTDFold plus FastRelax with distance constraints) IPTDFold, QUARK, RaptorX (RaptorX-Contact for CASP13), BAKER-ROSETTASERVER, and MULTICOM-CLUSTER on the 44 FM targets. (a) Results on the 24 FM targets of CASP13. (b) Results on the 20 FM targets of CASP14.

**Fig.9.**
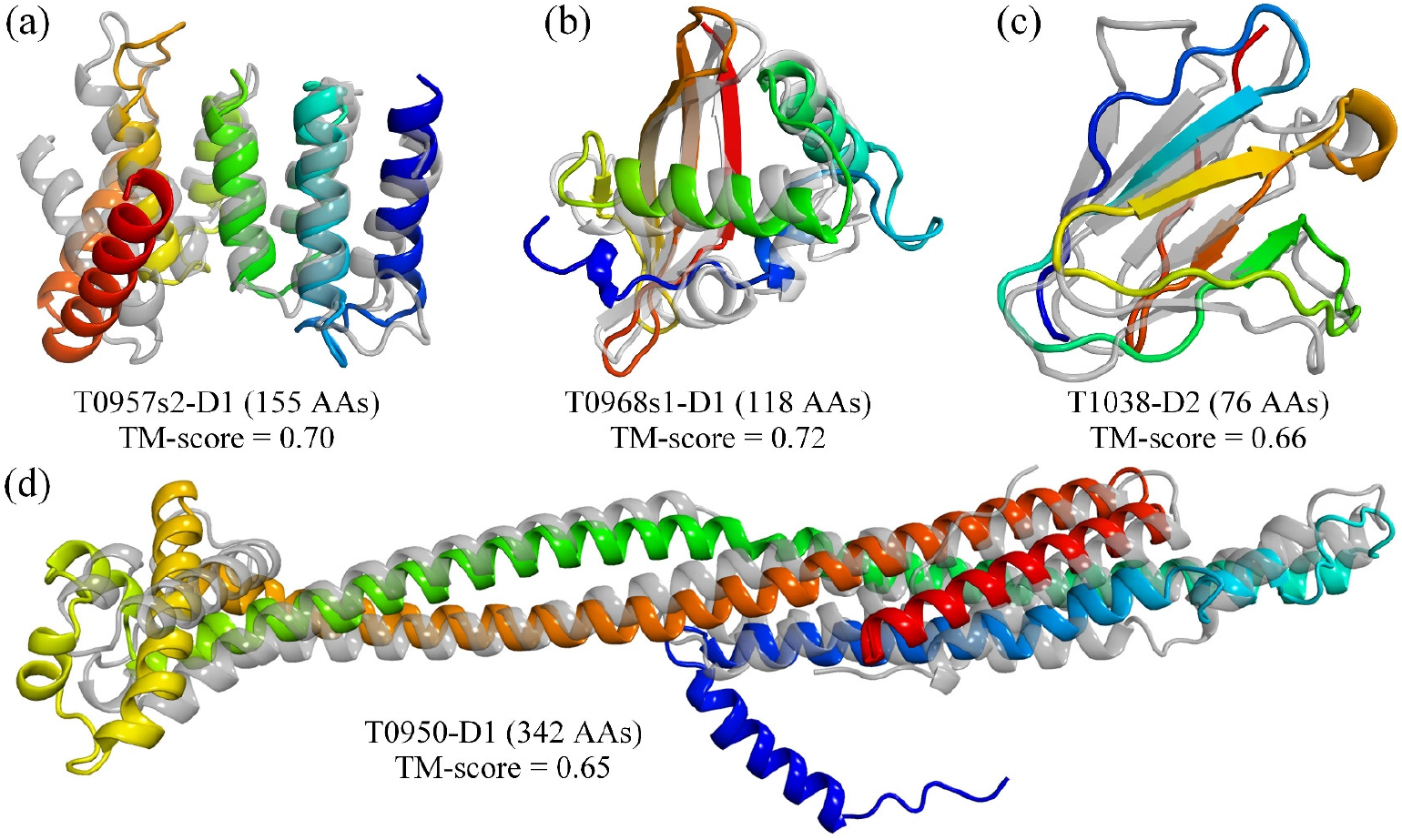
Superimposition between the first model (rainbow) by IPTDFold and the experimental structure (gray) for four FM targets (T0957s2-D1, T0968s1-D1, T1038-D2, and T0950-D1) of CASP13 and CASP14.

## 4 Conclusion

We developed a closed-loop iterative partition sampling, topology adjustment and residue-level distance deviation optimization algorithm called IPTDFold for de novo protein structure prediction. In the partition sampling module, random fragment insertion and local dihedral angle crossover and mutation operators are used to extensively explore the conformational space. In the topology adjustment module, a loop-specific dihedral rotation model is constructed, and the partial inter-residue distances are extracted to construct the rotation angles objective function. The differential evolution algorithm is used to find the rotation angles that satisfy the distance constraint as much as possible to realize spatial position adjustment between secondary structures. In the residue-level distance deviation optimization module, the residue distance deviation is estimated through the difference of the inter-residue distance map, and then the probability-biased residue dihedral angle optimization is performed according to the degree of residue distance deviation. The protein structure model is predicted by iterating the above three modules. IPTDFold is extensively tested on 462 benchmark test proteins, 24 FM targets of CASP13, and 20 FM targets of CASP14. Experimental results show that the prediction accuracy of IPTDFold is significantly better than that of the distance-assisted fragment assembly method Rosetta_D. When using the same *FastRelax* protocol, the prediction accuracy of IPTDFold is significantly superior to that of trRosetta without orientation constraints, and is equivalent to that of the full version of trRosetta. IPTDFold correctly folds 430 out of 462 benchmark test proteins and 21 out of FM targets of CASP13 and CASP14. Notably, the prediction accuracy of IPTDFold does not decrease as the increase in protein size. As the conformation space expands geometrically with the increase in the protein length, a challenging problem for the fragment assembly-based conformation sampling method is to ensure prediction accuracy for larger proteins. IPTDFold is developed on the basis of the traditional fragment assembly conformation sampling mechanism, and it significantly improves the accuracy of structure prediction. With the rapid development of model quality evaluation technology, if model evaluation can be integrated into the folding procedure to build a feedback mechanism, it will help to further improve the protein structure prediction accuracy.

## Supporting information

Suppport Information

## Funding

This work has been supported by the National Nature Science Foundation of China (No. 61773346) and the Key Project of Zhejiang Provincial Natural Science Foundation of China (No. LZ20F030002).

## Notes

### Competing Interest Statement

The authors have declared no competing interest.

