## Supplementary material for "A *de novo* protein structure prediction by iterative partition sampling, topology adjustment, and residue-level distance deviation optimization": Suppport Information

### Supplementary Information

#### Table of Content

**Supplementary Figures**

**Figure S1.** Flowchart of partition sampling.

**Figure S2.** Schematic construction of rotation matrix based on dihedral angles.

**Figure S3.** Clustered histogram of the TM-score of the first model predicted by IddFold, Rosetta D, IddFold relax, trRosetta D, IddFold relax2, and trRosetta on the benchmark dataset..

**Supplementary Tables**

**Table S1.** Detailed information of 462 proteins in the benchmark dataset.

**Table S2.** Results of the first model predicted by IddFold, Rosetta_D, IddFold_relax, trRosetta_D, IddFold_relax2 and trRosetta on the benchmark dataset.

**Table S3.** Results of the first model predicted by IddFold-AB and IddFold-A on the benchmark dataset.

**Table S4.** TM-score of the first model predicted by IddFold_relax1, IddFold and four server groups on the targets of CASP13.

**Table S5.** TM-score of the first model predicted by IddFold_relax1, IddFold and four server groups on the targets of CASP14.

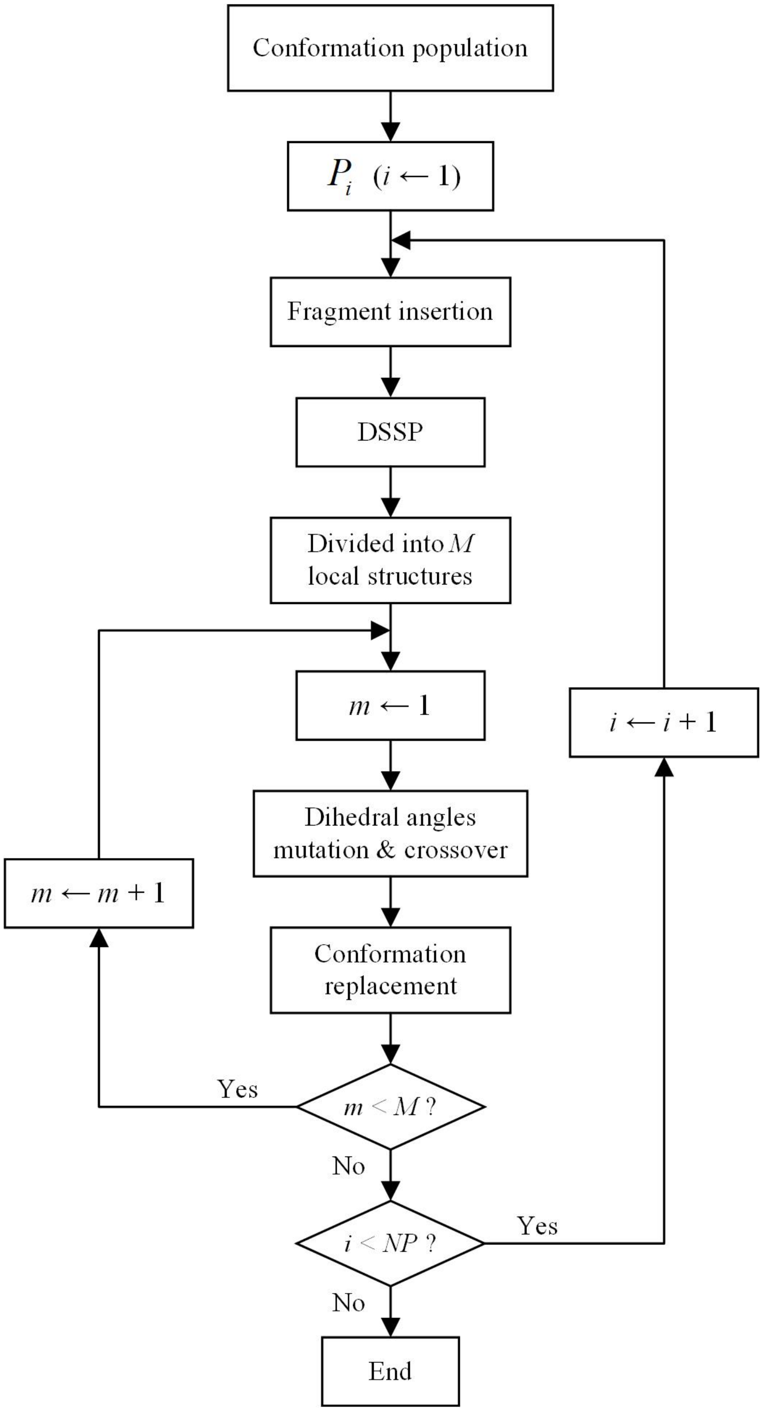


Figure S1: Flowchart of Partition sampling. $NP$ is the population size, and $M$ is number of local structures divided according to secondary structure.

Figure S2: Schematic construction of rotation matrix based on dihedral angles

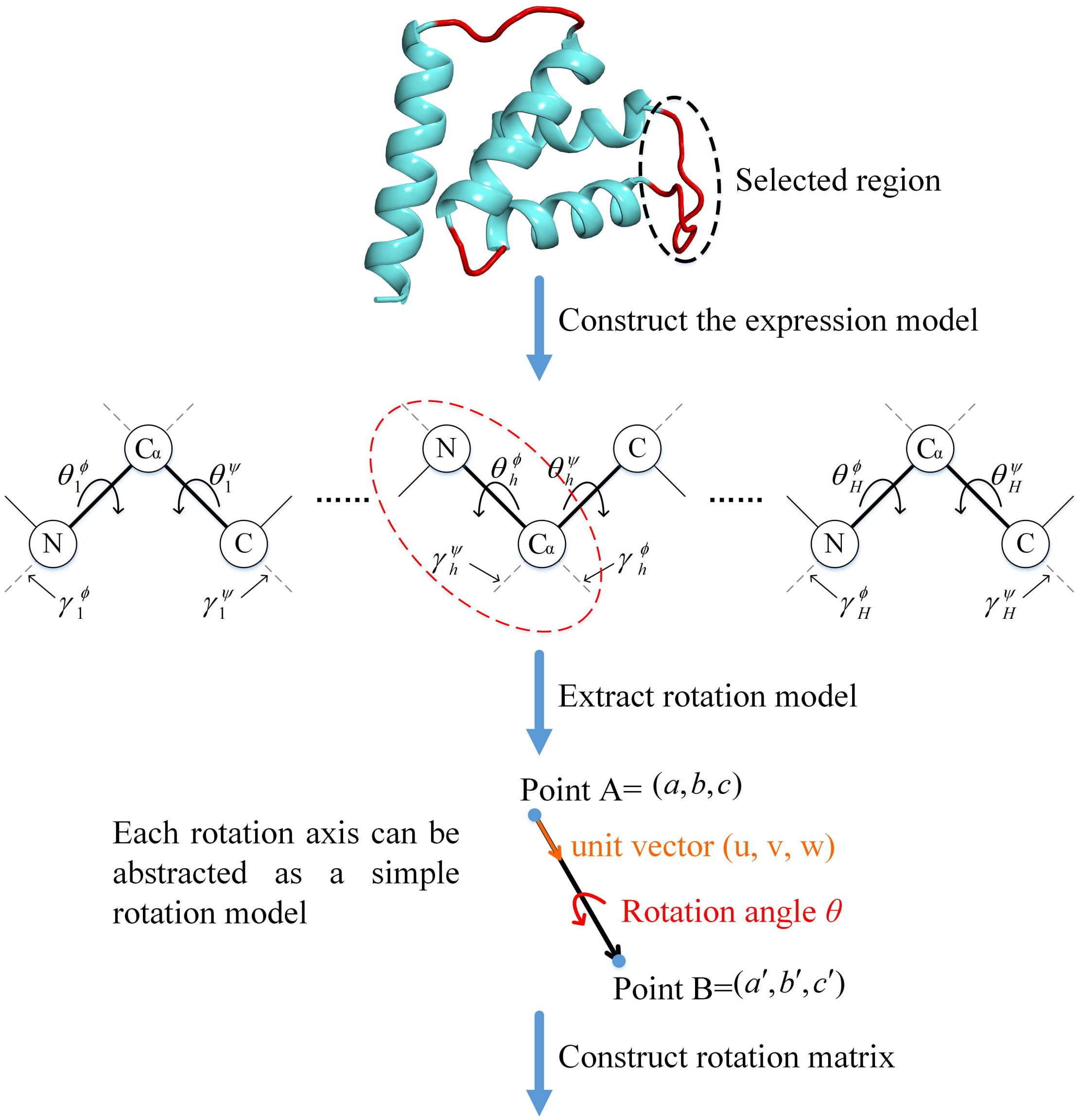


$$T(\theta)=\begin{bmatrix} u^2+(v^2+w^2)\cdot\cos\theta & uv\cdot(1-\cos\theta)-w\cdot\sin\theta & uw\cdot(1-\cos\theta)+v\cdot\sin\theta & t_{1,4} \\ uv\cdot(1-\cos\theta)+w\cdot\sin\theta & v^2+(u^2+w^2)\cdot\cos\theta & vw\cdot(1-\cos\theta)-u\cdot\sin\theta & t_{2,4} \\ uw\cdot(1-\cos\theta)-v\cdot\sin\theta & vw\cdot(1-\cos\theta)+u\cdot\sin\theta & w^2+(u^2+v^2)\cdot\cos\theta & t_{3,4} \\ 0 & 0 & 0 & 1 \end{bmatrix}$$

where

$$t_{1,4}=(a\cdot(v^2+w^2)-u\cdot(bv+cw))\cdot(1-\cos\theta)+(bw-cv)\cdot\sin\theta$$

$$t_{2,4}=(b\cdot(u^2+w^2)-v\cdot(ax+cw))\cdot(1-\cos\theta)+(cu-aw)\cdot\sin\theta$$

$$t_{3,4}=(c\cdot(u^2+v^2)-w\cdot(av+bv))\cdot(1-\cos\theta)+(av-bu)\cdot\sin\theta$$

All the rotation axes can form a joint rotation matrix:

$$T(\Theta)=T(\theta_1^{\phi})T(\theta_1^{\psi})\cdots T(\theta_h^{\phi})T(\theta_h^{\psi})\cdots T(\theta_H^{\phi})T(\theta_H^{\psi})$$

Figure S3: Clustered histogram of TM-score of the first model predicted by IddFold, Rosetta_D, IddFold_relax, trRosetta_D, IddFold_relax2, and trRosetta on the benchmark dataset.

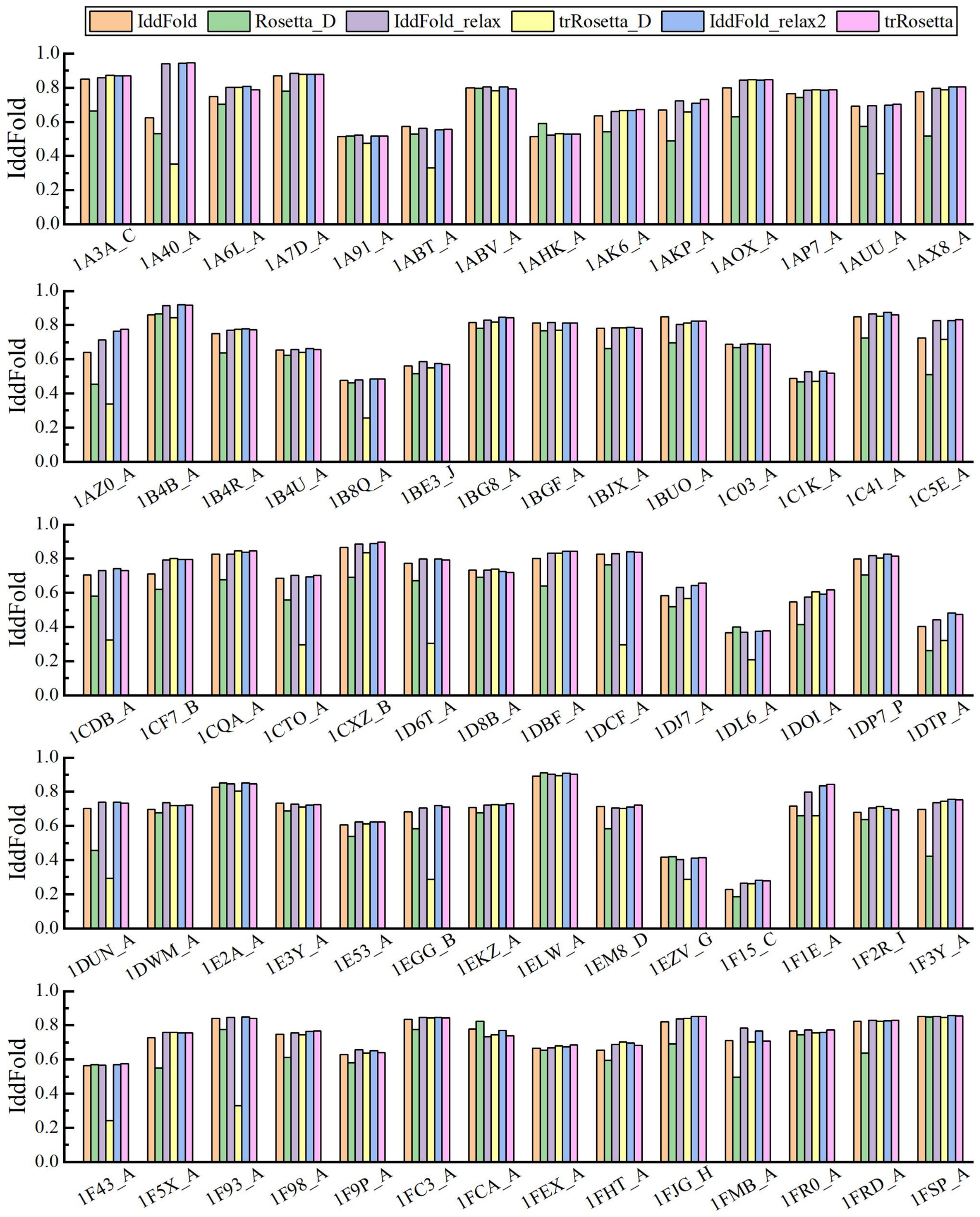

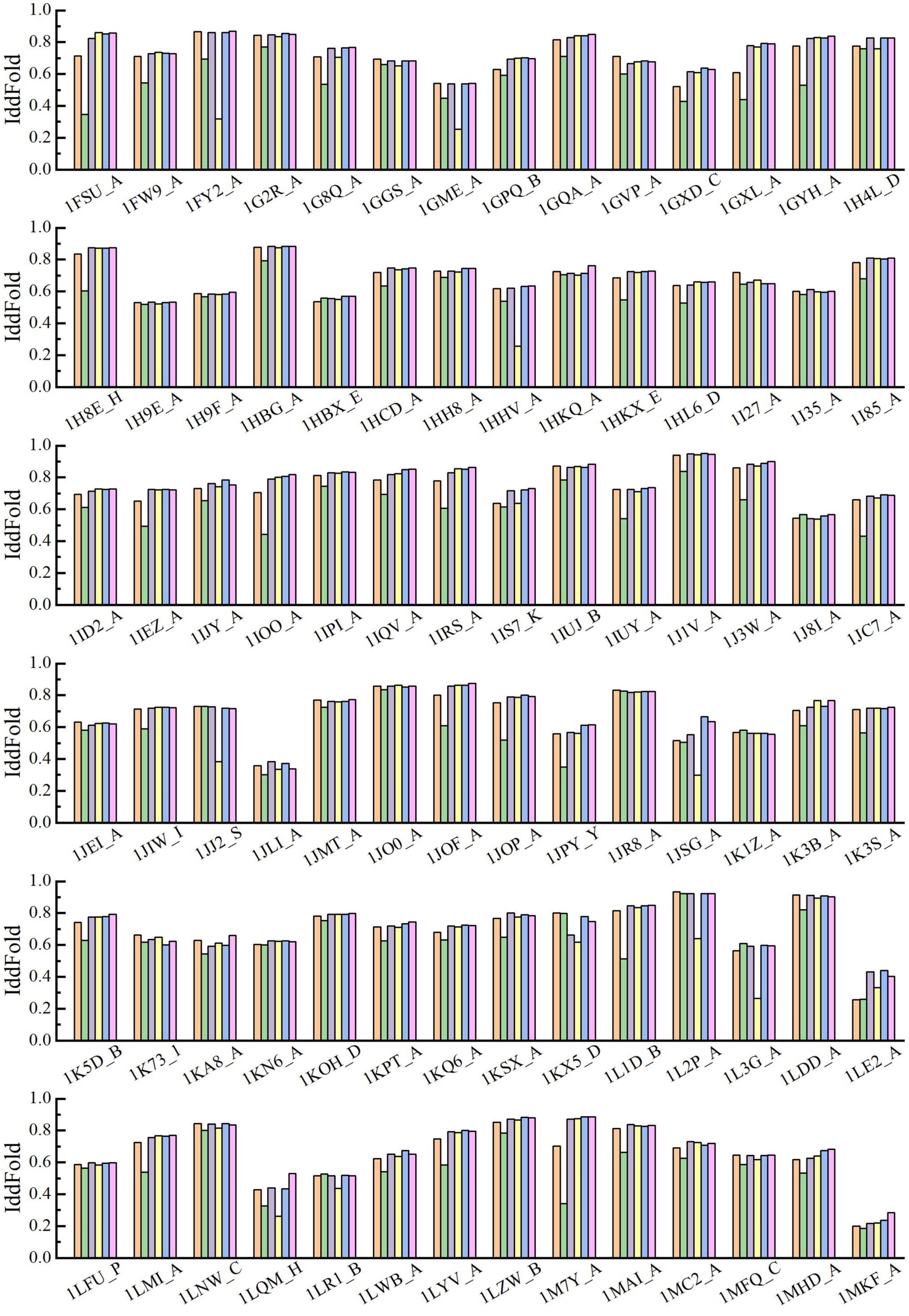

IddFold
1.0
0.8
0.6
0.4
0.2
0.0
1FSU_A
1FW9_A
1FY2_A
1G2R_A
1G8Q_A
1GGS_A
1GME_A
1GPQ_B
1GQA_A
1GVP_A
1GXD_C
1GXL_A
1GYH_A
1H4L_D
1H8E_H
1H9E_A
1H9F_A
1HBG_A
1HBX_E
1HCD_A
1HH8_A
1HHV_A
1HKQ_A
1HKX_E
1HL6_D
1I27_A
1I35_A
1I85_A
1ID2_A
1IEZ_A
1IJY_A
1IOO_A
1IPI_A
1IQV_A
1IRS_A
1IS7_K
1IUJ_B
1IUY_A
1J1V_A
1J3W_A
1J8I_A
1JC7_A
1JEI_A
1JIW_I
1JJ2_S
1JLI_A
1JMT_A
1JO0_A
1JOF_A
1JOP_A
1JPY_Y
1JR8_A
1JSG_A
1K1Z_A
1K3B_A
1K3S_A
1K5D_B
1K73_I
1KA8_A
1KN6_A
1KOH_D
1KPT_A
1KQ6_A
1KSX_A
1KX5_D
1L1D_B
1L2P_A
1L3G_A
1LDD_A
1LE2_A
1LFU_P
1LMI_A
1LNW_C
1LQM_H
1LR1_B
1LWB_A
1LYV_A
1LZW_B
1M7Y_A
1MAI_A
1MC2_A
1MFQ_C
1MHD_A
1MKF_A

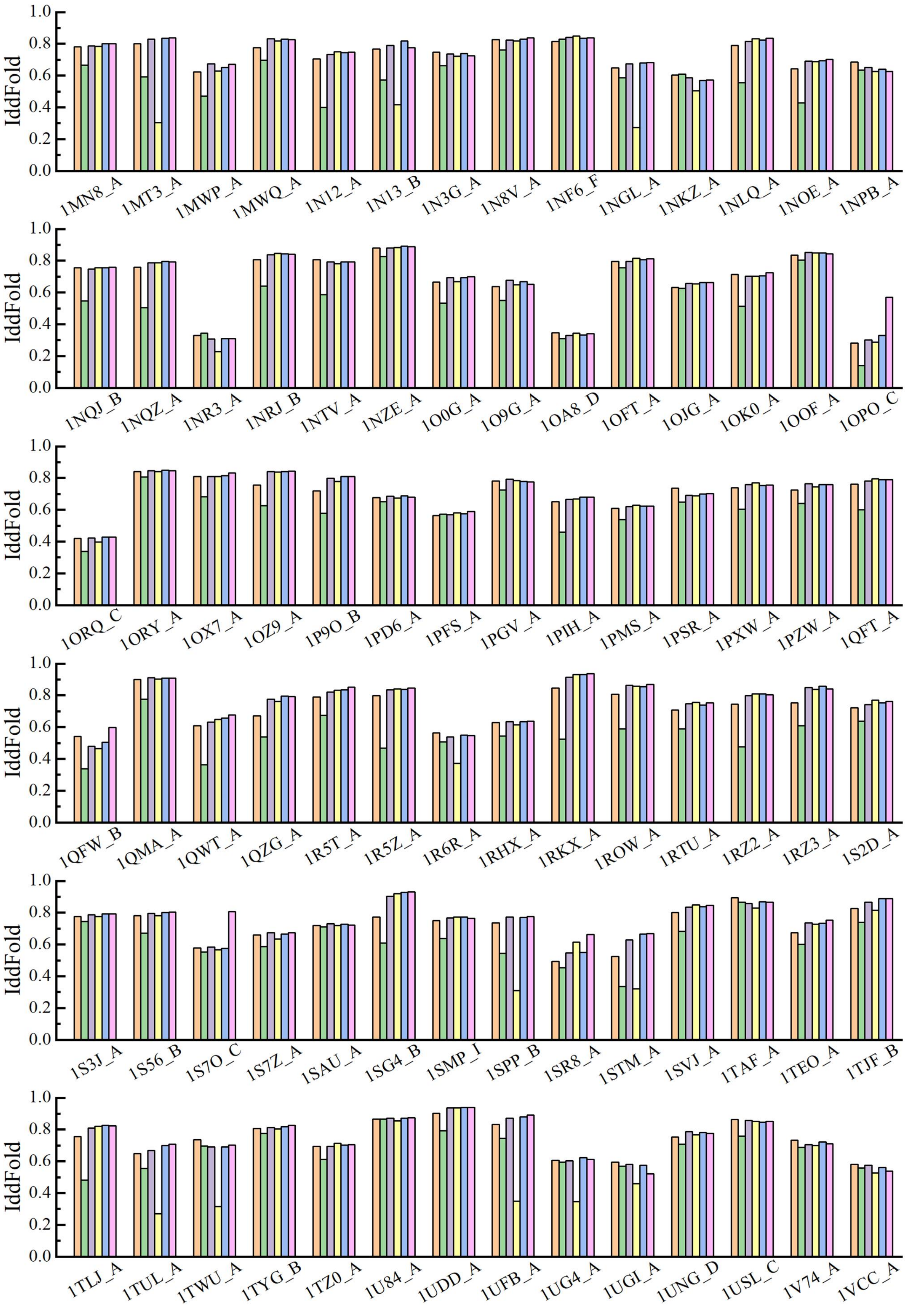
IddFold
0.0
0.2
0.4
0.6
0.8
1.0
1MN8_A
1MT3_A
1MWP_A
1MWQ_A
1N12_A
1N13_B
1N3G_A
1N8V_A
1NF6_F
1NGL_A
1NKZ_A
1NLQ_A
1NOE_A
1NPB_A
1NQJ_B
1NQZ_A
1NR3_A
1NRJ_B
1NTV_A
1NZE_A
1O0G_A
1O9G_A
1OA8_D
1OFT_A
1OJG_A
1OK0_A
1OOF_A
1OPO_C
1ORQ_C
1ORY_A
1OX7_A
1OZ9_A
1P9O_B
1PD6_A
1PFS_A
1PGV_A
1PIH_A
1PMS_A
1PSR_A
1PXW_A
1PZW_A
1QFT_A
1QFW_B
1QMA_A
1QWT_A
1QZG_A
1R5T_A
1R5Z_A
1R6R_A
1RHX_A
1RKX_A
1ROW_A
1RTU_A
1RZ2_A
1RZ3_A
1S2D_A
1S3J_A
1S56_B
1S7O_C
1S7Z_A
1SAU_A
1SG4_B
1SMP_I
1SPP_B
1SR8_A
1STM_A
1SVJ_A
1TAF_A
1TEO_A
1TJF_B
1TLJ_A
1TUL_A
1TWU_A
1TYG_B
1TZ0_A
1U84_A
1UDD_A
1UFB_A
1UG4_A
1UGI_A
1UNG_D
1USL_C
1V74_A
1VCC_A

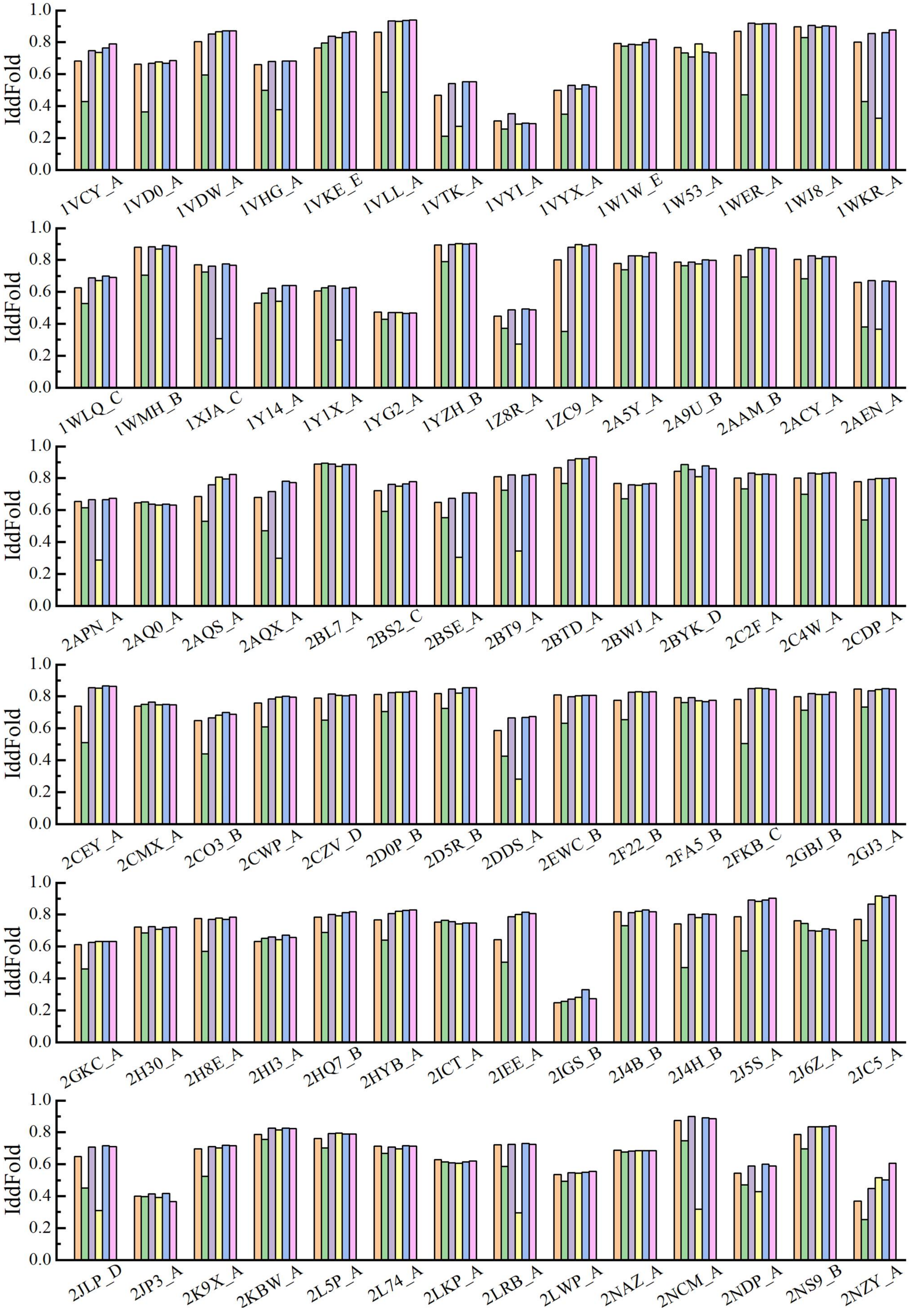
IddFold
1VCY_A
1VD0_A
1VDW_A
1VHG_A
1VKE_E
1VLL_A
1VTK_A
1VYI_A
1VYX_A
1W1W_E
1W53_A
1WER_A
1WJ8_A
1WKR_A
1WLQ_C
1WMH_B
1XJA_C
1Y14_A
1Y1X_A
1YG2_A
1YZH_B
1Z8R_A
1ZC9_A
2A5Y_A
2A9U_B
2AAM_B
2ACY_A
2AEN_A
2APN_A
2AQ0_A
2AQS_A
2AQX_A
2BL7_A
2BS2_C
2BSE_A
2BT9_A
2BTD_A
2BWJ_A
2BYK_D
2C2F_A
2C4W_A
2CDP_A
2CEY_A
2CMX_A
2CO3_B
2CWP_A
2CZV_D
2D0P_B
2D5R_B
2DDS_A
2EWC_B
2F22_B
2FA5_B
2FKB_C
2GBJ_B
2GJ3_A
2GKC_A
2H30_A
2H8E_A
2HI3_A
2HQ7_B
2HYB_A
2ICT_A
2IEE_A
2IGS_B
2J4B_B
2J4H_B
2J5S_A
2J6Z_A
2JC5_A
2JLP_D
2JP3_A
2K9X_A
2KBW_A
2L5P_A
2L74_A
2LKP_A
2LRB_A
2LWP_A
2NAZ_A
2NCM_A
2NDP_A
2NS9_B
2NZY_A

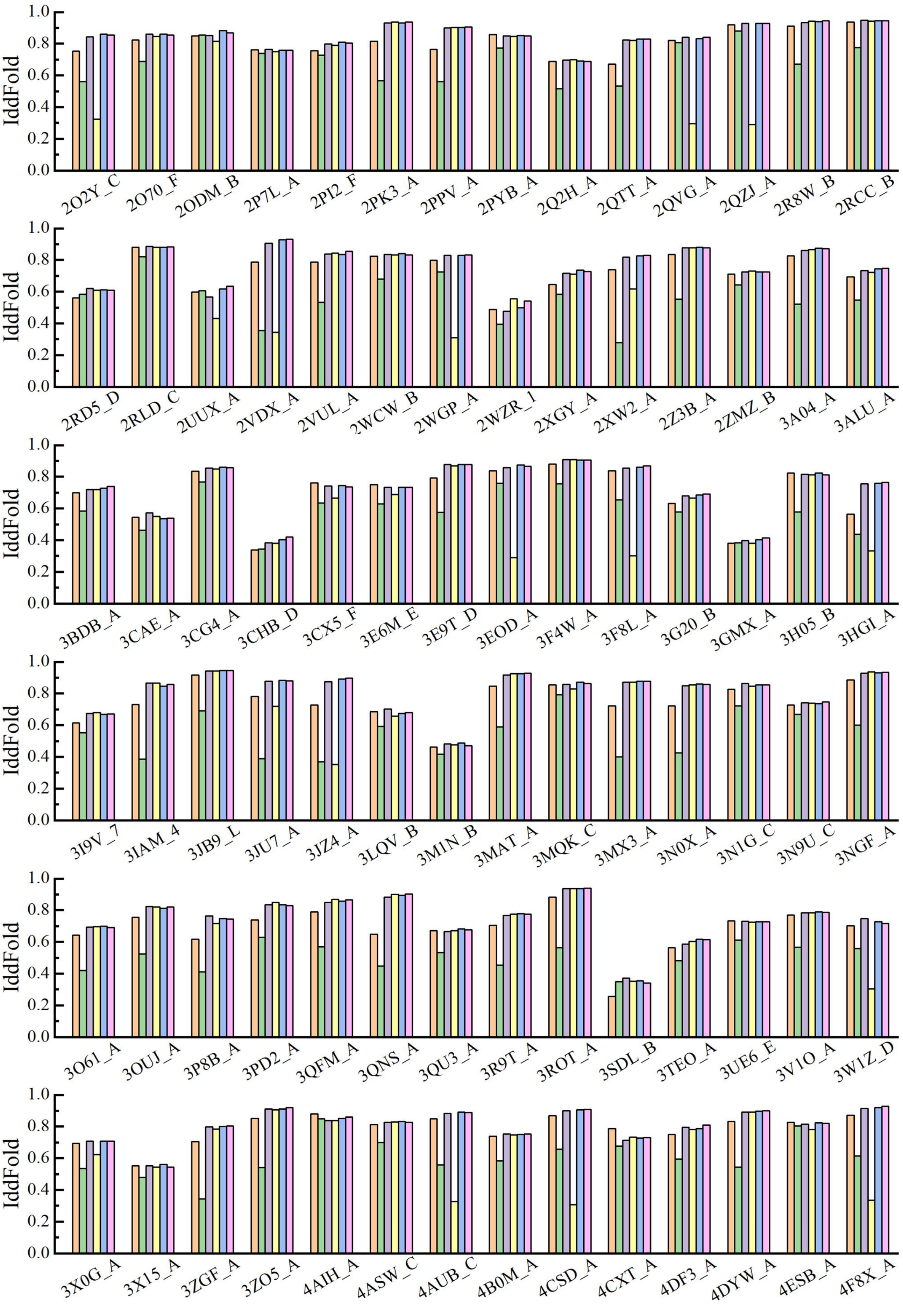
IddFold
1.0
0.8
0.6
0.4
0.2
0.0
2O2Y_C 2O70_F 2ODM_B 2P7L_A 2PI2_F 2PK3_A 2PPV_A 2PYB_A 2Q2H_A 2QTT_A 2QVG_A 2QZJ_A 2R8W_B 2RCC_B
2RD5_D 2RLD_C 2UUX_A 2VDX_A 2VUL_A 2WCW_B 2WGP_A 2WZR_1 2XGY_A 2XW2_A 2Z3B_A 2ZMZ_B 3A04_A 3ALU_A
3BDB_A 3CAE_A 3CG4_A 3CHB_D 3CX5_F 3E6M_E 3E9T_D 3EOD_A 3F4W_A 3F8L_A 3G20_B 3GMX_A 3H05_B 3HGI_A
3I9V_7 3IAM_4 3JB9_L 3JU7_A 3JZ4_A 3LQV_B 3M1N_B 3MAT_A 3MQK_C 3MX3_A 3N0X_A 3N1G_C 3N9U_C 3NGF_A
3O61_A 3OUJ_A 3P8B_A 3PD2_A 3QFM_A 3QNS_A 3QU3_A 3R9T_A 3ROT_A 3SDL_B 3TEO_A 3UE6_E 3V1O_A 3W1Z_D
3X0G_A 3X15_A 3ZGF_A 3ZO5_A 4AIH_A 4ASW_C 4AUB_C 4B0M_A 4CSD_A 4CXT_A 4DF3_A 4DYW_A 4ESB_A 4F8X_A

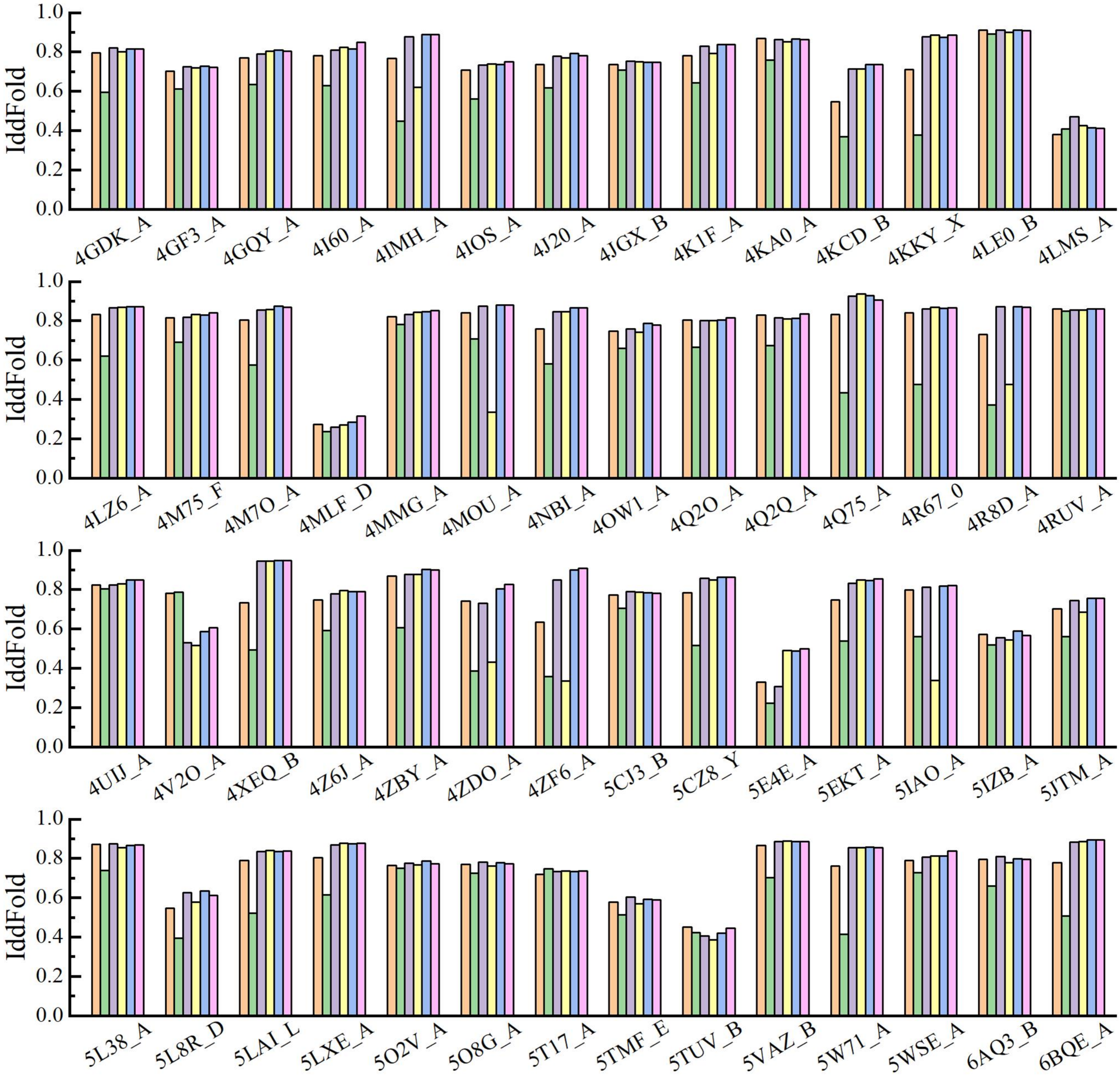

IddFold
1.0
0.8
0.6
0.4
0.2
0.0
4GDK_A
4GF3_A
4GQY_A
4I60_A
4IMH_A
4IOS_A
4J20_A
4JGX_B
4K1F_A
4KA0_A
4KCD_B
4KKY_X
4LE0_B
4LMS_A
4LZ6_A
4M75_F
4M7O_A
4MLF_D
4MMG_A
4MOU_A
4NBI_A
4OW1_A
4Q2O_A
4Q2Q_A
4Q75_A
4R67_0
4R8D_A
4RUV_A
4UIJ_A
4V2O_A
4XEQ_B
4Z6J_A
4ZBY_A
4ZDO_A
4ZF6_A
5CJ3_B
5CZ8_Y
5E4E_A
5EKT_A
5IAO_A
5IZB_A
5JTM_A
5L38_A
5L8R_D
5LAI_L
5LXE_A
5O2V_A
5O8G_A
5T17_A
5TMF_E
5TUV_B
5VAZ_B
5W71_A
5WSE_A
6AQ3_B
6BQE_A

Table S1: Detailed information of 462 proteins in the benchmark dataset.

| PDB | Type | Size | PDB | Type | Size | PDB | Type | Size | PDB | Type | Size | PDB | Type | Size |
|---|---|---|---|---|---|---|---|---|---|---|---|---|---|---|
| 1A3A_C | $\alpha/\beta$ | 146 | 1F1E_A | $\alpha/\beta$ | 151 | 1IUJ_B | $\alpha/\beta$ | 103 | 1N13_B | $\alpha/\beta$ | 112 | 1S7O_C | $\alpha$ | 108 |
| 1A40_A | $\alpha/\beta$ | 321 | 1F2R_I | $\alpha/\beta$ | 100 | 1IUY_A | $\alpha/\beta$ | 92 | 1N3G_A | $\alpha/\beta$ | 113 | 1S7Z_A | $\alpha$ | 106 |
| 1A6L_A | $\alpha/\beta$ | 106 | 1F3Y_A | $\alpha/\beta$ | 165 | 1J1V_A | $\alpha$ | 94 | 1N8V_A | $\alpha$ | 101 | 1SAU_A | $\alpha/\beta$ | 114 |
| 1A7D_A | $\alpha$ | 118 | 1F43_A | $\alpha$ | 61 | 1J3W_A | $\alpha/\beta$ | 134 | 1NF6_F | $\alpha/\beta$ | 171 | 1SG4_B | $\alpha/\beta$ | 258 |
| 1A91_A | $\alpha$ | 79 | 1F5X_A | $\alpha$ | 208 | 1J8I_A | $\alpha/\beta$ | 93 | 1NGL_A | $\alpha/\beta$ | 179 | 1SMP_I | $\alpha/\beta$ | 100 |
| 1ABT_A | $\alpha/\beta$ | 74 | 1F93_A | $\alpha/\beta$ | 103 | 1JC7_A | $\alpha/\beta$ | 129 | 1NKZ_A | $\alpha$ | 53 | 1SPP_B | $\alpha/\beta$ | 112 |
| 1ABV_A | $\alpha$ | 105 | 1F98_A | $\alpha/\beta$ | 125 | 1JEI_A | $\alpha$ | 53 | 1NLQ_A | $\alpha/\beta$ | 105 | 1SR8_A | $\alpha/\beta$ | 282 |
| 1AHK_A | $\beta$ | 129 | 1F9P_A | $\alpha/\beta$ | 81 | 1JIW_I | $\alpha/\beta$ | 105 | 1NOE_A | $\alpha/\beta$ | 86 | 1STM_A | $\alpha/\beta$ | 141 |
| 1AK6_A | $\alpha/\beta$ | 174 | 1FC3_A | $\alpha$ | 116 | 1JJ2_S | $\alpha/\beta$ | 119 | 1NPB_A | $\alpha/\beta$ | 140 | 1SVJ_A | $\alpha/\beta$ | 136 |
| 1AKP_A | $\beta$ | 114 | 1FCA_A | $\alpha/\beta$ | 55 | 1JLI_A | $\alpha$ | 112 | 1NQJ_B | $\alpha/\beta$ | 114 | 1TAF_A | $\alpha$ | 68 |
| 1AOX_A | $\alpha/\beta$ | 201 | 1FEX_A | $\alpha$ | 59 | 1JMT_A | $\alpha/\beta$ | 98 | 1NQZ_A | $\alpha/\beta$ | 171 | 1TEO_A | $\alpha/\beta$ | 173 |
| 1AP7_A | $\alpha$ | 168 | 1FHT_A | $\alpha/\beta$ | 116 | 1JO0_A | $\alpha/\beta$ | 97 | 1NR3_A | $\alpha/\beta$ | 122 | 1TJF_B | $\alpha/\beta$ | 186 |
| 1AUU_A | $\beta$ | 55 | 1FJG_H | $\alpha/\beta$ | 138 | 1JOF_A | $\alpha/\beta$ | 365 | 1NRJ_B | $\alpha/\beta$ | 191 | 1TLJ_A | $\alpha/\beta$ | 189 |
| 1AX8_A | $\alpha/\beta$ | 130 | 1FMB_A | $\alpha/\beta$ | 104 | 1JOP_A | $\alpha/\beta$ | 140 | 1NTV_A | $\alpha/\beta$ | 152 | 1TUL_A | $\alpha/\beta$ | 102 |
| 1AZ0_A | $\alpha/\beta$ | 237 | 1FR0_A | $\alpha$ | 125 | 1JPY_Y | $\alpha/\beta$ | 120 | 1NZE_A | $\alpha$ | 112 | 1TWU_A | $\alpha/\beta$ | 137 |
| 1B4B_A | $\alpha/\beta$ | 72 | 1FRD_A | $\alpha/\beta$ | 98 | 1JR8_A | $\alpha$ | 105 | 1O0G_A | $\alpha/\beta$ | 124 | 1TYG_B | $\alpha/\beta$ | 65 |
| 1B4R_A | $\beta$ | 80 | 1FSP_A | $\alpha/\beta$ | 124 | 1JSG_A | $\alpha/\beta$ | 111 | 1O9G_A | $\alpha/\beta$ | 249 | 1TZ0_A | $\alpha/\beta$ | 108 |
| 1B4U_A | $\alpha$ | 132 | 1FSU_A | $\alpha/\beta$ | 475 | 1K1Z_A | $\beta$ | 78 | 1OA8_D | $\alpha/\beta$ | 133 | 1U84_A | $\alpha$ | 81 |
| 1B8Q_A | $\alpha/\beta$ | 127 | 1FW9_A | $\alpha/\beta$ | 164 | 1K3B_A | $\alpha/\beta$ | 119 | 1OFT_A | $\alpha/\beta$ | 119 | 1UDD_A | $\alpha$ | 215 |
| 1BE3_J | $\alpha$ | 62 | 1FY2_A | $\alpha/\beta$ | 220 | 1K3S_A | $\alpha/\beta$ | 109 | 1OJG_A | $\alpha/\beta$ | 136 | 1UFB_A | $\alpha$ | 127 |
| 1BG8_A | $\alpha/\beta$ | 76 | 1G2R_A | $\alpha/\beta$ | 94 | 1K5D_B | $\alpha/\beta$ | 146 | 1OK0_A | $\beta$ | 74 | 1UG4_A | $\beta$ | 60 |
| 1BGF_A | $\alpha/\beta$ | 124 | 1G8Q_A | $\alpha$ | 90 | 1K73_1 | $\alpha/\beta$ | 73 | 1OOF_A | $\alpha/\beta$ | 124 | 1UGI_A | $\alpha/\beta$ | 83 |
| 1BJX_A | $\alpha/\beta$ | 110 | 1GGS_A | $\alpha/\beta$ | 81 | 1KA8_A | $\alpha/\beta$ | 100 | 1OPO_C | $\alpha/\beta$ | 268 | 1UNG_D | $\alpha$ | 149 |
| 1BUO_A | $\alpha/\beta$ | 121 | 1GME_A | $\alpha/\beta$ | 150 | 1KN6_A | $\alpha/\beta$ | 73 | 1ORQ_C | $\alpha$ | 223 | 1USL_C | $\alpha/\beta$ | 158 |
| 1C03_A | $\alpha/\beta$ | 163 | 1GPQ_B | $\alpha/\beta$ | 128 | 1KOH_D | $\alpha/\beta$ | 172 | 1ORY_A | $\alpha$ | 119 | 1V74_A | $\alpha/\beta$ | 107 |
| 1C1K_A | $\alpha/\beta$ | 217 | 1GQA_A | $\alpha/\beta$ | 130 | 1KPT_A | $\alpha/\beta$ | 105 | 1OX7_A | $\alpha/\beta$ | 158 | 1VCC_A | $\alpha/\beta$ | 77 |
| 1C41_A | $\alpha/\beta$ | 165 | 1GVP_A | $\alpha/\beta$ | 87 | 1KQ6_A | $\alpha/\beta$ | 140 | 1OZ9_A | $\alpha/\beta$ | 141 | 1VCY_A | $\alpha/\beta$ | 193 |
| 1C5E_A | $\alpha/\beta$ | 95 | 1GXD_C | $\alpha/\beta$ | 192 | 1KSX_A | $\alpha/\beta$ | 144 | 1P9O_B | $\alpha/\beta$ | 280 | 1VD0_A | $\alpha/\beta$ | 109 |
| 1CDB_A | $\beta$ | 105 | 1GXL_A | $\alpha/\beta$ | 205 | 1KX5_D | $\alpha$ | 107 | 1PD6_A | $\beta$ | 94 | 1VDW_A | $\alpha/\beta$ | 248 |
| 1CF7_B | $\alpha/\beta$ | 82 | 1GYH_A | $\alpha/\beta$ | 318 | 1L1D_B | $\alpha/\beta$ | 147 | 1PFS_A | $\beta$ | 78 | 1VHG_A | $\alpha/\beta$ | 185 |
| 1CQA_A | $\alpha/\beta$ | 123 | 1H4L_D | $\alpha$ | 147 | 1L2P_A | $\alpha$ | 61 | 1PGV_A | $\alpha/\beta$ | 167 | 1VKE_E | $\alpha$ | 119 |
| 1CTO_A | $\beta$ | 109 | 1H8E_H | $\alpha/\beta$ | 89 | 1L3G_A | $\alpha/\beta$ | 123 | 1PIH_A | $\alpha/\beta$ | 73 | 1VLL_A | $\alpha/\beta$ | 321 |
| 1CXZ_B | $\alpha$ | 86 | 1H9E_A | $\alpha$ | 56 | 1LDD_A | $\alpha/\beta$ | 71 | 1PMS_A | $\alpha/\beta$ | 135 | 1VTK_A | $\alpha/\beta$ | 313 |
| 1D6T_A | $\alpha/\beta$ | 117 | 1H9F_A | $\alpha$ | 57 | 1LE2_A | $\alpha$ | 144 | 1PSR_A | $\alpha/\beta$ | 100 | 1VYI_A | $\alpha/\beta$ | 111 |
| 1D8B_A | $\alpha$ | 81 | 1HBG_A | $\alpha$ | 147 | 1LFU_P | $\alpha$ | 82 | 1PXW_A | $\alpha/\beta$ | 128 | 1VYX_A | $\alpha/\beta$ | 60 |
| 1DBF_A | $\alpha/\beta$ | 127 | 1HBX_E | $\alpha/\beta$ | 89 | 1LMI_A | $\alpha/\beta$ | 131 | 1PZW_A | $\alpha/\beta$ | 80 | 1W1W_E | $\alpha/\beta$ | 70 |
| 1DCF_A | $\alpha/\beta$ | 133 | 1HCD_A | $\alpha/\beta$ | 118 | 1LNW_C | $\alpha/\beta$ | 139 | 1QFT_A | $\alpha/\beta$ | 175 | 1W53_A | $\alpha$ | 84 |
| 1DJ7_A | $\alpha/\beta$ | 109 | 1HH8_A | $\alpha/\beta$ | 192 | 1LQM_H | $\alpha/\beta$ | 84 | 1QFW_B | $\beta$ | 110 | 1WER_A | $\alpha$ | 324 |
| 1DL6_A | $\beta$ | 58 | 1HHV_A | $\alpha/\beta$ | 74 | 1LR1_B | $\alpha$ | 57 | 1QMA_A | $\alpha/\beta$ | 123 | 1WJ8_A | $\alpha$ | 117 |
| 1DOI_A | $\alpha/\beta$ | 128 | 1HKQ_A | $\alpha/\beta$ | 125 | 1LWB_A | $\alpha$ | 122 | 1QWT_A | $\alpha/\beta$ | 239 | 1WKR_A | $\alpha/\beta$ | 340 |
| 1DP7_P | $\alpha/\beta$ | 76 | 1HKX_E | $\alpha/\beta$ | 143 | 1LYV_A | $\alpha/\beta$ | 283 | 1QZG_A | $\alpha/\beta$ | 170 | 1WLQ_C | $\alpha/\beta$ | 185 |
| 1DTP_A | $\alpha/\beta$ | 190 | 1HL6_D | $\alpha/\beta$ | 143 | 1LZW_B | $\alpha/\beta$ | 146 | 1R5T_A | $\alpha/\beta$ | 141 | 1WMH_B | $\alpha/\beta$ | 82 |
| 1DUN_A | $\alpha/\beta$ | 120 | 1I27_A | $\alpha/\beta$ | 73 | 1M7Y_A | $\alpha/\beta$ | 424 | 1R5Z_A | $\alpha$ | 320 | 1XJA_C | $\alpha/\beta$ | 169 |
| 1DWM_A | $\alpha/\beta$ | 69 | 1I35_A | $\alpha/\beta$ | 95 | 1MAI_A | $\alpha/\beta$ | 119 | 1R6R_A | $\alpha$ | 80 | 1Y14_A | $\alpha$ | 133 |
| 1E2A_A | $\alpha$ | 102 | 1I85_A | $\beta$ | 110 | 1MC2_A | $\alpha/\beta$ | 122 | 1RHX_A | $\alpha/\beta$ | 87 | 1Y1X_A | $\alpha/\beta$ | 182 |
| 1E3Y_A | $\alpha$ | 104 | 1ID2_A | $\alpha/\beta$ | 106 | 1MFQ_C | $\alpha$ | 108 | 1RKX_A | $\alpha/\beta$ | 349 | 1YG2_A | $\alpha/\beta$ | 169 |
| 1E53_A | $\alpha/\beta$ | 59 | 1IEZ_A | $\alpha/\beta$ | 217 | 1MHD_A | $\alpha/\beta$ | 123 | 1ROW_A | $\alpha/\beta$ | 107 | 1YZH_B | $\alpha/\beta$ | 206 |
| 1EGG_B | $\alpha/\beta$ | 144 | 1IJY_A | $\alpha/\beta$ | 122 | 1MKF_A | $\alpha/\beta$ | 371 | 1RTU_A | $\alpha/\beta$ | 114 | 1Z8R_A | $\alpha/\beta$ | 150 |
| 1EKZ_A | $\alpha/\beta$ | 76 | 1IOO_A | $\alpha/\beta$ | 196 | 1MN8_A | $\alpha$ | 95 | 1RZ2_A | $\alpha/\beta$ | 214 | 1ZC9_A | $\alpha/\beta$ | 431 |
| 1ELW_A | $\alpha$ | 117 | 1IPI_A | $\alpha/\beta$ | 114 | 1MT3_A | $\alpha/\beta$ | 292 | 1RZ3_A | $\alpha/\beta$ | 184 | 2A5Y_A | $\alpha$ | 173 |
| 1EM8_D | $\alpha/\beta$ | 112 | 1IQV_A | $\alpha/\beta$ | 166 | 1MWP_A | $\alpha/\beta$ | 96 | 1S2D_A | $\alpha/\beta$ | 165 | 2A9U_B | $\alpha$ | 127 |
| 1EZV_G | $\alpha/\beta$ | 81 | 1IRS_A | $\alpha/\beta$ | 112 | 1MWQ_A | $\alpha/\beta$ | 99 | 1S3J_A | $\alpha/\beta$ | 143 | 2AAM_B | $\alpha/\beta$ | 287 |
| 1F15_C | $\alpha/\beta$ | 191 | 1IS7_K | $\alpha/\beta$ | 85 | 1N12_A | $\alpha/\beta$ | 138 | 1S56_B | $\alpha/\beta$ | 135 | 2ACY_A | $\alpha/\beta$ | 98 |

Continued on next page

Continue to last page

| PDB | Type | Size | PDB | Type | Size | PDB | Type | Size | PDB | Type | Size | PDB | Type | Size |
|---|---|---|---|---|---|---|---|---|---|---|---|---|---|---|
| 2AEN_A | $\alpha/\beta$ | 164 | 2J5S_A | $\alpha/\beta$ | 250 | 2XW2_A | $\alpha/\beta$ | 381 | 3R9T_A | $\alpha/\beta$ | 260 | 4MOU_A | $\alpha/\beta$ | 261 |
| 2APN_A | $\alpha/\beta$ | 114 | 2J6Z_A | $\alpha$ | 86 | 2Z3B_A | $\alpha/\beta$ | 180 | 3ROT_A | $\alpha/\beta$ | 273 | 4NBI_A | $\alpha/\beta$ | 163 |
| 2AQ0_A | $\alpha$ | 84 | 2JC5_A | $\alpha/\beta$ | 259 | 2ZMZ_B | $\alpha/\beta$ | 79 | 3SDL_B | $\alpha$ | 97 | 4OW1_A | $\alpha/\beta$ | 86 |
| 2AQS_A | $\alpha/\beta$ | 160 | 2JLP_D | $\alpha/\beta$ | 169 | 3A04_A | $\alpha/\beta$ | 367 | 3TEO_A | $\alpha/\beta$ | 202 | 4Q2O_A | $\alpha/\beta$ | 92 |
| 2AQX_A | $\alpha/\beta$ | 289 | 2JP3_A | $\alpha$ | 67 | 3ALU_A | $\alpha/\beta$ | 157 | 3UE6_E | $\alpha/\beta$ | 138 | 4Q2Q_A | $\alpha/\beta$ | 90 |
| 2BL7_A | $\alpha$ | 79 | 2K9X_A | $\alpha/\beta$ | 102 | 3BDB_A | $\alpha/\beta$ | 126 | 3V1O_A | $\alpha/\beta$ | 165 | 4Q75_A | $\alpha/\beta$ | 413 |
| 2BS2_C | $\alpha$ | 254 | 2KBW_A | $\alpha$ | 160 | 3CAE_A | $\alpha/\beta$ | 132 | 3W1Z_D | $\alpha/\beta$ | 110 | 4R67_0 | $\alpha/\beta$ | 199 |
| 2BSE_A | $\alpha/\beta$ | 107 | 2L5P_A | $\alpha/\beta$ | 175 | 3CG4_A | $\alpha/\beta$ | 126 | 3X0G_A | $\alpha$ | 93 | 4R8D_A | $\alpha/\beta$ | 369 |
| 2BT9_A | $\beta$ | 90 | 2L74_A | $\alpha/\beta$ | 125 | 3CHB_D | $\alpha/\beta$ | 103 | 3X15_A | $\alpha$ | 87 | 4RUV_A | $\alpha/\beta$ | 106 |
| 2BTD_A | $\alpha/\beta$ | 209 | 2LKP_A | $\alpha/\beta$ | 119 | 3CX5_F | $\alpha$ | 74 | 3ZGF_A | $\alpha/\beta$ | 291 | 4UIJ_A | $\alpha/\beta$ | 104 |
| 2BWJ_A | $\alpha/\beta$ | 196 | 2LRB_A | $\alpha/\beta$ | 165 | 3E6M_E | $\alpha/\beta$ | 147 | 3ZO5_A | $\alpha/\beta$ | 230 | 4V2O_A | $\alpha$ | 78 |
| 2BYK_D | $\alpha$ | 92 | 2LWP_A | $\alpha/\beta$ | 97 | 3E9T_D | $\beta$ | 102 | 4AIH_A | $\alpha/\beta$ | 139 | 4XEQ_B | $\alpha/\beta$ | 304 |
| 2C2F_A | $\alpha/\beta$ | 178 | 2NAZ_A | $\alpha/\beta$ | 109 | 3EOD_A | $\alpha/\beta$ | 115 | 4ASW_C | $\alpha/\beta$ | 81 | 4Z6J_A | $\beta$ | 133 |
| 2C4W_A | $\alpha/\beta$ | 168 | 2NCM_A | $\beta$ | 99 | 3F4W_A | $\alpha/\beta$ | 211 | 4AUB_C | $\alpha/\beta$ | 330 | 4ZBY_A | $\alpha/\beta$ | 194 |
| 2CDP_A | $\alpha/\beta$ | 138 | 2NDP_A | $\alpha/\beta$ | 99 | 3F8L_A | $\alpha/\beta$ | 162 | 4B0M_A | $\alpha/\beta$ | 131 | 4ZDO_A | $\alpha/\beta$ | 445 |
| 2CEY_A | $\alpha/\beta$ | 306 | 2NS9_B | $\alpha/\beta$ | 152 | 3G20_B | $\alpha/\beta$ | 119 | 4CSD_A | $\beta$ | 266 | 4ZF6_A | $\alpha/\beta$ | 460 |
| 2CMX_A | $\alpha/\beta$ | 70 | 2NZY_A | $\alpha/\beta$ | 343 | 3GMX_A | $\alpha/\beta$ | 153 | 4CXT_A | $\alpha/\beta$ | 132 | 5CJ3_B | $\alpha/\beta$ | 126 |
| 2CO3_B | $\alpha/\beta$ | 135 | 2O2Y_C | $\alpha/\beta$ | 293 | 3H05_B | $\alpha/\beta$ | 163 | 4DF3_A | $\alpha/\beta$ | 230 | 5CZ8_Y | $\alpha/\beta$ | 221 |
| 2CWP_A | $\alpha/\beta$ | 109 | 2O70_F | $\alpha$ | 168 | 3HGI_A | $\alpha/\beta$ | 258 | 4DYW_A | $\alpha/\beta$ | 129 | 5E4E_A | $\alpha/\beta$ | 111 |
| 2CZV_D | $\alpha/\beta$ | 119 | 2ODM_B | $\alpha$ | 83 | 3I9V_7 | $\alpha/\beta$ | 127 | 4ESB_A | $\alpha/\beta$ | 103 | 5EKT_A | $\alpha/\beta$ | 196 |
| 2D0P_B | $\alpha/\beta$ | 110 | 2P7L_A | $\alpha/\beta$ | 125 | 3IAM_4 | $\alpha/\beta$ | 378 | 4F8X_A | $\alpha/\beta$ | 335 | 5IAO_A | $\alpha/\beta$ | 171 |
| 2D5R_B | $\alpha/\beta$ | 116 | 2PI2_F | $\alpha/\beta$ | 119 | 3JB9_L | $\beta$ | 293 | 4GDK_A | $\alpha/\beta$ | 88 | 5IZB_A | $\alpha/\beta$ | 89 |
| 2DDS_A | $\alpha/\beta$ | 299 | 2PK3_A | $\alpha/\beta$ | 309 | 3JU7_A | $\alpha/\beta$ | 368 | 4GF3_A | $\alpha/\beta$ | 123 | 5JTM_A | $\alpha/\beta$ | 155 |
| 2EWC_B | $\alpha/\beta$ | 122 | 2PPV_A | $\alpha/\beta$ | 325 | 3JZ4_A | $\alpha/\beta$ | 481 | 4GQY_A | $\alpha/\beta$ | 147 | 5L38_A | $\alpha/\beta$ | 91 |
| 2F22_B | $\alpha/\beta$ | 143 | 2PYB_A | $\alpha$ | 151 | 3LQV_B | $\alpha/\beta$ | 115 | 4I60_A | $\alpha/\beta$ | 128 | 5L8R_D | $\alpha/\beta$ | 143 |
| 2FA5_B | $\alpha/\beta$ | 142 | 2Q2H_A | $\alpha/\beta$ | 118 | 3M1N_B | $\alpha/\beta$ | 168 | 4IMH_A | $\alpha/\beta$ | 347 | 5LAI_L | $\alpha/\beta$ | 222 |
| 2FKB_C | $\alpha/\beta$ | 167 | 2QTT_A | $\alpha/\beta$ | 248 | 3MAT_A | $\alpha/\beta$ | 264 | 4IOS_A | $\alpha/\beta$ | 100 | 5LXE_A | $\alpha/\beta$ | 324 |
| 2GBJ_B | $\alpha/\beta$ | 84 | 2QVG_A | $\alpha/\beta$ | 129 | 3MQK_C | $\beta$ | 75 | 4J20_A | $\alpha/\beta$ | 88 | 5O2V_A | $\alpha/\beta$ | 92 |
| 2GJ3_A | $\alpha/\beta$ | 119 | 2QZJ_A | $\alpha/\beta$ | 121 | 3MX3_A | $\alpha/\beta$ | 470 | 4JGX_B | $\alpha/\beta$ | 128 | 5O8G_A | $\alpha/\beta$ | 122 |
| 2GKC_A | $\alpha/\beta$ | 155 | 2R8W_B | $\alpha/\beta$ | 299 | 3N0X_A | $\alpha/\beta$ | 371 | 4K1F_A | $\alpha/\beta$ | 198 | 5T17_A | $\alpha/\beta$ | 85 |
| 2H30_A | $\alpha/\beta$ | 151 | 2RCC_B | $\alpha$ | 280 | 3N1G_C | $\alpha/\beta$ | 104 | 4KA0_A | $\alpha/\beta$ | 143 | 5TMF_E | $\alpha/\beta$ | 95 |
| 2H8E_A | $\alpha/\beta$ | 120 | 2RD5_D | $\alpha/\beta$ | 126 | 3N9U_C | $\alpha/\beta$ | 96 | 4KCD_B | $\alpha/\beta$ | 291 | 5TUV_B | $\alpha/\beta$ | 104 |
| 2HI3_A | $\alpha$ | 73 | 2RLD_C | $\alpha$ | 116 | 3NGF_A | $\alpha/\beta$ | 258 | 4KKY_X | $\alpha/\beta$ | 411 | 5VAZ_B | $\alpha/\beta$ | 321 |
| 2HQ7_B | $\alpha/\beta$ | 142 | 2UUX_A | $\alpha/\beta$ | 55 | 3O61_A | $\alpha/\beta$ | 187 | 4LE0_B | $\alpha/\beta$ | 133 | 5W71_A | $\alpha/\beta$ | 413 |
| 2HYB_A | $\alpha/\beta$ | 130 | 2VDX_A | $\alpha/\beta$ | 366 | 3OUJ_A | $\alpha/\beta$ | 226 | 4LMS_A | $\alpha/\beta$ | 80 | 5WSE_A | $\alpha/\beta$ | 114 |
| 2ICT_A | $\alpha$ | 94 | 2VUL_A | $\alpha/\beta$ | 193 | 3P8B_A | $\alpha/\beta$ | 60 | 4LZ6_A | $\alpha$ | 446 | 6AQ3_B | $\alpha/\beta$ | 171 |
| 2IEE_A | $\alpha/\beta$ | 251 | 2WCW_B | $\alpha/\beta$ | 122 | 3PD2_A | $\alpha/\beta$ | 147 | 4M75_F | $\alpha/\beta$ | 75 | 6BQE_A | $\alpha/\beta$ | 240 |
| 2IGS_B | $\alpha/\beta$ | 214 | 2WGP_A | $\alpha/\beta$ | 168 | 3QFM_A | $\alpha/\beta$ | 258 | 4M7O_A | $\alpha/\beta$ | 258 | | | |
| 2J4B_B | $\alpha/\beta$ | 133 | 2WZR_1 | $\alpha/\beta$ | 192 | 3QNS_A | $\alpha/\beta$ | 311 | 4MLF_D | $\beta$ | 61 | | | |
| 2J4H_B | $\alpha/\beta$ | 172 | 2XGY_A | $\alpha$ | 129 | 3QU3_A | $\alpha/\beta$ | 122 | 4MMG_A | $\alpha/\beta$ | 91 | | | |

Table S2: Results of the first model predicted by IddFold, Rosetta_D, IddFold_relax, trRosetta_D, IddFold_relax2 and trRosetta on the benchmark dataset.

| No. | PDB | IddFold | | Rosetta_D | | IddFold_relax | | trRosetta_D | | IddFold_relax2 | | trRosetta | |
|---|---|---|---|---|---|---|---|---|---|---|---|---|---|
| | | RMSD | TM-score | RMSD | TM-score | RMSD | TM-score | RMSD | TM-score | RMSD | TM-score | RMSD | TM-score |
| 1 | 1A3A_C | 2.359 | 0.8503 | 4.6815 | 0.6633 | 2.178 | 0.8594 | 1.988 | 0.8722 | 2.031 | 0.8695 | 1.984 | 0.87 |
| 2 | 1A40_A | 6.344 | 0.625 | 8.457 | 0.5323 | 1.747 | 0.9406 | 16.189 | 0.3545 | 1.693 | 0.9432 | 1.628 | 0.9463 |
| 3 | 1A6L_A | 2.836 | 0.7493 | 3.3745 | 0.7025 | 2.134 | 0.8032 | 2.131 | 0.8025 | 2.083 | 0.8078 | 2.226 | 0.7877 |
| 4 | 1A7D_A | 3.623 | 0.8702 | 4.0035 | 0.7784 | 3.642 | 0.8838 | 3.436 | 0.8781 | 3.963 | 0.8793 | 3.869 | 0.8787 |
| 5 | 1A91_A | 4.02 | 0.5135 | 4.279 | 0.5176 | 3.907 | 0.5226 | 4.523 | 0.4757 | 3.917 | 0.518 | 3.926 | 0.5158 |
| 6 | 1ABT_A | 4.315 | 0.5721 | 4.08 | 0.5287 | 4.77 | 0.5615 | 9.736 | 0.3299 | 5.121 | 0.5531 | 4.714 | 0.5573 |
| 7 | 1ABV_A | 2.302 | 0.7977 | 2.655 | 0.7959 | 2.354 | 0.8049 | 2.488 | 0.7818 | 2.359 | 0.8036 | 2.48 | 0.7933 |
| 8 | 1AHK_A | 5.921 | 0.5139 | 4.58 | 0.5902 | 5.828 | 0.5238 | 5.814 | 0.5324 | 5.78 | 0.5287 | 5.83 | 0.5291 |
| 9 | 1AK6_A | 6.062 | 0.6347 | 6.646 | 0.5421 | 6.391 | 0.6602 | 6.411 | 0.6677 | 6.527 | 0.6652 | 6.447 | 0.6717 |
| 10 | 1AKP_A | 4.94 | 0.6686 | 5.391 | 0.4893 | 3.122 | 0.7226 | 3.782 | 0.6582 | 3.455 | 0.7095 | 2.958 | 0.7305 |
| 11 | 1AOX_A | 3.763 | 0.7995 | 4.919 | 0.6307 | 3.161 | 0.8442 | 3.027 | 0.8464 | 3.171 | 0.8429 | 3.1 | 0.8469 |
| 12 | 1AP7_A | 3.155 | 0.7664 | 3.553 | 0.7437 | 3.117 | 0.7842 | 3.294 | 0.7881 | 3.033 | 0.7856 | 2.983 | 0.7883 |
| 13 | 1AUU_A | 3.441 | 0.6916 | 3.723 | 0.5726 | 3.906 | 0.6956 | 8.463 | 0.2955 | 3.741 | 0.6982 | 2.773 | 0.7039 |
| 14 | 1AX8_A | 3.195 | 0.7766 | 7.529 | 0.5161 | 3.095 | 0.7967 | 3.107 | 0.7889 | 3.1 | 0.8051 | 3.203 | 0.805 |
| 15 | 1AZ0_A | 7.638 | 0.6407 | 8.965 | 0.4535 | 5.723 | 0.7123 | 15.524 | 0.3378 | 4.649 | 0.7639 | 4.291 | 0.7749 |
| 16 | 1B4B_A | 2.85 | 0.859 | 3.1805 | 0.8659 | 2.772 | 0.9137 | 3.55 | 0.842 | 2.746 | 0.9187 | 2.728 | 0.9147 |
| 17 | 1B4R_A | 2.328 | 0.749 | 2.904 | 0.6367 | 2.15 | 0.7691 | 2.226 | 0.7764 | 2.23 | 0.7765 | 2.256 | 0.7716 |
| 18 | 1B4U_A | 4.496 | 0.6542 | 6.694 | 0.622 | 5.166 | 0.6561 | 6.283 | 0.6409 | 5.35 | 0.6617 | 5.447 | 0.656 |
| 19 | 1B8Q_A | 8.926 | 0.4772 | 10.006 | 0.4628 | 10.816 | 0.4789 | 12.505 | 0.2573 | 9.853 | 0.4833 | 9.948 | 0.4847 |
| 20 | 1BE3_J | 3.519 | 0.5606 | 3.911 | 0.515 | 3.232 | 0.5849 | 3.454 | 0.5488 | 3.394 | 0.5751 | 3.274 | 0.5696 |
| 21 | 1BG8_A | 1.771 | 0.815 | 2.3025 | 0.7808 | 1.766 | 0.8289 | 1.843 | 0.8182 | 1.648 | 0.8459 | 1.556 | 0.8421 |
| 22 | 1BGF_A | 2.507 | 0.8117 | 3.1995 | 0.7666 | 2.462 | 0.8133 | 2.87 | 0.768 | 2.467 | 0.8126 | 2.419 | 0.811 |
| 23 | 1BJX_A | 8.714 | 0.7801 | 7.867 | 0.6615 | 7.194 | 0.7844 | 6.939 | 0.7836 | 7.768 | 0.7876 | 7.706 | 0.7809 |
| 24 | 1BUO_A | 2.428 | 0.8481 | 4.339 | 0.6968 | 5.326 | 0.8038 | 5.435 | 0.813 | 5.781 | 0.8233 | 5.257 | 0.8244 |
| 25 | 1C03_A | 10.259 | 0.6888 | 7.3835 | 0.6687 | 9.05 | 0.687 | 9.433 | 0.6897 | 9.078 | 0.6867 | 11.457 | 0.6887 |
| 26 | 1C1K_A | 8.867 | 0.4859 | 8.5405 | 0.4688 | 8.784 | 0.5255 | 11.623 | 0.4702 | 8.819 | 0.5293 | 8.723 | 0.5177 |
| 27 | 1C41_A | 3.079 | 0.8476 | 4.81 | 0.7252 | 2.685 | 0.8651 | 3.325 | 0.8514 | 3.117 | 0.8741 | 3.731 | 0.8604 |
| 28 | 1C5E_A | 2.934 | 0.7248 | 4.9075 | 0.5111 | 2.021 | 0.8255 | 3.121 | 0.716 | 2 | 0.8266 | 1.952 | 0.8303 |
| 29 | 1CDB_A | 4.821 | 0.7055 | 5.4 | 0.5802 | 4.745 | 0.7311 | 10.891 | 0.324 | 4.693 | 0.7398 | 4.674 | 0.7307 |
| 30 | 1CF7_B | 2.784 | 0.711 | 3.387 | 0.6197 | 2.407 | 0.7914 | 2.053 | 0.8017 | 2.343 | 0.7951 | 2.282 | 0.7948 |
| 31 | 1CQA_A | 2.673 | 0.8246 | 3.728 | 0.676 | 2.681 | 0.826 | 2.514 | 0.8467 | 2.61 | 0.8367 | 2.457 | 0.8465 |
| 32 | 1CTO_A | 5.5 | 0.6841 | 5.57 | 0.5589 | 5.196 | 0.7006 | 10.614 | 0.2948 | 5.168 | 0.6923 | 4.993 | 0.7008 |
| 33 | 1CXZ_B | 2.093 | 0.8649 | 3.172 | 0.6914 | 1.678 | 0.8853 | 2.182 | 0.8334 | 1.702 | 0.8869 | 1.632 | 0.8962 |
| 34 | 1D6T_A | 4.325 | 0.7727 | 4.675 | 0.6707 | 4.437 | 0.7969 | 12.488 | 0.3047 | 4.491 | 0.7976 | 4.464 | 0.7921 |
| 35 | 1D8B_A | 2.494 | 0.7337 | 3.364 | 0.6906 | 2.562 | 0.7337 | 2.522 | 0.7386 | 2.601 | 0.7243 | 2.641 | 0.7174 |
| 36 | 1DBF_A | 3.333 | 0.8018 | 5.065 | 0.6408 | 3.656 | 0.8306 | 3.731 | 0.8314 | 3.312 | 0.843 | 3.17 | 0.8437 |
| 37 | 1DCF_A | 3.738 | 0.8249 | 3.8955 | 0.7635 | 4.139 | 0.8279 | 14.179 | 0.2957 | 3.811 | 0.8407 | 3.76 | 0.8359 |
| 38 | 1DJ7_A | 5.293 | 0.5828 | 8.419 | 0.5197 | 4.839 | 0.63 | 5.77 | 0.5672 | 4.728 | 0.6417 | 4.586 | 0.6557 |
| 39 | 1DL6_A | 16.326 | 0.3652 | 15.424 | 0.3996 | 12.952 | 0.3676 | 12.797 | 0.2092 | 12.88 | 0.3747 | 13.489 | 0.3779 |
| 40 | 1DOI_A | 8.564 | 0.5458 | 7.377 | 0.4153 | 6.614 | 0.5759 | 5.115 | 0.6048 | 6.229 | 0.593 | 6.102 | 0.6163 |
| 41 | 1DP7_P | 1.74 | 0.7979 | 2.638 | 0.7057 | 1.584 | 0.816 | 1.793 | 0.804 | 1.538 | 0.8265 | 1.594 | 0.8137 |
| 42 | 1DTP_A | 13.52 | 0.4031 | 15.409 | 0.2622 | 11.232 | 0.4424 | 15.675 | 0.3209 | 9.91 | 0.4809 | 10.691 | 0.4741 |
| 43 | 1DUN_A | 9.142 | 0.7018 | 8.999 | 0.4551 | 7.652 | 0.7376 | 11.812 | 0.2931 | 6.651 | 0.7385 | 7.91 | 0.7318 |
| 44 | 1DWM_A | 4.346 | 0.6962 | 3.116 | 0.6755 | 3.036 | 0.7348 | 3.074 | 0.7175 | 3.58 | 0.72 | 2.656 | 0.7215 |
| 45 | 1E2A_A | 1.842 | 0.8272 | 2.4635 | 0.8521 | 1.688 | 0.8464 | 2.014 | 0.803 | 1.669 | 0.8502 | 1.697 | 0.8449 |
| 46 | 1E3Y_A | 4.808 | 0.7321 | 6.3495 | 0.6886 | 5.002 | 0.7258 | 4.586 | 0.7091 | 4.943 | 0.7218 | 4.677 | 0.7238 |
| 47 | 1E53_A | 3.418 | 0.6057 | 3.778 | 0.5371 | 3.388 | 0.6216 | 3.377 | 0.6114 | 3.374 | 0.6239 | 3.321 | 0.6223 |
| 48 | 1EGG_B | 10.583 | 0.6821 | 9.415 | 0.5838 | 10.528 | 0.7059 | 15.361 | 0.2864 | 10.456 | 0.7174 | 10.555 | 0.7109 |
| 49 | 1EKZ_A | 5.07 | 0.7079 | 4.567 | 0.6755 | 4.757 | 0.7218 | 4.911 | 0.7229 | 4.803 | 0.7224 | 4.407 | 0.73 |
| 50 | 1ELW_A | 1.507 | 0.8918 | 1.337 | 0.9107 | 1.4 | 0.9032 | 1.513 | 0.8949 | 1.374 | 0.9065 | 1.458 | 0.9013 |
| 51 | 1EM8_D | 4.097 | 0.7131 | 4.549 | 0.5823 | 4.882 | 0.7039 | 4.53 | 0.7016 | 4.638 | 0.7101 | 4.482 | 0.7226 |

Continued on next page

Continue to last page

| No. | PDB | IddFold | | Rosetta_D | | IddFold_relax | | trRosetta_D | | IddFold_relax2 | | trRosetta | |
|---|---|---|---|---|---|---|---|---|---|---|---|---|---|
| | | RMSD | TMscore | RMSD | TMscore | RMSD | TMscore | RMSD | TMscore | RMSD | TMscore | RMSD | TMscore |
| 52 | 1EZV_G | 11.149 | 0.4177 | 18.048 | 0.4187 | 13.397 | 0.4029 | 14.821 | 0.2883 | 9.489 | 0.4122 | 11.055 | 0.4136 |
| 53 | 1F15_C | 22.374 | 0.2279 | 20.8255 | 0.1846 | 17.615 | 0.2649 | 19.576 | 0.2608 | 15.979 | 0.2813 | 20.556 | 0.2774 |
| 54 | 1F1E_A | 4.059 | 0.7149 | 8.3055 | 0.6594 | 3.422 | 0.7984 | 6.049 | 0.6597 | 2.691 | 0.8352 | 2.604 | 0.8431 |
| 55 | 1F2R_I | 4.999 | 0.6798 | 5.018 | 0.6355 | 5.877 | 0.7044 | 5.992 | 0.7136 | 5.816 | 0.7007 | 6.579 | 0.694 |
| 56 | 1F3Y_A | 6.231 | 0.6959 | 9.621 | 0.4226 | 4.669 | 0.7346 | 4.954 | 0.7443 | 4.636 | 0.7555 | 4.556 | 0.7512 |
| 57 | 1F43_A | 6.279 | 0.5648 | 6.509 | 0.5701 | 6.049 | 0.565 | 10.93 | 0.2408 | 6.055 | 0.5703 | 6.365 | 0.5743 |
| 58 | 1F5X_A | 10.065 | 0.7279 | 7.7945 | 0.55 | 8.111 | 0.759 | 7.037 | 0.7587 | 6.747 | 0.7559 | 7.435 | 0.7566 |
| 59 | 1F93_A | 1.831 | 0.8395 | 2.6605 | 0.7749 | 1.915 | 0.8449 | 10.528 | 0.3292 | 1.757 | 0.8496 | 1.978 | 0.8405 |
| 60 | 1F98_A | 7.148 | 0.7455 | 8.298 | 0.6118 | 7.122 | 0.7559 | 7.709 | 0.7438 | 7.838 | 0.7644 | 8.666 | 0.7675 |
| 61 | 1F9P_A | 10.286 | 0.6278 | 9.614 | 0.5811 | 6.17 | 0.6556 | 6.615 | 0.6366 | 6.74 | 0.6496 | 8.118 | 0.639 |
| 62 | 1FC3_A | 2.308 | 0.8337 | 3.135 | 0.7759 | 2.149 | 0.8459 | 2.164 | 0.8419 | 2.131 | 0.8467 | 2.132 | 0.8417 |
| 63 | 1FCA_A | 1.475 | 0.7777 | 1.6805 | 0.8233 | 1.699 | 0.7337 | 1.727 | 0.745 | 1.495 | 0.7704 | 1.644 | 0.7379 |
| 64 | 1FEX_A | 2.808 | 0.666 | 2.647 | 0.654 | 2.806 | 0.6688 | 2.366 | 0.6797 | 2.477 | 0.6721 | 2.295 | 0.6842 |
| 65 | 1FHT_A | 12.939 | 0.6536 | 12.905 | 0.5938 | 10.806 | 0.6868 | 6.163 | 0.7019 | 10.435 | 0.695 | 10.462 | 0.6825 |
| 66 | 1FJG_H | 2.919 | 0.8197 | 10.856 | 0.691 | 3.094 | 0.8374 | 2.851 | 0.8391 | 2.885 | 0.8518 | 2.781 | 0.8521 |
| 67 | 1FMB_A | 3.512 | 0.7099 | 6.144 | 0.4968 | 2.393 | 0.7845 | 3.675 | 0.7028 | 2.654 | 0.7657 | 3.367 | 0.7062 |
| 68 | 1FR0_A | 3.216 | 0.7671 | 3.049 | 0.7436 | 2.879 | 0.7717 | 3.344 | 0.7545 | 3.132 | 0.7575 | 2.951 | 0.7711 |
| 69 | 1FRD_A | 3.068 | 0.8241 | 3.7735 | 0.6368 | 2.076 | 0.828 | 2.154 | 0.8225 | 2.105 | 0.8268 | 2.049 | 0.8283 |
| 70 | 1FSP_A | 2.794 | 0.8518 | 2.9085 | 0.8492 | 2.912 | 0.8517 | 3.006 | 0.8457 | 2.741 | 0.8567 | 2.553 | 0.8528 |
| 71 | 1FSU_A | 11.221 | 0.7121 | 26.322 | 0.3467 | 6.71 | 0.8238 | 5.699 | 0.8606 | 5.961 | 0.8499 | 5.296 | 0.8582 |
| 72 | 1FW9_A | 8.849 | 0.7106 | 8.553 | 0.5444 | 8.199 | 0.7259 | 8.551 | 0.7365 | 8.304 | 0.7287 | 8.372 | 0.7284 |
| 73 | 1FY2_A | 2.536 | 0.8654 | 4.731 | 0.6942 | 2.598 | 0.861 | 14.546 | 0.3189 | 2.7 | 0.8586 | 2.568 | 0.8672 |
| 74 | 1G2R_A | 1.755 | 0.8417 | 2.3445 | 0.7702 | 1.67 | 0.8449 | 1.895 | 0.8344 | 1.61 | 0.8527 | 1.797 | 0.8476 |
| 75 | 1G8Q_A | 3.648 | 0.7067 | 5.3035 | 0.536 | 2.785 | 0.7597 | 3.67 | 0.704 | 2.605 | 0.7626 | 2.557 | 0.7667 |
| 76 | 1GGS_A | 10.083 | 0.6919 | 10.8635 | 0.6603 | 9.589 | 0.682 | 7.138 | 0.6512 | 9.957 | 0.6812 | 9.527 | 0.6827 |
| 77 | 1GME_A | 17.2 | 0.54 | 11.9795 | 0.449 | 15.464 | 0.539 | 17.133 | 0.2533 | 15.811 | 0.5377 | 16.237 | 0.54 |
| 78 | 1GPQ_B | 6.339 | 0.6294 | 6.435 | 0.5907 | 5.737 | 0.6925 | 5.707 | 0.6995 | 6.008 | 0.7016 | 5.802 | 0.6968 |
| 79 | 1GQA_A | 3.177 | 0.8152 | 6.886 | 0.71 | 2.394 | 0.8282 | 2.105 | 0.8407 | 2.201 | 0.8388 | 1.979 | 0.8491 |
| 80 | 1GVP_A | 4.919 | 0.7115 | 4.442 | 0.601 | 5.343 | 0.665 | 5.203 | 0.6755 | 5.189 | 0.6812 | 5.025 | 0.6772 |
| 81 | 1GXD_C | 15.535 | 0.5216 | 10.794 | 0.4284 | 9.729 | 0.6153 | 7.867 | 0.61 | 9.64 | 0.6365 | 7.115 | 0.6285 |
| 82 | 1GXL_A | 13.567 | 0.6072 | 9.1625 | 0.4384 | 3.297 | 0.7765 | 3.407 | 0.7699 | 3.181 | 0.7911 | 3.257 | 0.7895 |
| 83 | 1GYH_A | 5.513 | 0.7741 | 8.415 | 0.5298 | 4.876 | 0.8221 | 4.746 | 0.8277 | 4.791 | 0.825 | 4.505 | 0.8374 |
| 84 | 1H4L_D | 2.807 | 0.7763 | 3.391 | 0.7568 | 2.347 | 0.8264 | 3.012 | 0.7568 | 2.355 | 0.8258 | 2.394 | 0.8251 |
| 85 | 1H8E_H | 1.775 | 0.8344 | 11.469 | 0.6022 | 1.452 | 0.8729 | 1.462 | 0.8709 | 1.48 | 0.871 | 1.469 | 0.8745 |
| 86 | 1H9E_A | 5.969 | 0.5304 | 6.584 | 0.519 | 6.395 | 0.5318 | 6.065 | 0.5216 | 6.38 | 0.53 | 6.234 | 0.5329 |
| 87 | 1H9F_A | 5.088 | 0.5861 | 4.1985 | 0.5671 | 4.861 | 0.5826 | 4.193 | 0.5806 | 4.676 | 0.5835 | 4.523 | 0.5938 |
| 88 | 1HBG_A | 1.987 | 0.8753 | 2.812 | 0.7932 | 1.956 | 0.8822 | 2.075 | 0.8739 | 1.955 | 0.8833 | 1.935 | 0.8826 |
| 89 | 1HBX_E | 8.163 | 0.5355 | 12.16 | 0.5585 | 6.796 | 0.5553 | 8.829 | 0.5496 | 6.86 | 0.5682 | 7.581 | 0.5695 |
| 90 | 1HCD_A | 3.27 | 0.7199 | 4.239 | 0.634 | 2.958 | 0.7475 | 3.028 | 0.7346 | 2.984 | 0.7417 | 2.921 | 0.7467 |
| 91 | 1HH8_A | 14.325 | 0.7257 | 15.321 | 0.6869 | 15.591 | 0.7257 | 16.356 | 0.7211 | 15.756 | 0.7453 | 15.963 | 0.7432 |
| 92 | 1HHV_A | 5.116 | 0.6157 | 7.167 | 0.5375 | 6.981 | 0.6187 | 11.398 | 0.2556 | 6.483 | 0.6301 | 5.949 | 0.6345 |
| 93 | 1HKQ_A | 5.645 | 0.723 | 3.351 | 0.7049 | 7.473 | 0.7136 | 8.662 | 0.7027 | 8.589 | 0.7128 | 3.107 | 0.7621 |
| 94 | 1HKX_E | 6.203 | 0.6839 | 10.589 | 0.5466 | 5.716 | 0.7235 | 5.413 | 0.718 | 5.859 | 0.7229 | 5.511 | 0.727 |
| 95 | 1HL6_D | 6.672 | 0.6362 | 7.055 | 0.526 | 6.771 | 0.6396 | 5.011 | 0.6595 | 6.767 | 0.6568 | 6.549 | 0.66 |
| 96 | 1I27_A | 4.243 | 0.7187 | 6.356 | 0.6462 | 5.161 | 0.6573 | 5.422 | 0.672 | 6.433 | 0.6495 | 6.378 | 0.648 |
| 97 | 1I35_A | 4.332 | 0.6009 | 4.703 | 0.5797 | 4.179 | 0.6127 | 4.033 | 0.5968 | 4.436 | 0.5946 | 4.433 | 0.6006 |
| 98 | 1I85_A | 2.431 | 0.7802 | 4.286 | 0.6785 | 2.201 | 0.8098 | 2.222 | 0.8065 | 2.216 | 0.8046 | 2.159 | 0.809 |
| 99 | 1ID2_A | 6.822 | 0.693 | 5.696 | 0.6106 | 7.417 | 0.7132 | 7.48 | 0.7268 | 7.599 | 0.7237 | 7.271 | 0.7266 |
| 100 | 1IEZ_A | 5.83 | 0.6498 | 7.517 | 0.4917 | 4.466 | 0.7244 | 4.885 | 0.7206 | 4.533 | 0.7236 | 4.952 | 0.7212 |
| 101 | 1IJY_A | 3.936 | 0.7306 | 4.163 | 0.6543 | 2.975 | 0.7622 | 3.308 | 0.7404 | 2.63 | 0.7847 | 3.04 | 0.7538 |
| 102 | 1IOO_A | 5.303 | 0.7056 | 8.5345 | 0.4412 | 3.437 | 0.7879 | 3.365 | 0.8001 | 3.31 | 0.8055 | 3.136 | 0.8174 |
| 103 | 1IPI_A | 2.764 | 0.811 | 3.4375 | 0.744 | 2.459 | 0.829 | 2.413 | 0.826 | 2.462 | 0.834 | 2.452 | 0.8302 |

Continued on next page

Continue to last page

| No. | PDB | IddFold | | Rosetta_D | | IddFold_relax | | trRosetta_D | | IddFold_relax2 | | trRosetta | |
|---|---|---|---|---|---|---|---|---|---|---|---|---|---|
| | | RMSD | TMscore | RMSD | TMscore | RMSD | TMscore | RMSD | TMscore | RMSD | TMscore | RMSD | TMscore |
| 104 | 1IQV_A | 4.476 | 0.7833 | 5.6965 | 0.6931 | 5.008 | 0.8181 | 4.058 | 0.8235 | 3.213 | 0.849 | 3.201 | 0.8502 |
| 105 | 1IRS_A | 3.1 | 0.7791 | 4.245 | 0.6059 | 3.189 | 0.83 | 2.349 | 0.854 | 2.579 | 0.8505 | 2.144 | 0.8634 |
| 106 | 1IS7_K | 6.026 | 0.6377 | 6.4135 | 0.6142 | 4.876 | 0.7161 | 6.384 | 0.6372 | 4.78 | 0.7205 | 4.617 | 0.7285 |
| 107 | 1IUJ_B | 2.173 | 0.8703 | 2.5435 | 0.7839 | 1.871 | 0.8633 | 1.804 | 0.8678 | 1.847 | 0.8622 | 1.508 | 0.8814 |
| 108 | 1IUY_A | 9.849 | 0.7244 | 9.032 | 0.5403 | 9.8 | 0.7241 | 8.567 | 0.7094 | 9.768 | 0.7285 | 9.569 | 0.7348 |
| 109 | 1J1V_A | 0.967 | 0.9379 | 1.949 | 0.8372 | 0.979 | 0.9473 | 0.899 | 0.9417 | 0.82 | 0.9509 | 0.884 | 0.9454 |
| 110 | 1J3W_A | 2.787 | 0.8584 | 5.606 | 0.6606 | 1.98 | 0.882 | 2.03 | 0.8721 | 1.77 | 0.8892 | 1.59 | 0.8994 |
| 111 | 1J8I_A | 14.153 | 0.5433 | 18.927 | 0.5655 | 18.741 | 0.5407 | 17.79 | 0.539 | 17.881 | 0.5571 | 18.803 | 0.5651 |
| 112 | 1JC7_A | 4.795 | 0.6583 | 6.958 | 0.4309 | 4.452 | 0.6833 | 5.467 | 0.6706 | 4.386 | 0.6917 | 4.859 | 0.6875 |
| 113 | 1JEI_A | 6.306 | 0.6306 | 6.371 | 0.579 | 6.328 | 0.6104 | 6.479 | 0.6213 | 6.617 | 0.6265 | 6.623 | 0.6192 |
| 114 | 1JIW_I | 3.576 | 0.7124 | 4.304 | 0.5899 | 3.485 | 0.7192 | 3.155 | 0.7253 | 3.432 | 0.7238 | 3.264 | 0.7221 |
| 115 | 1JJ2_S | 4.075 | 0.7297 | 4.546 | 0.7313 | 3.793 | 0.7257 | 10.241 | 0.3817 | 3.829 | 0.7186 | 4.294 | 0.7167 |
| 116 | 1JLI_A | 15.002 | 0.3582 | 12.242 | 0.3002 | 13.414 | 0.3821 | 11.248 | 0.3337 | 12.51 | 0.3726 | 11.19 | 0.3391 |
| 117 | 1JMT_A | 4.372 | 0.7701 | 4.476 | 0.7243 | 4.305 | 0.7613 | 4.443 | 0.7568 | 4.253 | 0.76 | 4.147 | 0.7714 |
| 118 | 1JO0_A | 2.045 | 0.856 | 2.068 | 0.8349 | 1.941 | 0.8578 | 1.88 | 0.8634 | 1.971 | 0.8526 | 1.927 | 0.8577 |
| 119 | 1JOF_A | 6.435 | 0.801 | 7.641 | 0.6075 | 5.579 | 0.8555 | 5.434 | 0.8624 | 5.541 | 0.862 | 5.329 | 0.8745 |
| 120 | 1JOP_A | 3.443 | 0.7513 | 7.6865 | 0.5195 | 2.802 | 0.7897 | 2.936 | 0.7852 | 2.687 | 0.7991 | 2.768 | 0.7924 |
| 121 | 1JPY_Y | 18.606 | 0.5569 | 13.285 | 0.3481 | 12.4 | 0.5653 | 11.376 | 0.5605 | 11.666 | 0.6112 | 12.546 | 0.6144 |
| 122 | 1JR8_A | 1.989 | 0.8323 | 2.4595 | 0.8261 | 2.311 | 0.816 | 2.223 | 0.8203 | 2.182 | 0.8218 | 2.179 | 0.8237 |
| 123 | 1JSG_A | 7.493 | 0.5168 | 8.8995 | 0.5038 | 6.812 | 0.5521 | 13.384 | 0.2971 | 5.666 | 0.6639 | 5.945 | 0.6327 |
| 124 | 1K1Z_A | 7.306 | 0.5653 | 4.8685 | 0.5793 | 7.491 | 0.5618 | 5.626 | 0.5609 | 6.835 | 0.5614 | 8.136 | 0.5558 |
| 125 | 1K3B_A | 5.105 | 0.7039 | 5.546 | 0.6076 | 4.621 | 0.7247 | 3.419 | 0.7674 | 4.329 | 0.7298 | 3.347 | 0.7672 |
| 126 | 1K3S_A | 3.355 | 0.7096 | 4.91 | 0.5631 | 3.188 | 0.7192 | 3.186 | 0.718 | 3.094 | 0.7158 | 3.032 | 0.7252 |
| 127 | 1K5D_B | 14.936 | 0.7412 | 6.751 | 0.6284 | 13.399 | 0.7749 | 14.027 | 0.7745 | 7.963 | 0.7774 | 11.073 | 0.7915 |
| 128 | 1K73_1 | 4.571 | 0.6616 | 3.8745 | 0.6174 | 3.846 | 0.634 | 3.211 | 0.6469 | 3.614 | 0.6013 | 3.456 | 0.6226 |
| 129 | 1KA8_A | 7.377 | 0.6281 | 5.828 | 0.5444 | 4.79 | 0.5911 | 4.277 | 0.6113 | 4.778 | 0.5987 | 3.957 | 0.6607 |
| 130 | 1KN6_A | 3.564 | 0.6025 | 3.374 | 0.5994 | 3.147 | 0.6257 | 3.413 | 0.6239 | 3.244 | 0.6254 | 3.347 | 0.6212 |
| 131 | 1KOH_D | 6.771 | 0.7798 | 9.65 | 0.7532 | 5.43 | 0.7915 | 4.594 | 0.7912 | 5.611 | 0.7907 | 5.322 | 0.7975 |
| 132 | 1KPT_A | 3.706 | 0.7135 | 3.9315 | 0.6254 | 3.152 | 0.7179 | 3.174 | 0.7108 | 2.932 | 0.7325 | 2.932 | 0.7429 |
| 133 | 1KQ6_A | 10.219 | 0.6803 | 5.5 | 0.6307 | 5.074 | 0.7193 | 6.47 | 0.7118 | 5.615 | 0.7238 | 6.851 | 0.7207 |
| 134 | 1KSX_A | 3.062 | 0.7673 | 4.256 | 0.6489 | 2.783 | 0.8014 | 2.997 | 0.776 | 2.945 | 0.7883 | 3.001 | 0.7827 |
| 135 | 1KX5_D | 8.377 | 0.8009 | 7.438 | 0.7962 | 6.134 | 0.6613 | 4.993 | 0.6174 | 5.182 | 0.7783 | 5.098 | 0.7458 |
| 136 | 1L1D_B | 2.612 | 0.815 | 5.612 | 0.5137 | 2.228 | 0.8443 | 2.667 | 0.8339 | 2.214 | 0.8448 | 2.196 | 0.8474 |
| 137 | 1L2P_A | 0.726 | 0.9333 | 0.813 | 0.9208 | 0.792 | 0.9214 | 4.255 | 0.641 | 0.785 | 0.923 | 0.79 | 0.9226 |
| 138 | 1L3G_A | 7.509 | 0.5636 | 5.759 | 0.6093 | 7.597 | 0.5915 | 13.183 | 0.2635 | 7.559 | 0.5986 | 8.644 | 0.5933 |
| 139 | 1LDD_A | 0.971 | 0.9119 | 2.1675 | 0.8196 | 1.011 | 0.9096 | 1.192 | 0.8931 | 1.045 | 0.9063 | 1.131 | 0.9022 |
| 140 | 1LE2_A | 40.624 | 0.2564 | 42.29 | 0.2588 | 29.285 | 0.4295 | 10.829 | 0.3312 | 29.148 | 0.4382 | 9.472 | 0.4035 |
| 141 | 1LFU_P | 10.726 | 0.5865 | 10.941 | 0.5633 | 11.863 | 0.5962 | 10.454 | 0.5836 | 11.844 | 0.5948 | 11.726 | 0.5964 |
| 142 | 1LMI_A | 3.857 | 0.724 | 5.606 | 0.5394 | 3.289 | 0.7565 | 3.061 | 0.7653 | 3.131 | 0.7632 | 2.996 | 0.7684 |
| 143 | 1LNW_C | 4.665 | 0.8435 | 4.399 | 0.7999 | 4.474 | 0.8391 | 4.321 | 0.8157 | 4.432 | 0.8415 | 4.577 | 0.8343 |
| 144 | 1LQM_H | 11.022 | 0.4285 | 10.1 | 0.3254 | 10.346 | 0.4385 | 10.157 | 0.2617 | 10.028 | 0.433 | 7.748 | 0.5285 |
| 145 | 1LR1_B | 4.136 | 0.5162 | 3.829 | 0.5266 | 3.769 | 0.5167 | 5.186 | 0.4361 | 3.795 | 0.519 | 3.821 | 0.5156 |
| 146 | 1LWB_A | 5.501 | 0.6238 | 8.511 | 0.54 | 4.41 | 0.6508 | 4.473 | 0.6373 | 4.068 | 0.6733 | 4.265 | 0.651 |
| 147 | 1LYV_A | 4.837 | 0.7475 | 7.962 | 0.5842 | 4.178 | 0.7917 | 4.292 | 0.7851 | 4.036 | 0.8016 | 4.128 | 0.7959 |
| 148 | 1LZW_B | 4.238 | 0.8518 | 4.324 | 0.7832 | 3.667 | 0.8714 | 3.434 | 0.8648 | 3.459 | 0.8813 | 3.487 | 0.8798 |
| 149 | 1M7Y_A | 8.196 | 0.7009 | 24.5005 | 0.3409 | 4.301 | 0.8709 | 4.312 | 0.8741 | 3.733 | 0.8856 | 3.736 | 0.886 |
| 150 | 1MAI_A | 2.641 | 0.811 | 3.9845 | 0.6616 | 2.716 | 0.8381 | 2.66 | 0.8292 | 2.811 | 0.8261 | 2.656 | 0.831 |
| 151 | 1MC2_A | 8.206 | 0.691 | 7.283 | 0.6255 | 3.873 | 0.7287 | 4.422 | 0.7229 | 5.7 | 0.7081 | 6.296 | 0.7182 |
| 152 | 1MFQ_C | 8.976 | 0.6453 | 8.903 | 0.586 | 9.127 | 0.6428 | 9.529 | 0.6162 | 9.046 | 0.6432 | 8.978 | 0.6447 |
| 153 | 1MHD_A | 4.694 | 0.6183 | 5.2295 | 0.5338 | 4.002 | 0.6242 | 4.05 | 0.6392 | 3.589 | 0.6732 | 3.489 | 0.6814 |
| 154 | 1MKF_A | 32.228 | 0.2007 | 31.0145 | 0.1864 | 23.25 | 0.2156 | 20.012 | 0.2195 | 25.511 | 0.2362 | 21.651 | 0.2853 |
| 155 | 1MN8_A | 2.907 | 0.7803 | 3.802 | 0.664 | 2.673 | 0.7853 | 2.576 | 0.7849 | 2.392 | 0.7998 | 2.462 | 0.8008 |

Continued on next page

Continue to last page

| No. | PDB | IddFold | | Rosetta_D | | IddFold_relax | | trRosetta_D | | IddFold_relax2 | | trRosetta | |
|---|---|---|---|---|---|---|---|---|---|---|---|---|---|
| | | RMSD | TMscore | RMSD | TMscore | RMSD | TMscore | RMSD | TMscore | RMSD | TMscore | RMSD | TMscore |
| 156 | 1MT3_A | 4.122 | 0.8012 | 7.85 | 0.5928 | 3.504 | 0.8289 | 17.276 | 0.3027 | 3.472 | 0.8329 | 3.306 | 0.8384 |
| 157 | 1MWP_A | 5.581 | 0.6231 | 7.156 | 0.4715 | 3.515 | 0.6732 | 4.479 | 0.6281 | 3.929 | 0.6502 | 3.519 | 0.6697 |
| 158 | 1MWQ_A | 3.7 | 0.7752 | 3.149 | 0.6962 | 2.4 | 0.8324 | 2.773 | 0.8169 | 2.551 | 0.828 | 2.8 | 0.825 |
| 159 | 1N12_A | 4.25 | 0.7034 | 9.1035 | 0.4003 | 3.536 | 0.734 | 3.279 | 0.7494 | 3.447 | 0.7429 | 3.299 | 0.7471 |
| 160 | 1N13_B | 3.124 | 0.7674 | 5.313 | 0.5713 | 2.595 | 0.7904 | 7.169 | 0.4177 | 2.223 | 0.8162 | 2.544 | 0.7755 |
| 161 | 1N3G_A | 6.392 | 0.746 | 6.351 | 0.6611 | 6.371 | 0.7363 | 7.058 | 0.7219 | 6.367 | 0.7376 | 7.839 | 0.7247 |
| 162 | 1N8V_A | 1.864 | 0.8259 | 2.9675 | 0.7607 | 1.879 | 0.8225 | 1.884 | 0.8179 | 1.82 | 0.8295 | 1.767 | 0.8361 |
| 163 | 1NF6_F | 9.661 | 0.8159 | 8.3755 | 0.8282 | 7.685 | 0.839 | 5 | 0.849 | 9.123 | 0.8344 | 8.998 | 0.8376 |
| 164 | 1NGL_A | 7.876 | 0.6481 | 6.366 | 0.586 | 6.913 | 0.6737 | 17.155 | 0.2719 | 7.124 | 0.6796 | 7.816 | 0.6833 |
| 165 | 1NKZ_A | 4.563 | 0.6016 | 4.052 | 0.6076 | 4.86 | 0.5865 | 6.236 | 0.5035 | 4.887 | 0.5693 | 4.823 | 0.572 |
| 166 | 1NLQ_A | 2.666 | 0.7888 | 4.589 | 0.5555 | 2.412 | 0.8141 | 2.24 | 0.8307 | 2.242 | 0.8238 | 2.252 | 0.8351 |
| 167 | 1NOE_A | 4.495 | 0.6421 | 5.413 | 0.4291 | 4.063 | 0.69 | 3.505 | 0.6866 | 3.518 | 0.692 | 3.337 | 0.7025 |
| 168 | 1NPB_A | 7.237 | 0.6851 | 8.085 | 0.6341 | 8.657 | 0.6513 | 10.762 | 0.6258 | 10.469 | 0.6401 | 10.531 | 0.6269 |
| 169 | 1NQJ_B | 7.504 | 0.7543 | 8.079 | 0.5472 | 7.33 | 0.7479 | 7.369 | 0.7548 | 7.353 | 0.7546 | 7.299 | 0.7595 |
| 170 | 1NQZ_A | 9.867 | 0.7584 | 8.81 | 0.5038 | 8.839 | 0.7868 | 8.478 | 0.7852 | 8.554 | 0.7945 | 8.849 | 0.7919 |
| 171 | 1NR3_A | 13.204 | 0.3288 | 11.071 | 0.3447 | 19.677 | 0.3074 | 20.216 | 0.2276 | 18.714 | 0.3095 | 18.508 | 0.3103 |
| 172 | 1NRJ_B | 3.802 | 0.8053 | 5.385 | 0.6386 | 3.184 | 0.8385 | 2.717 | 0.8451 | 2.847 | 0.8427 | 2.775 | 0.8408 |
| 173 | 1NTV_A | 3.824 | 0.8052 | 5.339 | 0.5864 | 4.663 | 0.7917 | 4.711 | 0.7793 | 4.969 | 0.7913 | 4.887 | 0.7906 |
| 174 | 1NZE_A | 2.366 | 0.8805 | 4.154 | 0.8269 | 1.847 | 0.879 | 1.758 | 0.8826 | 1.654 | 0.8903 | 1.728 | 0.8887 |
| 175 | 1O0G_A | 4.109 | 0.6659 | 5.457 | 0.5329 | 4.031 | 0.6936 | 4.486 | 0.6678 | 3.931 | 0.6923 | 3.895 | 0.6991 |
| 176 | 1O9G_A | 10.633 | 0.6359 | 8.755 | 0.5493 | 8.991 | 0.6753 | 11.613 | 0.6476 | 8.759 | 0.668 | 11.099 | 0.6513 |
| 177 | 1OA8_D | 13.897 | 0.3474 | 12.5265 | 0.3083 | 13.5 | 0.3306 | 13.596 | 0.3432 | 13.653 | 0.3312 | 14.411 | 0.3419 |
| 178 | 1OFT_A | 2.659 | 0.7936 | 3.0435 | 0.7554 | 2.631 | 0.7956 | 2.464 | 0.8147 | 2.485 | 0.8058 | 2.456 | 0.812 |
| 179 | 1OJG_A | 4.043 | 0.6318 | 4.6725 | 0.6242 | 3.679 | 0.6556 | 3.684 | 0.6531 | 3.635 | 0.6636 | 3.628 | 0.6625 |
| 180 | 1OK0_A | 2.852 | 0.712 | 3.755 | 0.5125 | 2.917 | 0.7009 | 2.994 | 0.7028 | 2.983 | 0.7054 | 2.76 | 0.7254 |
| 181 | 1OOF_A | 2.05 | 0.8337 | 4.1035 | 0.8038 | 1.921 | 0.8513 | 1.93 | 0.8476 | 1.96 | 0.8479 | 1.999 | 0.8439 |
| 182 | 1OPO_C | 17.678 | 0.2825 | 33.382 | 0.1414 | 15.809 | 0.302 | 15.739 | 0.2857 | 15.021 | 0.33 | 9.588 | 0.569 |
| 183 | 1ORQ_C | 46.591 | 0.4208 | 26.917 | 0.3377 | 28.922 | 0.4216 | 26.641 | 0.3972 | 31.516 | 0.4267 | 31.294 | 0.4277 |
| 184 | 1ORY_A | 2.603 | 0.8394 | 3.475 | 0.8062 | 2.301 | 0.846 | 2.083 | 0.8388 | 2.398 | 0.8474 | 2.407 | 0.8453 |
| 185 | 1OX7_A | 3.329 | 0.8099 | 4.019 | 0.6811 | 2.87 | 0.8087 | 2.959 | 0.8095 | 2.814 | 0.8132 | 2.758 | 0.8301 |
| 186 | 1OZ9_A | 6.745 | 0.7552 | 6.0315 | 0.6253 | 2.189 | 0.8409 | 2.394 | 0.8369 | 2.45 | 0.8408 | 2.459 | 0.8441 |
| 187 | 1P9O_B | 8.166 | 0.7193 | 8.948 | 0.5775 | 4.692 | 0.798 | 6.194 | 0.779 | 4.369 | 0.8087 | 4.694 | 0.8085 |
| 188 | 1PD6_A | 5.03 | 0.6756 | 4.982 | 0.651 | 4.943 | 0.6852 | 4.513 | 0.6734 | 4.804 | 0.6885 | 4.854 | 0.6785 |
| 189 | 1PFS_A | 5.327 | 0.5636 | 5.026 | 0.5711 | 5.044 | 0.5688 | 4.883 | 0.5798 | 4.876 | 0.5741 | 4.674 | 0.5889 |
| 190 | 1PGV_A | 3.278 | 0.7803 | 4.441 | 0.7233 | 3.105 | 0.7906 | 3.167 | 0.7845 | 3.282 | 0.778 | 3.33 | 0.7745 |
| 191 | 1PIH_A | 4.658 | 0.6497 | 5.8185 | 0.4601 | 4.457 | 0.665 | 5.094 | 0.6692 | 4.701 | 0.6802 | 4.688 | 0.6788 |
| 192 | 1PMS_A | 8.985 | 0.6074 | 12.409 | 0.5383 | 8.51 | 0.6186 | 8.26 | 0.6273 | 8.575 | 0.6224 | 8.518 | 0.6238 |
| 193 | 1PSR_A | 3.613 | 0.7342 | 3.81 | 0.6469 | 3.68 | 0.689 | 6.135 | 0.6885 | 3.886 | 0.6978 | 4.326 | 0.7008 |
| 194 | 1PXW_A | 6.206 | 0.7383 | 6.548 | 0.6023 | 6.475 | 0.7592 | 4.405 | 0.7704 | 6.947 | 0.7515 | 6.47 | 0.7539 |
| 195 | 1PZW_A | 2.55 | 0.724 | 3.4745 | 0.6389 | 2.327 | 0.7651 | 2.502 | 0.7449 | 2.504 | 0.757 | 2.384 | 0.7576 |
| 196 | 1QFT_A | 4.995 | 0.7607 | 6.124 | 0.6001 | 4.809 | 0.7814 | 3.954 | 0.7937 | 4.25 | 0.7904 | 4.237 | 0.7881 |
| 197 | 1QFW_B | 14.524 | 0.5411 | 12.191 | 0.3383 | 10.887 | 0.4786 | 8.155 | 0.465 | 11.025 | 0.5051 | 6.476 | 0.5979 |
| 198 | 1QMA_A | 1.61 | 0.8983 | 3.2015 | 0.7753 | 1.637 | 0.9095 | 1.833 | 0.9021 | 1.698 | 0.9081 | 1.748 | 0.9082 |
| 199 | 1QWT_A | 14.196 | 0.61 | 17.3535 | 0.3635 | 10.075 | 0.6325 | 8.373 | 0.6493 | 7.711 | 0.6562 | 7.573 | 0.6756 |
| 200 | 1QZG_A | 6.075 | 0.6695 | 6.414 | 0.5369 | 3.367 | 0.7755 | 3.915 | 0.7614 | 3.187 | 0.7956 | 3.261 | 0.7931 |
| 201 | 1R5T_A | 3.927 | 0.7891 | 4.612 | 0.6741 | 3.396 | 0.8207 | 3.267 | 0.8326 | 3.502 | 0.8343 | 2.961 | 0.8508 |
| 202 | 1R5Z_A | 4.093 | 0.7983 | 11.1675 | 0.4687 | 3.3 | 0.8355 | 3.228 | 0.8401 | 3.28 | 0.8384 | 3.208 | 0.8443 |
| 203 | 1R6R_A | 8.836 | 0.564 | 6.208 | 0.5075 | 8.517 | 0.5384 | 11.725 | 0.3707 | 8.346 | 0.5507 | 8.327 | 0.5457 |
| 204 | 1RHX_A | 4.245 | 0.6275 | 4.509 | 0.5439 | 4.205 | 0.6326 | 4.341 | 0.6141 | 4.229 | 0.6345 | 4.202 | 0.6357 |
| 205 | 1RKX_A | 4.596 | 0.8455 | 10.045 | 0.5244 | 2.617 | 0.9123 | 2.107 | 0.9297 | 2.155 | 0.9312 | 2.104 | 0.9355 |
| 206 | 1ROW_A | 2.402 | 0.8067 | 3.7335 | 0.5879 | 1.616 | 0.8622 | 1.659 | 0.8566 | 1.651 | 0.8548 | 1.563 | 0.8688 |
| 207 | 1RTU_A | 6.875 | 0.7085 | 5.7755 | 0.5886 | 3.421 | 0.7474 | 3.082 | 0.7541 | 3.958 | 0.7386 | 2.921 | 0.7526 |

Continued on next page

Continue to last page

| No. | PDB | IddFold | | Rosetta_D | | IddFold_relax | | trRosetta_D | | IddFold_relax2 | | trRosetta | |
|---|---|---|---|---|---|---|---|---|---|---|---|---|---|
| | | RMSD | TMscore | RMSD | TMscore | RMSD | TMscore | RMSD | TMscore | RMSD | TMscore | RMSD | TMscore |
| 208 | 1RZ2_A | 17.939 | 0.7443 | 14.138 | 0.4752 | 4.21 | 0.7989 | 4.05 | 0.81 | 3.977 | 0.8089 | 4.3 | 0.8021 |
| 209 | 1RZ3_A | 5.222 | 0.7523 | 6.502 | 0.6098 | 2.943 | 0.849 | 3.324 | 0.8385 | 2.842 | 0.8575 | 3.201 | 0.8398 |
| 210 | 1S2D_A | 7.77 | 0.7204 | 7.602 | 0.6374 | 4.917 | 0.7415 | 3.827 | 0.7685 | 4.83 | 0.7517 | 4.772 | 0.7606 |
| 211 | 1S3J_A | 2.87 | 0.7756 | 3.066 | 0.7445 | 2.765 | 0.787 | 3.054 | 0.7739 | 2.645 | 0.7925 | 2.83 | 0.7932 |
| 212 | 1S56_B | 8.14 | 0.7794 | 8.328 | 0.6716 | 7.452 | 0.7935 | 8.442 | 0.781 | 8.124 | 0.8009 | 8.44 | 0.803 |
| 213 | 1S7O_C | 10.19 | 0.5764 | 10.894 | 0.5517 | 9.822 | 0.5826 | 9.723 | 0.5649 | 9.875 | 0.5754 | 2.941 | 0.8071 |
| 214 | 1S7Z_A | 5.71 | 0.6602 | 6.303 | 0.5867 | 5.289 | 0.6748 | 5.145 | 0.6354 | 4.941 | 0.6645 | 4.974 | 0.6745 |
| 215 | 1SAU_A | 3.135 | 0.7192 | 3.604 | 0.7099 | 2.938 | 0.7312 | 3.049 | 0.7191 | 2.963 | 0.7261 | 2.951 | 0.7215 |
| 216 | 1SG4_B | 9.24 | 0.771 | 10.0915 | 0.6092 | 2.412 | 0.9016 | 1.965 | 0.9179 | 1.824 | 0.9263 | 1.723 | 0.9312 |
| 217 | 1SMP_I | 2.759 | 0.7488 | 4.272 | 0.6371 | 2.639 | 0.7673 | 2.653 | 0.7709 | 2.527 | 0.7732 | 2.592 | 0.7644 |
| 218 | 1SPP_B | 4.142 | 0.7343 | 5.208 | 0.5448 | 2.757 | 0.772 | 11.199 | 0.309 | 2.781 | 0.769 | 2.656 | 0.7752 |
| 219 | 1SR8_A | 15.778 | 0.4934 | 10.2555 | 0.4527 | 15.328 | 0.5474 | 10.078 | 0.6139 | 15.917 | 0.5487 | 5.766 | 0.662 |
| 220 | 1STM_A | 8.337 | 0.5251 | 9.6525 | 0.3353 | 4.914 | 0.6271 | 11.973 | 0.3216 | 4.604 | 0.6644 | 4.294 | 0.6667 |
| 221 | 1SVJ_A | 2.768 | 0.8006 | 4.509 | 0.6813 | 2.387 | 0.8355 | 2.207 | 0.8487 | 2.233 | 0.8375 | 2.135 | 0.8444 |
| 222 | 1TAF_A | 1.2 | 0.8925 | 1.298 | 0.8647 | 1.319 | 0.8561 | 1.808 | 0.8275 | 1.272 | 0.8681 | 1.257 | 0.8665 |
| 223 | 1TEO_A | 8.5 | 0.6745 | 8.599 | 0.6008 | 7.513 | 0.7358 | 8.329 | 0.7267 | 7.322 | 0.734 | 8.065 | 0.7513 |
| 224 | 1TJF_B | 6.308 | 0.8256 | 5.836 | 0.7386 | 4.244 | 0.866 | 6.759 | 0.8156 | 4.056 | 0.8873 | 4.242 | 0.8874 |
| 225 | 1TLJ_A | 3.668 | 0.7562 | 6.563 | 0.4823 | 3.243 | 0.808 | 2.934 | 0.8209 | 3.036 | 0.827 | 2.887 | 0.8237 |
| 226 | 1TUL_A | 3.512 | 0.647 | 7.555 | 0.5542 | 3.139 | 0.6673 | 11.198 | 0.2698 | 3.011 | 0.7002 | 2.986 | 0.7074 |
| 227 | 1TWU_A | 5.266 | 0.7352 | 4.162 | 0.6959 | 5.607 | 0.6918 | 13.044 | 0.3155 | 5.585 | 0.6895 | 5.909 | 0.7015 |
| 228 | 1TYG_B | 1.821 | 0.8061 | 2.035 | 0.7748 | 2.338 | 0.8104 | 1.752 | 0.8041 | 2.127 | 0.8177 | 1.605 | 0.8257 |
| 229 | 1TZ0_A | 3.806 | 0.6926 | 4.41 | 0.6109 | 3.772 | 0.6928 | 3.37 | 0.7121 | 3.633 | 0.7027 | 3.744 | 0.7051 |
| 230 | 1U84_A | 1.613 | 0.8663 | 1.687 | 0.8655 | 1.601 | 0.8701 | 1.703 | 0.854 | 1.612 | 0.8707 | 1.603 | 0.8734 |
| 231 | 1UDD_A | 2.122 | 0.9016 | 3.471 | 0.791 | 1.583 | 0.9351 | 1.563 | 0.9348 | 1.53 | 0.9377 | 1.571 | 0.9382 |
| 232 | 1UFB_A | 3.993 | 0.8315 | 3.8175 | 0.7443 | 2.402 | 0.8697 | 11.223 | 0.35 | 2.081 | 0.8787 | 1.698 | 0.8911 |
| 233 | 1UG4_A | 5.694 | 0.6067 | 4.986 | 0.5945 | 4.622 | 0.6016 | 6.839 | 0.346 | 4.4 | 0.6221 | 4.111 | 0.6113 |
| 234 | 1UGI_A | 4.56 | 0.5949 | 7.178 | 0.5693 | 4.063 | 0.579 | 6.715 | 0.4586 | 3.911 | 0.5747 | 7.909 | 0.5208 |
| 235 | 1UNG_D | 2.914 | 0.7521 | 3.833 | 0.7085 | 2.822 | 0.7857 | 3.192 | 0.7659 | 2.848 | 0.78 | 2.934 | 0.7743 |
| 236 | 1USL_C | 3.326 | 0.8618 | 5.897 | 0.7574 | 5.901 | 0.8574 | 6.876 | 0.8507 | 7.099 | 0.8455 | 6.835 | 0.8517 |
| 237 | 1V74_A | 4.629 | 0.7314 | 3.7935 | 0.6889 | 4.963 | 0.7041 | 4.765 | 0.6975 | 4.05 | 0.7224 | 5.336 | 0.7103 |
| 238 | 1VCC_A | 5.068 | 0.5815 | 4.231 | 0.5576 | 5.451 | 0.5742 | 5.525 | 0.5273 | 5.242 | 0.5617 | 5.237 | 0.5387 |
| 239 | 1VCY_A | 5.786 | 0.6833 | 7.927 | 0.4281 | 3.87 | 0.7461 | 4.382 | 0.7365 | 3.617 | 0.7644 | 3.267 | 0.7902 |
| 240 | 1VD0_A | 11.802 | 0.6623 | 11.0115 | 0.3634 | 8.928 | 0.6687 | 8.996 | 0.6751 | 8.625 | 0.6675 | 8.605 | 0.6859 |
| 241 | 1VDW_A | 4.738 | 0.8043 | 8.253 | 0.5947 | 3.292 | 0.8524 | 2.987 | 0.866 | 2.957 | 0.8699 | 2.942 | 0.8705 |
| 242 | 1VHG_A | 12.265 | 0.6605 | 9.5735 | 0.498 | 12.398 | 0.6782 | 13.413 | 0.3763 | 12.499 | 0.6808 | 12.484 | 0.6825 |
| 243 | 1VKE_E | 5.257 | 0.7632 | 3.018 | 0.7957 | 2.064 | 0.8369 | 2.466 | 0.8279 | 1.89 | 0.8593 | 1.807 | 0.8653 |
| 244 | 1VLL_A | 3.357 | 0.8612 | 18.5585 | 0.4872 | 2.094 | 0.9324 | 1.988 | 0.9314 | 1.911 | 0.935 | 1.845 | 0.9374 |
| 245 | 1VTK_A | 18.284 | 0.467 | 24.759 | 0.2102 | 19.997 | 0.5419 | 21.186 | 0.2737 | 19.048 | 0.5511 | 19.219 | 0.5519 |
| 246 | 1VYI_A | 24.498 | 0.3072 | 18.546 | 0.2559 | 8.319 | 0.3509 | 12.305 | 0.2883 | 11.499 | 0.2926 | 11.216 | 0.291 |
| 247 | 1VYX_A | 5.199 | 0.4974 | 5.159 | 0.349 | 4.82 | 0.5301 | 5.097 | 0.5072 | 4.78 | 0.5322 | 4.546 | 0.5207 |
| 248 | 1W1W_E | 2.188 | 0.7929 | 2.1615 | 0.7758 | 2.111 | 0.786 | 2.12 | 0.783 | 1.94 | 0.797 | 1.871 | 0.8161 |
| 249 | 1W53_A | 2.526 | 0.7678 | 3.133 | 0.7336 | 3.92 | 0.7078 | 2.246 | 0.7886 | 3.043 | 0.7374 | 3.064 | 0.732 |
| 250 | 1WER_A | 3.379 | 0.8686 | 8.923 | 0.4703 | 2.36 | 0.9185 | 2.39 | 0.9135 | 2.276 | 0.9158 | 2.297 | 0.9153 |
| 251 | 1WJ8_A | 1.477 | 0.8961 | 3.148 | 0.8285 | 1.406 | 0.9042 | 1.511 | 0.8944 | 1.443 | 0.9017 | 1.462 | 0.8978 |
| 252 | 1WKR_A | 5.491 | 0.8006 | 15.0065 | 0.429 | 4.053 | 0.855 | 18.098 | 0.3251 | 4.149 | 0.8588 | 3.792 | 0.8761 |
| 253 | 1WLQ_C | 7.107 | 0.6248 | 8.413 | 0.5271 | 8.399 | 0.6874 | 8.096 | 0.6718 | 8.367 | 0.6978 | 6.685 | 0.691 |
| 254 | 1WMH_B | 1.395 | 0.8786 | 3.08 | 0.7056 | 1.386 | 0.8814 | 1.404 | 0.8691 | 1.285 | 0.8898 | 1.323 | 0.8857 |
| 255 | 1XJA_C | 4.716 | 0.7686 | 3.852 | 0.723 | 4.227 | 0.7617 | 14.691 | 0.307 | 4.21 | 0.7738 | 4.37 | 0.766 |
| 256 | 1Y14_A | 9.251 | 0.529 | 6.708 | 0.5907 | 7.163 | 0.6214 | 7.605 | 0.5421 | 6.263 | 0.6402 | 6.555 | 0.6408 |
| 257 | 1Y1X_A | 7.883 | 0.6051 | 6.067 | 0.6266 | 7.167 | 0.6362 | 14.187 | 0.2971 | 8.041 | 0.6232 | 7.658 | 0.6287 |
| 258 | 1YG2_A | 20.152 | 0.4728 | 14.664 | 0.4291 | 17.697 | 0.4692 | 17.784 | 0.469 | 17.84 | 0.4635 | 17.797 | 0.468 |
| 259 | 1YZH_B | 2.226 | 0.8944 | 3.438 | 0.7885 | 2.155 | 0.8976 | 2.098 | 0.9006 | 2.189 | 0.8997 | 2.102 | 0.9027 |

Continued on next page

Continue to last page

| No. | PDB | IddFold | | Rosetta_D | | IddFold_relax | | trRosetta_D | | IddFold_relax2 | | trRosetta | |
|---|---|---|---|---|---|---|---|---|---|---|---|---|---|
| | | RMSD | TMscore | RMSD | TMscore | RMSD | TMscore | RMSD | TMscore | RMSD | TMscore | RMSD | TMscore |
| 260 | 1Z8R_A | 12.6 | 0.4465 | 11.185 | 0.3725 | 8.948 | 0.4881 | 14.407 | 0.2742 | 9.46 | 0.494 | 9.152 | 0.4881 |
| 261 | 1ZC9_A | 5.351 | 0.7992 | 23.6695 | 0.3518 | 3.384 | 0.8793 | 3.029 | 0.8955 | 3.183 | 0.8888 | 2.955 | 0.8967 |
| 262 | 2A5Y_A | 3.486 | 0.7773 | 3.847 | 0.7372 | 3.341 | 0.8268 | 2.934 | 0.8247 | 3.349 | 0.8207 | 2.658 | 0.8447 |
| 263 | 2A9U_B | 9.314 | 0.7869 | 8.664 | 0.7649 | 10.934 | 0.7873 | 10.853 | 0.7748 | 9.933 | 0.7996 | 9.783 | 0.7964 |
| 264 | 2AAM_B | 5.255 | 0.8282 | 6.698 | 0.6929 | 3.301 | 0.8663 | 3.001 | 0.8758 | 3.184 | 0.8776 | 3.168 | 0.8705 |
| 265 | 2ACY_A | 2.563 | 0.8045 | 3.253 | 0.6825 | 2.31 | 0.8257 | 2.844 | 0.8086 | 2.345 | 0.8191 | 2.619 | 0.8214 |
| 266 | 2AEN_A | 6.328 | 0.6581 | 10.536 | 0.3789 | 5.21 | 0.6711 | 13.148 | 0.3672 | 5.373 | 0.6691 | 4.43 | 0.6652 |
| 267 | 2APN_A | 7.034 | 0.6547 | 6.017 | 0.6146 | 6.924 | 0.6658 | 11.63 | 0.2862 | 7.076 | 0.665 | 6.548 | 0.6728 |
| 268 | 2AQ0_A | 7.995 | 0.6448 | 7.636 | 0.6497 | 8.217 | 0.6369 | 7.567 | 0.6325 | 8.292 | 0.6379 | 8.577 | 0.6321 |
| 269 | 2AQS_A | 7.015 | 0.6835 | 8.008 | 0.5307 | 4.354 | 0.7571 | 3.838 | 0.8053 | 3.685 | 0.7942 | 3.39 | 0.8218 |
| 270 | 2AQX_A | 10.67 | 0.6792 | 21.177 | 0.4701 | 9.145 | 0.7151 | 19.996 | 0.2978 | 8.102 | 0.7799 | 8.373 | 0.7729 |
| 271 | 2BL7_A | 1.577 | 0.8876 | 1.7665 | 0.8949 | 1.641 | 0.8879 | 1.621 | 0.8724 | 1.694 | 0.8837 | 1.602 | 0.8853 |
| 272 | 2BS2_C | 8.135 | 0.7224 | 7.559 | 0.5904 | 5.727 | 0.761 | 5.54 | 0.7497 | 5.245 | 0.7635 | 4.754 | 0.7778 |
| 273 | 2BSE_A | 3.97 | 0.6494 | 8.0715 | 0.5511 | 4.14 | 0.674 | 10.438 | 0.303 | 3.239 | 0.7063 | 3.028 | 0.7079 |
| 274 | 2BT9_A | 2.038 | 0.8098 | 2.852 | 0.7229 | 1.901 | 0.8192 | 8.849 | 0.3445 | 1.952 | 0.8165 | 1.907 | 0.822 |
| 275 | 2BTD_A | 3.958 | 0.8642 | 3.651 | 0.767 | 1.958 | 0.9132 | 1.794 | 0.9208 | 1.825 | 0.9228 | 1.655 | 0.9318 |
| 276 | 2BWJ_A | 5.751 | 0.7659 | 4.889 | 0.6707 | 5.727 | 0.7587 | 5.703 | 0.7548 | 5.842 | 0.7628 | 5.528 | 0.7677 |
| 277 | 2BYK_D | 1.895 | 0.8422 | 1.8745 | 0.8864 | 1.77 | 0.8545 | 2.063 | 0.8093 | 1.703 | 0.8764 | 1.773 | 0.8588 |
| 278 | 2C2F_A | 8.149 | 0.7993 | 6.351 | 0.7325 | 5.236 | 0.8305 | 5.545 | 0.8243 | 5.748 | 0.8248 | 6.066 | 0.8232 |
| 279 | 2C4W_A | 3.918 | 0.801 | 4.65 | 0.6993 | 2.933 | 0.8301 | 3.829 | 0.825 | 3.06 | 0.8322 | 4.26 | 0.8353 |
| 280 | 2CDP_A | 3.259 | 0.7775 | 5.683 | 0.5393 | 2.733 | 0.7927 | 2.774 | 0.7966 | 2.69 | 0.7989 | 2.68 | 0.7998 |
| 281 | 2CEY_A | 4.759 | 0.7376 | 9.31 | 0.5112 | 2.833 | 0.8534 | 2.904 | 0.8507 | 2.701 | 0.8654 | 2.726 | 0.8633 |
| 282 | 2CMX_A | 2.205 | 0.7378 | 2.3005 | 0.749 | 2.027 | 0.7636 | 1.899 | 0.7478 | 2.112 | 0.7492 | 2.133 | 0.748 |
| 283 | 2CO3_B | 5.463 | 0.6474 | 8.253 | 0.4391 | 5.657 | 0.6662 | 5.567 | 0.6806 | 4.912 | 0.6989 | 4.856 | 0.6877 |
| 284 | 2CWP_A | 5.136 | 0.7584 | 5.636 | 0.6094 | 4.689 | 0.7842 | 4.749 | 0.7957 | 4.922 | 0.8004 | 5.013 | 0.7959 |
| 285 | 2CZV_D | 7.656 | 0.7887 | 6.851 | 0.652 | 8.568 | 0.8146 | 8.473 | 0.8058 | 8.565 | 0.8029 | 8.424 | 0.8075 |
| 286 | 2D0P_B | 2.489 | 0.8122 | 5.9275 | 0.7042 | 2.66 | 0.8226 | 3.045 | 0.8271 | 2.867 | 0.8255 | 2.716 | 0.8321 |
| 287 | 2D5R_B | 2.981 | 0.8181 | 3.459 | 0.724 | 2.4 | 0.8469 | 2.638 | 0.8191 | 2.17 | 0.8548 | 1.976 | 0.8543 |
| 288 | 2DDS_A | 11.689 | 0.5852 | 26.915 | 0.4252 | 9.987 | 0.6657 | 20.225 | 0.2806 | 10.31 | 0.6677 | 9.852 | 0.6724 |
| 289 | 2EWC_B | 3.023 | 0.8101 | 7.73 | 0.6298 | 3.576 | 0.7988 | 3.387 | 0.8033 | 3.418 | 0.806 | 3.495 | 0.8068 |
| 290 | 2F22_B | 3.647 | 0.7746 | 5.1115 | 0.6545 | 3.01 | 0.8254 | 3.147 | 0.8282 | 3.013 | 0.826 | 2.94 | 0.83 |
| 291 | 2FA5_B | 3.743 | 0.7928 | 3.8615 | 0.7601 | 3.502 | 0.7928 | 4.075 | 0.7717 | 3.571 | 0.7667 | 3.096 | 0.7761 |
| 292 | 2FKB_C | 3.692 | 0.7813 | 6.874 | 0.5031 | 2.434 | 0.8493 | 2.366 | 0.85 | 2.484 | 0.8491 | 2.676 | 0.843 |
| 293 | 2GBJ_B | 4.444 | 0.7976 | 3.439 | 0.7134 | 3.954 | 0.8172 | 3.126 | 0.812 | 3.857 | 0.8107 | 2.649 | 0.8266 |
| 294 | 2GJ3_A | 2.629 | 0.8448 | 3.7415 | 0.7315 | 2.849 | 0.8335 | 2.768 | 0.8438 | 2.683 | 0.8478 | 2.724 | 0.8457 |
| 295 | 2GKC_A | 5.86 | 0.6102 | 6.261 | 0.4589 | 5.089 | 0.6265 | 4.787 | 0.6315 | 4.649 | 0.6319 | 4.838 | 0.63 |
| 296 | 2H30_A | 4.776 | 0.7223 | 4.644 | 0.6835 | 4.337 | 0.7232 | 5.315 | 0.7086 | 4.781 | 0.7196 | 5.393 | 0.7211 |
| 297 | 2H8E_A | 2.803 | 0.7753 | 4.4875 | 0.5686 | 2.957 | 0.769 | 2.766 | 0.7773 | 2.92 | 0.7692 | 2.656 | 0.784 |
| 298 | 2HI3_A | 6.811 | 0.6325 | 5.721 | 0.6521 | 6.491 | 0.6595 | 6.589 | 0.6413 | 6.506 | 0.6708 | 6.167 | 0.657 |
| 299 | 2HQ7_B | 2.952 | 0.7826 | 4.211 | 0.6873 | 2.924 | 0.8004 | 2.82 | 0.7929 | 2.789 | 0.811 | 2.504 | 0.8169 |
| 300 | 2HYB_A | 3.531 | 0.7659 | 4.908 | 0.6394 | 2.961 | 0.8068 | 2.519 | 0.8211 | 2.883 | 0.8249 | 2.344 | 0.8293 |
| 301 | 2ICT_A | 4.924 | 0.7525 | 4.438 | 0.7625 | 4.857 | 0.7551 | 4.516 | 0.7424 | 4.494 | 0.7477 | 4.17 | 0.7481 |
| 302 | 2IEE_A | 6.962 | 0.6424 | 10.831 | 0.5028 | 4.361 | 0.7853 | 3.502 | 0.8011 | 3.295 | 0.8147 | 3.394 | 0.8058 |
| 303 | 2IGS_B | 27.086 | 0.2463 | 17.8845 | 0.2548 | 17.679 | 0.2703 | 16.074 | 0.2812 | 17.253 | 0.3288 | 16.751 | 0.2736 |
| 304 | 2J4B_B | 2.215 | 0.8182 | 3.2665 | 0.7286 | 2.19 | 0.8118 | 2.127 | 0.8198 | 2.05 | 0.83 | 2.142 | 0.8182 |
| 305 | 2J4H_B | 5.615 | 0.7421 | 10.417 | 0.4686 | 3.356 | 0.801 | 5.561 | 0.7805 | 3.283 | 0.8042 | 3.305 | 0.8018 |
| 306 | 2J5S_A | 5.335 | 0.786 | 7.593 | 0.5719 | 2.401 | 0.8902 | 2.493 | 0.8809 | 2.361 | 0.8905 | 2.117 | 0.9008 |
| 307 | 2J6Z_A | 2.457 | 0.761 | 2.452 | 0.7443 | 3.745 | 0.699 | 4.175 | 0.6971 | 3.381 | 0.7098 | 3.398 | 0.7055 |
| 308 | 2JC5_A | 4.977 | 0.7706 | 5.457 | 0.636 | 2.992 | 0.8642 | 2.077 | 0.9173 | 2.181 | 0.9077 | 1.99 | 0.9188 |
| 309 | 2JLP_D | 5.696 | 0.649 | 7.763 | 0.4516 | 4.861 | 0.7083 | 13.17 | 0.3108 | 4.737 | 0.7148 | 4.79 | 0.7092 |
| 310 | 2JP3_A | 10.65 | 0.4007 | 12.059 | 0.397 | 11.362 | 0.4149 | 10.04 | 0.3909 | 10.879 | 0.4156 | 9.986 | 0.3655 |
| 311 | 2K9X_A | 4.483 | 0.6957 | 4.906 | 0.5231 | 4.287 | 0.7093 | 4.055 | 0.7004 | 4.202 | 0.7182 | 4.245 | 0.7148 |

Continued on next page

Continue to last page

| No. | PDB | IddFold | | Rosetta_D | | IddFold_relax | | trRosetta_D | | IddFold_relax2 | | trRosetta | |
|---|---|---|---|---|---|---|---|---|---|---|---|---|---|
| | | RMSD | TMscore | RMSD | TMscore | RMSD | TMscore | RMSD | TMscore | RMSD | TMscore | RMSD | TMscore |
| 312 | 2KBW_A | 3.765 | 0.7867 | 3.983 | 0.7562 | 3.257 | 0.8258 | 3.377 | 0.8132 | 3.233 | 0.8266 | 3.103 | 0.823 |
| 313 | 2L5P_A | 4.41 | 0.7618 | 4.056 | 0.7024 | 3.669 | 0.7931 | 3.559 | 0.7951 | 3.715 | 0.7889 | 3.637 | 0.7899 |
| 314 | 2L74_A | 5.364 | 0.7137 | 4.4195 | 0.6692 | 5.6 | 0.7083 | 5.858 | 0.6964 | 5.397 | 0.7164 | 5.453 | 0.7128 |
| 315 | 2LKP_A | 12.147 | 0.6292 | 11.729 | 0.6154 | 11.644 | 0.6076 | 11.98 | 0.607 | 11.464 | 0.6153 | 11.823 | 0.6195 |
| 316 | 2LRB_A | 4.134 | 0.7202 | 5.5505 | 0.5862 | 3.997 | 0.7253 | 14.855 | 0.295 | 3.837 | 0.7296 | 3.972 | 0.7246 |
| 317 | 2LWP_A | 9.716 | 0.5339 | 8.499 | 0.4938 | 8.836 | 0.5458 | 7.16 | 0.545 | 8.696 | 0.5495 | 7.018 | 0.5544 |
| 318 | 2NAZ_A | 4.528 | 0.6872 | 3.8545 | 0.6768 | 4.48 | 0.6824 | 4.237 | 0.6856 | 4.327 | 0.685 | 4.391 | 0.6834 |
| 319 | 2NCM_A | 1.504 | 0.8749 | 3.102 | 0.7477 | 1.419 | 0.8986 | 10.233 | 0.3173 | 1.512 | 0.8921 | 1.564 | 0.885 |
| 320 | 2NDP_A | 6.145 | 0.5449 | 7.494 | 0.469 | 6.514 | 0.5876 | 13.352 | 0.4281 | 6.599 | 0.5992 | 7.159 | 0.5883 |
| 321 | 2NS9_B | 5.279 | 0.7852 | 4.533 | 0.6961 | 4.535 | 0.8329 | 4.609 | 0.8335 | 4.637 | 0.8332 | 3.504 | 0.8408 |
| 322 | 2NZY_A | 23.549 | 0.37 | 23.2915 | 0.2537 | 19.323 | 0.4484 | 13.072 | 0.5145 | 13.664 | 0.5024 | 7.687 | 0.6061 |
| 323 | 2O2Y_C | 5.971 | 0.7517 | 8.745 | 0.5618 | 4.043 | 0.8435 | 18.324 | 0.3223 | 3.782 | 0.8584 | 3.879 | 0.853 |
| 324 | 2O70_F | 2.432 | 0.823 | 4.815 | 0.6864 | 2.085 | 0.8593 | 2.247 | 0.8444 | 2.099 | 0.8591 | 2.148 | 0.8537 |
| 325 | 2ODM_B | 2.003 | 0.8492 | 2.1335 | 0.8535 | 1.904 | 0.8521 | 2.106 | 0.8149 | 1.517 | 0.8809 | 1.826 | 0.868 |
| 326 | 2P7L_A | 4.17 | 0.7597 | 3.851 | 0.7372 | 4.494 | 0.7626 | 4.898 | 0.7507 | 4.372 | 0.758 | 4.848 | 0.7571 |
| 327 | 2PI2_F | 3.813 | 0.7557 | 3.3025 | 0.7278 | 2.83 | 0.7966 | 2.726 | 0.7888 | 2.683 | 0.8094 | 2.747 | 0.8041 |
| 328 | 2PK3_A | 4.814 | 0.8153 | 8.933 | 0.5671 | 1.96 | 0.9308 | 1.881 | 0.9353 | 2.01 | 0.9302 | 1.921 | 0.9351 |
| 329 | 2PPV_A | 6.116 | 0.765 | 8.09 | 0.5614 | 2.936 | 0.9002 | 2.719 | 0.9012 | 2.67 | 0.9033 | 2.407 | 0.9035 |
| 330 | 2PYB_A | 3.074 | 0.8572 | 4.058 | 0.7722 | 3.147 | 0.8478 | 3.103 | 0.8462 | 3.07 | 0.8498 | 3.058 | 0.8477 |
| 331 | 2Q2H_A | 7.44 | 0.6884 | 8.692 | 0.5146 | 7.795 | 0.6965 | 6.412 | 0.6987 | 9.005 | 0.6913 | 8.575 | 0.687 |
| 332 | 2QTT_A | 9.197 | 0.6718 | 7.222 | 0.5333 | 4.47 | 0.8217 | 4 | 0.8212 | 4.114 | 0.8287 | 3.645 | 0.8299 |
| 333 | 2QVG_A | 2.921 | 0.8213 | 2.9195 | 0.8058 | 2.311 | 0.8396 | 12.787 | 0.2943 | 2.478 | 0.8307 | 2.239 | 0.8391 |
| 334 | 2QZJ_A | 1.267 | 0.9203 | 2.9665 | 0.8805 | 1.167 | 0.9286 | 13.354 | 0.291 | 1.192 | 0.9279 | 1.199 | 0.9277 |
| 335 | 2R8W_B | 2.38 | 0.9101 | 5.097 | 0.6711 | 1.873 | 0.9344 | 1.69 | 0.9413 | 1.728 | 0.94 | 1.635 | 0.9446 |
| 336 | 2RCC_B | 1.725 | 0.9346 | 3.988 | 0.776 | 1.55 | 0.946 | 1.604 | 0.9413 | 1.572 | 0.9443 | 1.587 | 0.9437 |
| 337 | 2RD5_D | 13.344 | 0.5602 | 8.6495 | 0.5824 | 11.966 | 0.6198 | 12.273 | 0.6076 | 12.245 | 0.612 | 12.684 | 0.6078 |
| 338 | 2RLD_C | 3.482 | 0.8801 | 3.25 | 0.8188 | 3.293 | 0.8837 | 3.091 | 0.8807 | 3.432 | 0.8797 | 2.824 | 0.8829 |
| 339 | 2UUX_A | 4.057 | 0.5962 | 4.092 | 0.6053 | 4.586 | 0.5663 | 4.975 | 0.4315 | 4.32 | 0.6179 | 7.35 | 0.6326 |
| 340 | 2VDX_A | 5.995 | 0.7875 | 23.7815 | 0.3546 | 5.299 | 0.9048 | 17.706 | 0.343 | 4.085 | 0.927 | 4.041 | 0.9299 |
| 341 | 2VUL_A | 3.463 | 0.7873 | 6.34 | 0.5323 | 2.699 | 0.8385 | 2.525 | 0.8438 | 2.657 | 0.834 | 2.349 | 0.8529 |
| 342 | 2WCW_B | 2.873 | 0.8238 | 3.8885 | 0.6803 | 2.701 | 0.8341 | 2.684 | 0.8304 | 2.774 | 0.8411 | 2.858 | 0.8327 |
| 343 | 2WGP_A | 6.248 | 0.7978 | 5.055 | 0.7248 | 4.998 | 0.83 | 13.22 | 0.3101 | 5.036 | 0.8278 | 4.674 | 0.8308 |
| 344 | 2WZR_1 | 13.94 | 0.4861 | 20.6925 | 0.3949 | 14.639 | 0.4765 | 9.616 | 0.554 | 18.001 | 0.4997 | 10.502 | 0.5414 |
| 345 | 2XGY_A | 6.443 | 0.6448 | 5.9605 | 0.5841 | 4.487 | 0.7169 | 4.84 | 0.709 | 4.394 | 0.7352 | 4.462 | 0.7274 |
| 346 | 2XW2_A | 5.649 | 0.7396 | 24.497 | 0.2791 | 4.41 | 0.8185 | 17.459 | 0.6169 | 4.172 | 0.826 | 4.272 | 0.8277 |
| 347 | 2Z3B_A | 3.243 | 0.8334 | 6.959 | 0.5519 | 2.154 | 0.8771 | 2.077 | 0.8779 | 2.132 | 0.878 | 2.138 | 0.8777 |
| 348 | 2ZMZ_B | 4.235 | 0.7111 | 3.8085 | 0.6418 | 4.376 | 0.7233 | 3.928 | 0.731 | 4.215 | 0.7247 | 4.051 | 0.7233 |
| 349 | 3A04_A | 4.272 | 0.827 | 9.962 | 0.5202 | 4.127 | 0.8603 | 4.788 | 0.8645 | 3.956 | 0.8737 | 4.019 | 0.8714 |
| 350 | 3ALU_A | 5.887 | 0.6928 | 6.242 | 0.5472 | 5.145 | 0.7339 | 5.046 | 0.7225 | 4.939 | 0.7446 | 4.802 | 0.7476 |
| 351 | 3BDB_A | 6.242 | 0.6978 | 6.611 | 0.5844 | 6.681 | 0.7183 | 6.094 | 0.7186 | 5.083 | 0.7262 | 4.79 | 0.7396 |
| 352 | 3CAE_A | 25.729 | 0.544 | 24.182 | 0.4614 | 25.884 | 0.5715 | 25.661 | 0.5491 | 25.928 | 0.5364 | 25.86 | 0.5369 |
| 353 | 3CG4_A | 2.69 | 0.8332 | 3.459 | 0.7654 | 2.584 | 0.8536 | 2.561 | 0.8485 | 2.539 | 0.8584 | 2.524 | 0.8579 |
| 354 | 3CHB_D | 13.061 | 0.3387 | 12.365 | 0.3438 | 8.189 | 0.382 | 8.055 | 0.38 | 9.851 | 0.4039 | 9.952 | 0.4187 |
| 355 | 3CX5_F | 2.651 | 0.7604 | 3.249 | 0.6346 | 2.912 | 0.7411 | 3.105 | 0.6661 | 2.844 | 0.7444 | 2.725 | 0.7361 |
| 356 | 3E6M_E | 5.458 | 0.751 | 9.668 | 0.6293 | 5.553 | 0.7341 | 5.803 | 0.688 | 5.921 | 0.7321 | 5.767 | 0.7314 |
| 357 | 3E9T_D | 2.95 | 0.7913 | 4.759 | 0.5758 | 1.662 | 0.8762 | 1.728 | 0.8669 | 1.693 | 0.8779 | 1.605 | 0.8778 |
| 358 | 3EOD_A | 3.127 | 0.8367 | 3.97 | 0.7592 | 2.208 | 0.856 | 12.406 | 0.2897 | 1.858 | 0.8726 | 1.902 | 0.8639 |
| 359 | 3F4W_A | 3.427 | 0.8803 | 4.14 | 0.7564 | 2.143 | 0.9066 | 1.964 | 0.9085 | 2.071 | 0.9048 | 1.991 | 0.9054 |
| 360 | 3F8L_A | 2.51 | 0.8382 | 4.301 | 0.6542 | 3.246 | 0.8551 | 14.607 | 0.3009 | 3.214 | 0.8603 | 3.131 | 0.869 |
| 361 | 3G20_B | 7.558 | 0.6301 | 7.675 | 0.5763 | 5.532 | 0.6794 | 6.992 | 0.6647 | 5.092 | 0.6861 | 4.698 | 0.6911 |
| 362 | 3GMX_A | 14.947 | 0.3801 | 8.7665 | 0.3825 | 18.173 | 0.397 | 18.688 | 0.3794 | 18.814 | 0.4026 | 19.363 | 0.4141 |
| 363 | 3H05_B | 3.346 | 0.8238 | 6.2055 | 0.578 | 3.245 | 0.8134 | 3.417 | 0.8117 | 3.377 | 0.8217 | 3.287 | 0.8122 |

Continued on next page

Continue to last page

| No. | PDB | IddFold | | Rosetta_D | | IddFold_relax | | trRosetta_D | | IddFold_relax2 | | trRosetta | |
|---|---|---|---|---|---|---|---|---|---|---|---|---|---|
| | | RMSD | TMscore | RMSD | TMscore | RMSD | TMscore | RMSD | TMscore | RMSD | TMscore | RMSD | TMscore |
| 364 | 3HGI_A | 10.538 | 0.5629 | 12.144 | 0.4352 | 10.697 | 0.755 | 15.362 | 0.3309 | 10.634 | 0.7578 | 10.266 | 0.7636 |
| 365 | 3I9V_7 | 4.908 | 0.6131 | 5.63 | 0.5533 | 4.623 | 0.6727 | 4.334 | 0.6797 | 5.291 | 0.6677 | 5.046 | 0.6698 |
| 366 | 3IAM_4 | 8.878 | 0.7297 | 16.4465 | 0.3845 | 4.369 | 0.8649 | 4.891 | 0.8646 | 8.422 | 0.8446 | 8.575 | 0.8557 |
| 367 | 3JB9_L | 2.717 | 0.9158 | 4.817 | 0.6914 | 2.161 | 0.9412 | 2.103 | 0.9421 | 2.086 | 0.9442 | 2.07 | 0.9448 |
| 368 | 3JU7_A | 5.435 | 0.7795 | 20.8995 | 0.3898 | 2.848 | 0.8772 | 8.587 | 0.7184 | 2.703 | 0.883 | 2.742 | 0.8795 |
| 369 | 3JZ4_A | 7.833 | 0.7284 | 26.7285 | 0.3697 | 7.625 | 0.8735 | 18.384 | 0.3522 | 7.352 | 0.8919 | 7.642 | 0.8976 |
| 370 | 3LQV_B | 6.581 | 0.6839 | 6.887 | 0.5907 | 5.932 | 0.7027 | 7.446 | 0.6573 | 5.634 | 0.6737 | 5.433 | 0.6801 |
| 371 | 3M1N_B | 13.704 | 0.4622 | 17.786 | 0.4176 | 14.209 | 0.4815 | 14.17 | 0.4769 | 13.623 | 0.4873 | 20.063 | 0.4704 |
| 372 | 3MAT_A | 4.333 | 0.8442 | 6.905 | 0.5885 | 2.626 | 0.917 | 2.828 | 0.925 | 2.631 | 0.9259 | 2.577 | 0.9263 |
| 373 | 3MQK_C | 1.487 | 0.8528 | 2.2535 | 0.7922 | 1.498 | 0.856 | 1.785 | 0.8296 | 1.382 | 0.871 | 1.503 | 0.8632 |
| 374 | 3MX3_A | 8.467 | 0.7219 | 29.5115 | 0.3991 | 3.458 | 0.8723 | 3.39 | 0.8721 | 3.455 | 0.8762 | 3.312 | 0.8773 |
| 375 | 3N0X_A | 6.772 | 0.7203 | 23.711 | 0.4259 | 3.224 | 0.8491 | 3.166 | 0.8528 | 3.049 | 0.8587 | 3.068 | 0.8573 |
| 376 | 3N1G_C | 2.313 | 0.8271 | 3.2455 | 0.7201 | 1.826 | 0.8625 | 2.145 | 0.8446 | 2.032 | 0.8533 | 1.98 | 0.853 |
| 377 | 3N9U_C | 5.87 | 0.728 | 5.87 | 0.6672 | 5.473 | 0.7416 | 5.508 | 0.7385 | 5.837 | 0.7358 | 5.696 | 0.7469 |
| 378 | 3NGF_A | 2.727 | 0.8847 | 5.575 | 0.6009 | 1.836 | 0.9269 | 1.696 | 0.9352 | 1.764 | 0.9306 | 1.701 | 0.9339 |
| 379 | 3O61_A | 14.268 | 0.6416 | 15.667 | 0.4205 | 13.822 | 0.6925 | 13.869 | 0.6963 | 13.454 | 0.6993 | 13.619 | 0.6916 |
| 380 | 3OUJ_A | 7.119 | 0.7548 | 7.655 | 0.5227 | 5.898 | 0.8218 | 5.976 | 0.8195 | 6.05 | 0.8125 | 5.844 | 0.8204 |
| 381 | 3P8B_A | 2.54 | 0.6162 | 3.859 | 0.4099 | 1.754 | 0.7635 | 2.132 | 0.7162 | 1.999 | 0.7479 | 2.021 | 0.7451 |
| 382 | 3PD2_A | 4.311 | 0.7392 | 5.892 | 0.6272 | 2.493 | 0.8349 | 2.369 | 0.8496 | 2.448 | 0.8352 | 2.874 | 0.8285 |
| 383 | 3QFM_A | 4.774 | 0.7893 | 6.648 | 0.5691 | 3.348 | 0.8479 | 3.051 | 0.8672 | 3.126 | 0.8566 | 3.032 | 0.8645 |
| 384 | 3QNS_A | 9.5 | 0.6487 | 20.746 | 0.448 | 3.264 | 0.8821 | 2.832 | 0.8999 | 3.141 | 0.8949 | 3.056 | 0.9012 |
| 385 | 3QU3_A | 4.292 | 0.6704 | 6.453 | 0.5317 | 3.941 | 0.6661 | 3.99 | 0.6697 | 3.837 | 0.6817 | 3.86 | 0.6767 |
| 386 | 3R9T_A | 18.28 | 0.7059 | 16.585 | 0.454 | 17.846 | 0.767 | 17.724 | 0.7745 | 17.655 | 0.7788 | 17.62 | 0.7759 |
| 387 | 3ROT_A | 3.114 | 0.8824 | 8.209 | 0.5641 | 1.84 | 0.9345 | 1.744 | 0.9365 | 1.799 | 0.9369 | 1.741 | 0.9396 |
| 388 | 3SDL_B | 31.314 | 0.2557 | 11.713 | 0.3485 | 12.5 | 0.3724 | 9.747 | 0.3528 | 11.012 | 0.3534 | 13.611 | 0.3414 |
| 389 | 3TEO_A | 17.01 | 0.5642 | 13.426 | 0.4813 | 14.612 | 0.5856 | 14.054 | 0.6036 | 14.14 | 0.6159 | 15.178 | 0.6148 |
| 390 | 3UE6_E | 6.721 | 0.732 | 5.937 | 0.6126 | 6.586 | 0.7285 | 6.519 | 0.7234 | 6.796 | 0.7281 | 5.837 | 0.7284 |
| 391 | 3V1O_A | 5.298 | 0.7686 | 5.894 | 0.5651 | 4.4 | 0.7827 | 4.857 | 0.7845 | 4.723 | 0.7892 | 4.848 | 0.7861 |
| 392 | 3W1Z_D | 10.677 | 0.7016 | 7.854 | 0.5582 | 10.638 | 0.7455 | 10.74 | 0.3045 | 10.697 | 0.7265 | 10.86 | 0.717 |
| 393 | 3X0G_A | 3.229 | 0.6943 | 5.626 | 0.5359 | 3.149 | 0.7084 | 6.358 | 0.6235 | 3.276 | 0.7066 | 3.411 | 0.706 |
| 394 | 3X15_A | 7.866 | 0.5513 | 13.557 | 0.4799 | 19.317 | 0.5524 | 19.107 | 0.5432 | 19.344 | 0.5607 | 19.153 | 0.545 |
| 395 | 3ZGF_A | 18.075 | 0.7035 | 21.962 | 0.3434 | 4.969 | 0.7966 | 10.456 | 0.7842 | 8.033 | 0.8005 | 9.112 | 0.8033 |
| 396 | 3ZO5_A | 2.982 | 0.8521 | 7.601 | 0.5406 | 2.178 | 0.9098 | 2.349 | 0.9057 | 2.14 | 0.9115 | 1.963 | 0.9179 |
| 397 | 4AIH_A | 1.821 | 0.8795 | 2.4435 | 0.849 | 2.254 | 0.8374 | 2.394 | 0.837 | 2.097 | 0.8505 | 2.018 | 0.8584 |
| 398 | 4ASW_C | 2.019 | 0.8112 | 3.1185 | 0.698 | 2.087 | 0.8262 | 1.99 | 0.8278 | 2.005 | 0.8313 | 1.965 | 0.8257 |
| 399 | 4AUB_C | 4.471 | 0.847 | 8.226 | 0.5579 | 3.829 | 0.8818 | 17.804 | 0.3254 | 3.688 | 0.8895 | 3.698 | 0.8881 |
| 400 | 4B0M_A | 7.097 | 0.7388 | 5.826 | 0.5828 | 6.293 | 0.7525 | 6.129 | 0.7457 | 6.367 | 0.7499 | 6.295 | 0.7512 |
| 401 | 4CSD_A | 2.543 | 0.8693 | 5.9665 | 0.6568 | 2.194 | 0.8981 | 17.873 | 0.308 | 2.126 | 0.9036 | 2.073 | 0.9087 |
| 402 | 4CXT_A | 2.831 | 0.7863 | 4.41 | 0.6775 | 5.17 | 0.7135 | 10.548 | 0.7315 | 5.284 | 0.7274 | 10.339 | 0.731 |
| 403 | 4DF3_A | 4.142 | 0.751 | 6.03 | 0.5937 | 3.68 | 0.7934 | 4.13 | 0.7801 | 3.681 | 0.7869 | 3.516 | 0.8083 |
| 404 | 4DYW_A | 2.9 | 0.8309 | 4.814 | 0.5427 | 1.623 | 0.8919 | 1.636 | 0.8917 | 1.55 | 0.8967 | 1.594 | 0.8987 |
| 405 | 4ESB_A | 2.205 | 0.8247 | 2.5295 | 0.8043 | 2.515 | 0.8148 | 2.81 | 0.7817 | 2.445 | 0.8235 | 2.488 | 0.8215 |
| 406 | 4F8X_A | 3.329 | 0.8703 | 6.477 | 0.6154 | 2.31 | 0.9141 | 17.116 | 0.3352 | 2.133 | 0.9198 | 1.991 | 0.9265 |
| 407 | 4GDK_A | 2.237 | 0.7945 | 3.6925 | 0.5951 | 1.985 | 0.8201 | 2.307 | 0.8008 | 2.107 | 0.8142 | 2.083 | 0.8142 |
| 408 | 4GF3_A | 6.875 | 0.701 | 7.679 | 0.6101 | 6.813 | 0.7234 | 6.955 | 0.7173 | 6.821 | 0.7267 | 6.838 | 0.7219 |
| 409 | 4GQY_A | 4.537 | 0.7699 | 6.825 | 0.633 | 7.956 | 0.7896 | 7.765 | 0.8028 | 7.6 | 0.8078 | 7.515 | 0.8028 |
| 410 | 4I60_A | 3.513 | 0.7818 | 4.305 | 0.6288 | 2.971 | 0.8083 | 2.835 | 0.823 | 2.927 | 0.8143 | 2.415 | 0.8473 |
| 411 | 4IMH_A | 4.691 | 0.7653 | 20.8865 | 0.4481 | 2.813 | 0.8769 | 15.263 | 0.6206 | 2.672 | 0.887 | 2.653 | 0.8873 |
| 412 | 4IOS_A | 2.719 | 0.7068 | 4.7775 | 0.5604 | 2.633 | 0.7331 | 2.693 | 0.7382 | 2.674 | 0.7367 | 2.649 | 0.7483 |
| 413 | 4J20_A | 2.975 | 0.7364 | 3.954 | 0.6163 | 2.45 | 0.778 | 2.507 | 0.7706 | 2.224 | 0.7923 | 2.563 | 0.7818 |
| 414 | 4JGX_B | 4.376 | 0.7354 | 4.111 | 0.7073 | 4.143 | 0.7526 | 4.121 | 0.7498 | 4.193 | 0.747 | 4.202 | 0.7475 |
| 415 | 4K1F_A | 3.844 | 0.7818 | 5.467 | 0.6438 | 2.919 | 0.8284 | 4.13 | 0.792 | 2.833 | 0.8384 | 2.791 | 0.8364 |

Continued on next page

Continue to last page

| No. | PDB | IddFold | | Rosetta_D | | IddFold_relax | | trRosetta_D | | IddFold_relax2 | | trRosetta | |
|---|---|---|---|---|---|---|---|---|---|---|---|---|---|
| | | RMSD | TMscore | RMSD | TMscore | RMSD | TMscore | RMSD | TMscore | RMSD | TMscore | RMSD | TMscore |
| 416 | 4KA0_A | 2.02 | 0.868 | 3.2585 | 0.7595 | 2.122 | 0.8616 | 2.221 | 0.8518 | 2.113 | 0.865 | 2.171 | 0.8638 |
| 417 | 4KCD_B | 9.076 | 0.5475 | 20.929 | 0.3691 | 7.269 | 0.7142 | 7.531 | 0.7139 | 7.2 | 0.7343 | 7.492 | 0.7346 |
| 418 | 4KKY_X | 6.97 | 0.7109 | 14.2825 | 0.3762 | 3.425 | 0.8771 | 3.102 | 0.8855 | 3.417 | 0.8748 | 3.094 | 0.8847 |
| 419 | 4LE0_B | 1.47 | 0.9099 | 1.6595 | 0.8905 | 1.439 | 0.9108 | 1.55 | 0.8995 | 1.486 | 0.9092 | 1.517 | 0.9081 |
| 420 | 4LMS_A | 10.225 | 0.3811 | 7.244 | 0.4074 | 5.501 | 0.4713 | 8.558 | 0.4247 | 8.48 | 0.4146 | 7.646 | 0.4098 |
| 421 | 4LZ6_A | 4.206 | 0.8312 | 7.303 | 0.6202 | 3.524 | 0.8666 | 3.508 | 0.8682 | 3.396 | 0.8721 | 3.411 | 0.872 |
| 422 | 4M75_F | 2.76 | 0.8142 | 4.377 | 0.6897 | 3.402 | 0.8166 | 4.729 | 0.8323 | 3.364 | 0.8299 | 4.449 | 0.8402 |
| 423 | 4M7O_A | 3.409 | 0.8026 | 7.1695 | 0.5735 | 2.682 | 0.8548 | 2.56 | 0.8569 | 2.415 | 0.8733 | 2.431 | 0.869 |
| 424 | 4MLF_D | 12.066 | 0.274 | 12.013 | 0.2363 | 14.111 | 0.26 | 14.344 | 0.2703 | 14.445 | 0.2844 | 13.042 | 0.3149 |
| 425 | 4MMG_A | 2.23 | 0.8195 | 3.3435 | 0.7793 | 2.005 | 0.8317 | 1.999 | 0.8414 | 1.956 | 0.8443 | 1.881 | 0.8509 |
| 426 | 4MOU_A | 4.271 | 0.839 | 4.983 | 0.7067 | 3.616 | 0.8741 | 17.853 | 0.3362 | 3.502 | 0.8792 | 3.423 | 0.8803 |
| 427 | 4NBI_A | 5.378 | 0.7595 | 6.194 | 0.5792 | 2.798 | 0.8446 | 3.323 | 0.8451 | 2.378 | 0.8649 | 2.551 | 0.8663 |
| 428 | 4OW1_A | 2.634 | 0.7472 | 3.5745 | 0.6599 | 2.267 | 0.7589 | 2.867 | 0.7424 | 2.14 | 0.7854 | 2.19 | 0.7786 |
| 429 | 4Q2O_A | 3.537 | 0.802 | 3.918 | 0.6655 | 4.782 | 0.8016 | 4.94 | 0.8009 | 4.913 | 0.8044 | 4.859 | 0.8133 |
| 430 | 4Q2Q_A | 2.493 | 0.8293 | 4.131 | 0.6723 | 2.816 | 0.8132 | 3.694 | 0.8077 | 2.702 | 0.8119 | 2.973 | 0.8344 |
| 431 | 4Q75_A | 5.12 | 0.8321 | 25.623 | 0.4341 | 2.427 | 0.9246 | 2.158 | 0.9368 | 2.439 | 0.9275 | 5.76 | 0.9038 |
| 432 | 4R67_0 | 2.899 | 0.8392 | 8.2095 | 0.4763 | 2.438 | 0.8605 | 2.374 | 0.8668 | 2.419 | 0.8626 | 2.369 | 0.8648 |
| 433 | 4R8D_A | 8.893 | 0.7312 | 22.468 | 0.3713 | 7.825 | 0.8718 | 14.346 | 0.4754 | 8.068 | 0.8714 | 8.275 | 0.8694 |
| 434 | 4RUV_A | 1.727 | 0.8597 | 1.9015 | 0.8485 | 1.734 | 0.8554 | 1.721 | 0.854 | 1.726 | 0.8608 | 1.698 | 0.8607 |
| 435 | 4UIJ_A | 2.582 | 0.8243 | 2.7535 | 0.8034 | 2.269 | 0.8219 | 2.298 | 0.8287 | 1.984 | 0.8481 | 1.83 | 0.8472 |
| 436 | 4V2O_A | 1.926 | 0.7818 | 1.819 | 0.7875 | 6.744 | 0.531 | 6.851 | 0.5164 | 5.546 | 0.5873 | 4.8 | 0.6044 |
| 437 | 4XEQ_B | 5.997 | 0.7324 | 12.183 | 0.4928 | 1.666 | 0.944 | 1.648 | 0.9453 | 1.63 | 0.9461 | 1.596 | 0.9479 |
| 438 | 4Z6J_A | 3.665 | 0.746 | 5.279 | 0.5904 | 3.875 | 0.7787 | 3.617 | 0.7957 | 3.7 | 0.7901 | 3.706 | 0.7882 |
| 439 | 4ZBY_A | 2.449 | 0.8693 | 5.667 | 0.606 | 2.345 | 0.8763 | 2.264 | 0.8762 | 1.977 | 0.9029 | 1.973 | 0.898 |
| 440 | 4ZDO_A | 6.83 | 0.7404 | 22.797 | 0.3853 | 6.215 | 0.7296 | 15.046 | 0.432 | 4.685 | 0.8025 | 4.693 | 0.8253 |
| 441 | 4ZF6_A | 12.151 | 0.6341 | 23.463 | 0.3581 | 5.274 | 0.8489 | 19.993 | 0.3357 | 3.111 | 0.9003 | 3.364 | 0.9066 |
| 442 | 5CJ3_B | 4.053 | 0.7729 | 3.781 | 0.704 | 5.165 | 0.7893 | 5.132 | 0.7853 | 5.063 | 0.7835 | 5.474 | 0.7819 |
| 443 | 5CZ8_Y | 4.227 | 0.784 | 15.694 | 0.5166 | 3.788 | 0.8555 | 3.231 | 0.847 | 3.347 | 0.8614 | 3.598 | 0.8637 |
| 444 | 5E4E_A | 15.244 | 0.3282 | 13.68 | 0.2233 | 14.533 | 0.3076 | 6.572 | 0.4909 | 6.408 | 0.4887 | 5.645 | 0.4985 |
| 445 | 5EKT_A | 6.245 | 0.7471 | 7.161 | 0.5392 | 3.461 | 0.8317 | 2.705 | 0.8486 | 3.127 | 0.8449 | 2.987 | 0.8551 |
| 446 | 5IAO_A | 3.717 | 0.7987 | 6.201 | 0.5601 | 3.457 | 0.8123 | 12.534 | 0.3382 | 3.507 | 0.8171 | 3.444 | 0.8196 |
| 447 | 5IZB_A | 11.29 | 0.5708 | 14.791 | 0.5178 | 13.248 | 0.5559 | 13.126 | 0.5446 | 13.769 | 0.5893 | 12.784 | 0.567 |
| 448 | 5JTM_A | 10.167 | 0.7023 | 9.087 | 0.5616 | 10.46 | 0.7447 | 10.934 | 0.6851 | 10.381 | 0.7551 | 10.434 | 0.7548 |
| 449 | 5L38_A | 2.156 | 0.8711 | 6.409 | 0.7373 | 2.02 | 0.8748 | 2.443 | 0.855 | 2.108 | 0.8647 | 2.223 | 0.8683 |
| 450 | 5L8R_D | 9.338 | 0.5479 | 14.941 | 0.3931 | 6.254 | 0.6244 | 11.497 | 0.5763 | 6.134 | 0.6348 | 8.082 | 0.6128 |
| 451 | 5LAI_L | 5.406 | 0.7895 | 7.041 | 0.5199 | 3.744 | 0.8329 | 3.88 | 0.8404 | 3.468 | 0.8348 | 3.722 | 0.8367 |
| 452 | 5LXE_A | 6.453 | 0.8036 | 7.243 | 0.6147 | 3.295 | 0.8677 | 3.133 | 0.8778 | 3.18 | 0.8746 | 3.146 | 0.8776 |
| 453 | 5O2V_A | 4.179 | 0.7641 | 2.7745 | 0.751 | 3.6 | 0.7752 | 3.632 | 0.7659 | 3.323 | 0.7861 | 3.489 | 0.7717 |
| 454 | 5O8G_A | 4.427 | 0.7688 | 5.1215 | 0.7238 | 3.905 | 0.7809 | 4.411 | 0.7603 | 4.153 | 0.7768 | 3.46 | 0.772 |
| 455 | 5T17_A | 2.696 | 0.7186 | 2.731 | 0.747 | 2.576 | 0.7319 | 2.571 | 0.7366 | 2.563 | 0.7318 | 2.535 | 0.7368 |
| 456 | 5TMF_E | 7.115 | 0.5779 | 7.842 | 0.5126 | 5.672 | 0.6036 | 5.975 | 0.5702 | 5.699 | 0.5906 | 5.788 | 0.5894 |
| 457 | 5TUV_B | 14.051 | 0.4494 | 10.706 | 0.4218 | 11.537 | 0.4064 | 13.002 | 0.3847 | 11.411 | 0.4209 | 13.556 | 0.4449 |
| 458 | 5VAZ_B | 3.066 | 0.8643 | 5.221 | 0.7021 | 2.655 | 0.8863 | 2.628 | 0.888 | 2.67 | 0.8848 | 2.669 | 0.8851 |
| 459 | 5W71_A | 9.894 | 0.7602 | 22.338 | 0.4143 | 4.611 | 0.8544 | 4.757 | 0.8537 | 4.67 | 0.8571 | 5.561 | 0.8543 |
| 460 | 5WSE_A | 4.203 | 0.7887 | 3.93 | 0.7258 | 3.176 | 0.8073 | 2.58 | 0.813 | 3.369 | 0.8103 | 2.308 | 0.8369 |
| 461 | 6AQ3_B | 3.647 | 0.7953 | 4.719 | 0.6604 | 3.144 | 0.8095 | 3.877 | 0.7792 | 3.35 | 0.7989 | 3.515 | 0.7953 |
| 462 | 6BQE_A | 5.631 | 0.7783 | 8.738 | 0.5079 | 4.516 | 0.8822 | 4.15 | 0.8847 | 4.466 | 0.8927 | 4.426 | 0.8943 |

Table S3: Results of the first model predicted by IddFold-AB and IddFold-A on the benchmark dataset.

| No. | PDB | IddFold-AB | | IddFold-A | | No. | PDB | IddFold-AB | | IddFold-A | |
|---|---|---|---|---|---|---|---|---|---|---|---|
| | | RMSD | TM-score | RMSD | TM-score | | | RMSD | TM-score | RMSD | TM-score |
| 1 | 1A3A_C | 7.264 | 0.6868 | 4.217 | 0.6749 | 53 | 1F15_C | 23.684 | 0.2099 | 21.002 | 0.2538 |
| 2 | 1A40_A | 10.9 | 0.4242 | 13.799 | 0.4138 | 54 | 1F1E_A | 12.096 | 0.4682 | 5.784 | 0.6149 |
| 3 | 1A6L_A | 5.86 | 0.6721 | 4.574 | 0.6304 | 55 | 1F2R_I | 6.613 | 0.6468 | 5.947 | 0.6519 |
| 4 | 1A7D_A | 4.285 | 0.7953 | 3.927 | 0.7619 | 56 | 1F3Y_A | 10.203 | 0.4481 | 10.24 | 0.4318 |
| 5 | 1A91_A | 3.955 | 0.5083 | 4.215 | 0.4899 | 57 | 1F43_A | 7.399 | 0.5761 | 7.387 | 0.5718 |
| 6 | 1ABT_A | 5.648 | 0.488 | 6.17 | 0.4936 | 58 | 1F5X_A | 8.183 | 0.7502 | 11.399 | 0.5415 |
| 7 | 1ABV_A | 2.951 | 0.7383 | 2.415 | 0.7842 | 59 | 1F93_A | 3.172 | 0.8027 | 2.971 | 0.6886 |
| 8 | 1AHK_A | 6.904 | 0.4415 | 6.784 | 0.4642 | 60 | 1F98_A | 8.563 | 0.6423 | 9.225 | 0.6015 |
| 9 | 1AK6_A | 8.209 | 0.5233 | 9.239 | 0.4975 | 61 | 1F9P_A | 9.916 | 0.6325 | 10.908 | 0.5226 |
| 10 | 1AKP_A | 7.157 | 0.4818 | 6.148 | 0.487 | 62 | 1FC3_A | 2.82 | 0.7916 | 3.129 | 0.7521 |
| 11 | 1AOX_A | 3.867 | 0.7894 | 5.97 | 0.5496 | 63 | 1FCA_A | 1.446 | 0.782 | 1.203 | 0.8322 |
| 12 | 1AP7_A | 3.65 | 0.7036 | 3.198 | 0.7548 | 64 | 1FEX_A | 3.258 | 0.6337 | 2.537 | 0.677 |
| 13 | 1AUU_A | 3.331 | 0.6603 | 4.405 | 0.6 | 65 | 1FHT_A | 13.552 | 0.6334 | 14.425 | 0.556 |
| 14 | 1AX8_A | 9.431 | 0.6642 | 8.442 | 0.5619 | 66 | 1FJG_H | 4.3 | 0.6748 | 6.478 | 0.6085 |
| 15 | 1AZ0_A | 7.986 | 0.5974 | 10.75 | 0.4682 | 67 | 1FMB_A | 5.349 | 0.6451 | 6.731 | 0.4778 |
| 16 | 1B4B_A | 3.099 | 0.7918 | 3.012 | 0.7932 | 68 | 1FR0_A | 4.026 | 0.7224 | 3.594 | 0.7358 |
| 17 | 1B4R_A | 3.05 | 0.6549 | 3.635 | 0.6312 | 69 | 1FRD_A | 5.349 | 0.6451 | 4.116 | 0.6669 |
| 18 | 1B4U_A | 7.348 | 0.598 | 10.166 | 0.6244 | 70 | 1FSP_A | 3.295 | 0.8346 | 3.501 | 0.8178 |
| 19 | 1B8Q_A | 11.845 | 0.4503 | 14.184 | 0.4389 | 71 | 1FSU_A | 16.863 | 0.4478 | 18.736 | 0.3525 |
| 20 | 1BE3_J | 3.989 | 0.5613 | 3.914 | 0.5106 | 72 | 1FW9_A | 9.808 | 0.5422 | 11.372 | 0.5107 |
| 21 | 1BG8_A | 3.378 | 0.7556 | 4.207 | 0.6634 | 73 | 1FY2_A | 2.897 | 0.8603 | 7.695 | 0.5903 |
| 22 | 1BGF_A | 2.921 | 0.7622 | 3.287 | 0.735 | 74 | 1G2R_A | 2.132 | 0.8038 | 3.176 | 0.6906 |
| 23 | 1BJX_A | 8.607 | 0.7441 | 9.764 | 0.7291 | 75 | 1G8Q_A | 3.728 | 0.6825 | 8.24 | 0.5258 |
| 24 | 1BUO_A | 3.1 | 0.7508 | 5.248 | 0.6557 | 76 | 1GGS_A | 11.149 | 0.6951 | 10.425 | 0.6644 |
| 25 | 1C03_A | 12.715 | 0.6774 | 12.233 | 0.6324 | 77 | 1GME_A | 16.239 | 0.481 | 15.011 | 0.4625 |
| 26 | 1C1K_A | 8.795 | 0.4728 | 11.248 | 0.3861 | 78 | 1GPQ_B | 6.464 | 0.6043 | 9.413 | 0.4737 |
| 27 | 1C41_A | 3.856 | 0.7525 | 4.456 | 0.6907 | 79 | 1GQA_A | 5.119 | 0.7348 | 4.685 | 0.6762 |
| 28 | 1C5E_A | 6.631 | 0.4772 | 5.56 | 0.4814 | 80 | 1GVP_A | 5.287 | 0.6477 | 5.383 | 0.5507 |
| 29 | 1CDB_A | 6.369 | 0.6175 | 6.006 | 0.5317 | 81 | 1GXD_C | 14.782 | 0.4141 | 16.318 | 0.3565 |
| 30 | 1CF7_B | 2.853 | 0.7064 | 2.698 | 0.7261 | 82 | 1GXL_A | 11.256 | 0.6047 | 11.462 | 0.3479 |
| 31 | 1CQA_A | 3.558 | 0.7305 | 4.2 | 0.643 | 83 | 1GYH_A | 8.563 | 0.5698 | 11.813 | 0.4598 |
| 32 | 1CTO_A | 5.959 | 0.5516 | 7.377 | 0.4752 | 84 | 1H4L_D | 3.713 | 0.737 | 5.882 | 0.5852 |
| 33 | 1CXZ_B | 2.961 | 0.8188 | 2.348 | 0.8349 | 85 | 1H8E_H | 3.624 | 0.6774 | 4.619 | 0.5725 |
| 34 | 1D6T_A | 4.325 | 0.6951 | 5.78 | 0.6423 | 86 | 1H9E_A | 6.395 | 0.5333 | 6.473 | 0.5139 |
| 35 | 1D8B_A | 3.199 | 0.6721 | 4.157 | 0.6453 | 87 | 1H9F_A | 5.195 | 0.5548 | 4.735 | 0.5618 |
| 36 | 1DBF_A | 5.064 | 0.6969 | 6.476 | 0.6091 | 88 | 1HBG_A | 2.616 | 0.8183 | 3.719 | 0.7109 |
| 37 | 1DCF_A | 4.163 | 0.7805 | 5.794 | 0.6598 | 89 | 1HBX_E | 12.032 | 0.5728 | 11.971 | 0.5617 |
| 38 | 1DJ7_A | 7.676 | 0.5365 | 7.336 | 0.4909 | 90 | 1HCD_A | 4.228 | 0.6403 | 4.663 | 0.5908 |
| 39 | 1DL6_A | 15.104 | 0.3865 | 13.715 | 0.3833 | 91 | 1HH8_A | 19.555 | 0.6905 | 16.695 | 0.6925 |
| 40 | 1DOI_A | 9.951 | 0.4367 | 9.726 | 0.4015 | 92 | 1HHV_A | 7.385 | 0.6006 | 8.06 | 0.5691 |
| 41 | 1DP7_P | 2.585 | 0.7554 | 3.369 | 0.6675 | 93 | 1HKQ_A | 4.667 | 0.6982 | 6.516 | 0.6655 |
| 42 | 1DTP_A | 15.358 | 0.2884 | 19.733 | 0.2418 | 94 | 1HKX_E | 6.977 | 0.5531 | 6.981 | 0.5106 |
| 43 | 1DUN_A | 10.872 | 0.5092 | 11.038 | 0.4277 | 95 | 1HL6_D | 7.868 | 0.5118 | 7.654 | 0.5446 |
| 44 | 1DWM_A | 2.942 | 0.6725 | 3.369 | 0.6531 | 96 | 1I27_A | 6.766 | 0.7052 | 6.674 | 0.6692 |
| 45 | 1E2A_A | 1.852 | 0.8306 | 2.41 | 0.7649 | 97 | 1I35_A | 4.466 | 0.5558 | 5.231 | 0.4987 |
| 46 | 1E3Y_A | 4.751 | 0.7224 | 4.959 | 0.6888 | 98 | 1I85_A | 2.895 | 0.7381 | 6.091 | 0.505 |
| 47 | 1E53_A | 3.185 | 0.6154 | 3.566 | 0.5825 | 99 | 1ID2_A | 10.402 | 0.5896 | 10.236 | 0.5503 |
| 48 | 1EGG_B | 8.763 | 0.6037 | 10.842 | 0.5379 | 100 | 1IEZ_A | 7.178 | 0.641 | 9.004 | 0.4293 |
| 49 | 1EKZ_A | 5.006 | 0.6785 | 5.859 | 0.6548 | 101 | 1IJY_A | 3.588 | 0.7006 | 5.512 | 0.5859 |
| 50 | 1ELW_A | 1.459 | 0.8982 | 1.542 | 0.8866 | 102 | 1IOO_A | 9.294 | 0.4631 | 10.848 | 0.4118 |
| 51 | 1EM8_D | 6.426 | 0.6061 | 5.29 | 0.6076 | 103 | 1IPI_A | 3.122 | 0.7488 | 3.148 | 0.7546 |
| 52 | 1EZV_G | 11.8 | 0.4307 | 12.216 | 0.4034 | 104 | 1IQV_A | 4.022 | 0.8093 | 7.317 | 0.6558 |

Continued on next page

Continue to last page

| No. | PDB | IddFold-AB | | IddFold-A | | No. | PDB | IddFold-AB | | IddFold-A | |
|---|---|---|---|---|---|---|---|---|---|---|---|
| | | RMSD | TM-score | RMSD | TM-score | | | RMSD | TM-score | RMSD | TM-score |
| 105 | 1IRS_A | 4.424 | 0.6208 | 6.171 | 0.5263 | 157 | 1MWP_A | 8.279 | 0.5275 | 8.416 | 0.4024 |
| 106 | 1IS7_K | 5.705 | 0.5937 | 7.511 | 0.5663 | 158 | 1MWQ_A | 3.954 | 0.7255 | 3.692 | 0.6665 |
| 107 | 1IUJ_B | 2.614 | 0.8101 | 3.086 | 0.7168 | 159 | 1N12_A | 6.345 | 0.489 | 10.716 | 0.2825 |
| 108 | 1IUY_A | 9.268 | 0.6994 | 9.306 | 0.653 | 160 | 1N13_B | 6.921 | 0.5384 | 6.46 | 0.5818 |
| 109 | 1J1V_A | 1.094 | 0.9242 | 2.046 | 0.8328 | 161 | 1N3G_A | 6.688 | 0.7246 | 6.456 | 0.7132 |
| 110 | 1J3W_A | 4.626 | 0.6972 | 4.893 | 0.6638 | 162 | 1N8V_A | 2.138 | 0.7962 | 2.676 | 0.7537 |
| 111 | 1J8I_A | 15.475 | 0.5492 | 11.259 | 0.515 | 163 | 1NF6_F | 11.087 | 0.7654 | 9.861 | 0.7245 |
| 112 | 1JC7_A | 9.059 | 0.4024 | 9.082 | 0.4431 | 164 | 1NGL_A | 7.524 | 0.5523 | 8.254 | 0.4934 |
| 113 | 1JEI_A | 6.606 | 0.5722 | 6.39 | 0.548 | 165 | 1NKZ_A | 4.595 | 0.5831 | 3.772 | 0.6216 |
| 114 | 1JIW_I | 5.033 | 0.6086 | 5.677 | 0.4989 | 166 | 1NLQ_A | 5.726 | 0.5445 | 9.997 | 0.2839 |
| 115 | 1JJ2_S | 5.598 | 0.6316 | 5.652 | 0.5566 | 167 | 1NOE_A | 7.099 | 0.4533 | 9.308 | 0.3654 |
| 116 | 1JLI_A | 15.831 | 0.3252 | 16.291 | 0.3251 | 168 | 1NPB_A | 7.261 | 0.626 | 9.213 | 0.4518 |
| 117 | 1JMT_A | 4.385 | 0.7206 | 4.989 | 0.6952 | 169 | 1NQJ_B | 7.462 | 0.595 | 7.436 | 0.5497 |
| 118 | 1JO0_A | 2.504 | 0.8201 | 2.885 | 0.7408 | 170 | 1NQZ_A | 9.958 | 0.5989 | 10.438 | 0.5117 |
| 119 | 1JOF_A | 9.394 | 0.5968 | 10.142 | 0.5525 | 171 | 1NR3_A | 18.964 | 0.3008 | 19.928 | 0.2987 |
| 120 | 1JOP_A | 4.943 | 0.6492 | 8.234 | 0.4017 | 172 | 1NRJ_B | 5.413 | 0.7158 | 8.696 | 0.5637 |
| 121 | 1JPY_Y | 15.465 | 0.3553 | 18.445 | 0.293 | 173 | 1NTV_A | 7.2 | 0.5492 | 7.446 | 0.4697 |
| 122 | 1JR8_A | 2.147 | 0.8216 | 2.563 | 0.7715 | 174 | 1NZE_A | 2.891 | 0.8449 | 2.469 | 0.8689 |
| 123 | 1JSG_A | 11.475 | 0.3492 | 12.129 | 0.3389 | 175 | 1O0G_A | 6.187 | 0.4855 | 11.6 | 0.2747 |
| 124 | 1K1Z_A | 8.509 | 0.5171 | 5.406 | 0.5297 | 176 | 1O9G_A | 10.862 | 0.6228 | 10.4 | 0.5104 |
| 125 | 1K3B_A | 6.297 | 0.5173 | 7.62 | 0.5651 | 177 | 1OA8_D | 13.905 | 0.3211 | 18.653 | 0.2405 |
| 126 | 1K3S_A | 4.771 | 0.605 | 3.867 | 0.6249 | 178 | 1OFT_A | 2.982 | 0.7429 | 3.307 | 0.7226 |
| 127 | 1K5D_B | 11.882 | 0.6057 | 9.617 | 0.5233 | 179 | 1OJG_A | 5.166 | 0.5809 | 5.71 | 0.5514 |
| 128 | 1K73_1 | 2.773 | 0.7098 | 5.503 | 0.5715 | 180 | 1OK0_A | 3.918 | 0.5757 | 4.967 | 0.5069 |
| 129 | 1KA8_A | 5.801 | 0.5939 | 7.266 | 0.5327 | 181 | 1OOF_A | 2.256 | 0.8182 | 2.786 | 0.7505 |
| 130 | 1KN6_A | 3.26 | 0.6088 | 4.461 | 0.5071 | 182 | 1OPO_C | 14.942 | 0.2702 | 21.779 | 0.225 |
| 131 | 1KOH_D | 5.613 | 0.7155 | 6.855 | 0.7072 | 183 | 1ORQ_C | 36.157 | 0.424 | 46.879 | 0.3597 |
| 132 | 1KPT_A | 5.918 | 0.5411 | 6.481 | 0.5597 | 184 | 1ORY_A | 2.597 | 0.831 | 2.817 | 0.7942 |
| 133 | 1KQ6_A | 8.666 | 0.6484 | 8.831 | 0.5953 | 185 | 1OX7_A | 4.29 | 0.6707 | 7.048 | 0.5803 |
| 134 | 1KSX_A | 3.443 | 0.7268 | 5.639 | 0.5811 | 186 | 1OZ9_A | 5.173 | 0.6798 | 8.98 | 0.5918 |
| 135 | 1KX5_D | 4.351 | 0.8069 | 8.472 | 0.7612 | 187 | 1P9O_B | 8.748 | 0.6139 | 16.186 | 0.3685 |
| 136 | 1L1D_B | 5.96 | 0.5565 | 9.585 | 0.3963 | 188 | 1PD6_A | 5.593 | 0.6355 | 6.084 | 0.5838 |
| 137 | 1L2P_A | 0.767 | 0.9257 | 1.136 | 0.8674 | 189 | 1PFS_A | 4.744 | 0.4775 | 5.668 | 0.4709 |
| 138 | 1L3G_A | 13.974 | 0.5515 | 15.628 | 0.4754 | 190 | 1PGV_A | 4.318 | 0.7094 | 3.042 | 0.7795 |
| 139 | 1LDD_A | 1.639 | 0.8428 | 2.475 | 0.7949 | 191 | 1PIH_A | 8.667 | 0.3715 | 8.647 | 0.4366 |
| 140 | 1LE2_A | 44.791 | 0.2556 | 48.767 | 0.2563 | 192 | 1PMS_A | 8.837 | 0.5705 | 8.988 | 0.5331 |
| 141 | 1LFU_P | 11.317 | 0.5739 | 10.939 | 0.5843 | 193 | 1PSR_A | 3.656 | 0.7134 | 5.277 | 0.6367 |
| 142 | 1LMI_A | 7.045 | 0.5196 | 6.412 | 0.4898 | 194 | 1PXW_A | 6.02 | 0.6652 | 7.318 | 0.649 |
| 143 | 1LNW_C | 4.547 | 0.8192 | 5.551 | 0.7667 | 195 | 1PZW_A | 3.536 | 0.6456 | 3.198 | 0.6688 |
| 144 | 1LQM_H | 4.992 | 0.5011 | 11.193 | 0.4093 | 196 | 1QFT_A | 5.855 | 0.6067 | 7.356 | 0.4576 |
| 145 | 1LR1_B | 3.919 | 0.5372 | 4.334 | 0.528 | 197 | 1QFW_B | 16.152 | 0.3823 | 14.975 | 0.2712 |
| 146 | 1LWB_A | 9.604 | 0.6022 | 10.099 | 0.5688 | 198 | 1QMA_A | 3.203 | 0.7134 | 3.709 | 0.7086 |
| 147 | 1LYV_A | 8.397 | 0.5892 | 10.438 | 0.5095 | 199 | 1QWT_A | 16.82 | 0.5782 | 22.861 | 0.2413 |
| 148 | 1LZW_B | 4.453 | 0.8207 | 4.113 | 0.7392 | 200 | 1QZG_A | 6.471 | 0.5854 | 9.481 | 0.4263 |
| 149 | 1M7Y_A | 17.398 | 0.4486 | 17.796 | 0.4287 | 201 | 1R5T_A | 5.46 | 0.7185 | 4.513 | 0.6437 |
| 150 | 1MAI_A | 4.524 | 0.6259 | 7.226 | 0.4752 | 202 | 1R5Z_A | 10.412 | 0.4882 | 10.801 | 0.5073 |
| 151 | 1MC2_A | 10.735 | 0.6191 | 5.706 | 0.5736 | 203 | 1R6R_A | 9.914 | 0.5093 | 7.791 | 0.5137 |
| 152 | 1MFQ_C | 8.633 | 0.6354 | 8.766 | 0.6386 | 204 | 1RHX_A | 4.866 | 0.5523 | 4.888 | 0.5208 |
| 153 | 1MHD_A | 6.695 | 0.4787 | 12.041 | 0.4057 | 205 | 1RKX_A | 11.429 | 0.4952 | 14.675 | 0.4644 |
| 154 | 1MKF_A | 26.457 | 0.1929 | 46.166 | 0.1305 | 206 | 1ROW_A | 3.33 | 0.732 | 5.168 | 0.532 |
| 155 | 1MN8_A | 3.736 | 0.6538 | 3.805 | 0.6035 | 207 | 1RTU_A | 9.122 | 0.568 | 7.546 | 0.5551 |
| 156 | 1MT3_A | 6.959 | 0.7194 | 9.038 | 0.5403 | 208 | 1RZ2_A | 16.359 | 0.5529 | 12.761 | 0.4783 |

Continued on next page

Continue to last page

| No. | PDB | IddFold-AB | | IddFold-A | | No. | PDB | IddFold-AB | | IddFold-A | |
|---|---|---|---|---|---|---|---|---|---|---|---|
| | | RMSD | TM-score | RMSD | TM-score | | | RMSD | TM-score | RMSD | TM-score |
| 209 | 1RZ3_A | 11.008 | 0.5998 | 9.026 | 0.492 | 261 | 1ZC9_A | 13.561 | 0.482 | 17.809 | 0.3014 |
| 210 | 1S2D_A | 9.396 | 0.6598 | 8.843 | 0.6103 | 262 | 2A5Y_A | 4.177 | 0.7287 | 6.313 | 0.5764 |
| 211 | 1S3J_A | 3.167 | 0.7642 | 3.369 | 0.7249 | 263 | 2A9U_B | 9.685 | 0.7595 | 10.525 | 0.7874 |
| 212 | 1S56_B | 8.326 | 0.7388 | 8.876 | 0.6704 | 264 | 2AAM_B | 8.103 | 0.8128 | 7.516 | 0.617 |
| 213 | 1S7O_C | 11.197 | 0.5578 | 10.058 | 0.5612 | 265 | 2ACY_A | 2.951 | 0.7315 | 4.495 | 0.5967 |
| 214 | 1S7Z_A | 8.573 | 0.5558 | 5.902 | 0.6146 | 266 | 2AEN_A | 15.457 | 0.2913 | 17.64 | 0.2364 |
| 215 | 1SAU_A | 3.494 | 0.6981 | 4.821 | 0.6414 | 267 | 2APN_A | 7.289 | 0.6307 | 7.593 | 0.5229 |
| 216 | 1SG4_B | 9.48 | 0.7571 | 13.244 | 0.52 | 268 | 2AQ0_A | 7.59 | 0.646 | 7.09 | 0.6294 |
| 217 | 1SMP_I | 3.896 | 0.6649 | 5.395 | 0.5426 | 269 | 2AQS_A | 7.14 | 0.5367 | 9.819 | 0.4377 |
| 218 | 1SPP_B | 4.918 | 0.629 | 5.377 | 0.5395 | 270 | 2AQX_A | 11.243 | 0.4301 | 18.143 | 0.3871 |
| 219 | 1SR8_A | 17.611 | 0.4129 | 20.375 | 0.3099 | 271 | 2BL7_A | 1.633 | 0.8689 | 1.89 | 0.8415 |
| 220 | 1STM_A | 11.423 | 0.3217 | 13.971 | 0.2563 | 272 | 2BS2_C | 8.433 | 0.6793 | 10.452 | 0.5305 |
| 221 | 1SVJ_A | 4.52 | 0.6824 | 4.983 | 0.616 | 273 | 2BSE_A | 7.119 | 0.4917 | 10.322 | 0.3698 |
| 222 | 1TAF_A | 1.326 | 0.8794 | 1.269 | 0.9 | 274 | 2BT9_A | 2.376 | 0.7588 | 3.485 | 0.6687 |
| 223 | 1TEO_A | 10.264 | 0.5916 | 11.548 | 0.5593 | 275 | 2BTD_A | 3.214 | 0.8581 | 5.218 | 0.6802 |
| 224 | 1TJF_B | 6.386 | 0.779 | 7.188 | 0.7445 | 276 | 2BWJ_A | 5.539 | 0.6351 | 8.239 | 0.5835 |
| 225 | 1TLJ_A | 6.025 | 0.6114 | 8.713 | 0.4389 | 277 | 2BYK_D | 1.834 | 0.8497 | 1.844 | 0.8484 |
| 226 | 1TUL_A | 4.613 | 0.538 | 7.241 | 0.4016 | 278 | 2C2F_A | 10.632 | 0.759 | 8.012 | 0.7013 |
| 227 | 1TWU_A | 4.478 | 0.6767 | 4.871 | 0.6178 | 279 | 2C4W_A | 5.682 | 0.691 | 6.405 | 0.6284 |
| 228 | 1TYG_B | 1.83 | 0.7684 | 2.122 | 0.7366 | 280 | 2CDP_A | 6.371 | 0.5306 | 9.553 | 0.4195 |
| 229 | 1TZ0_A | 5.345 | 0.6024 | 5.427 | 0.539 | 281 | 2CEY_A | 8.064 | 0.5518 | 16.318 | 0.3283 |
| 230 | 1U84_A | 2.4 | 0.8338 | 1.758 | 0.8229 | 282 | 2CMX_A | 2.272 | 0.7299 | 2.428 | 0.707 |
| 231 | 1UDD_A | 2.214 | 0.8938 | 4.342 | 0.7665 | 283 | 2CO3_B | 8.217 | 0.4811 | 9.33 | 0.3952 |
| 232 | 1UFB_A | 3.493 | 0.7989 | 5.176 | 0.6907 | 284 | 2CWP_A | 5.508 | 0.6371 | 7.443 | 0.6174 |
| 233 | 1UG4_A | 5.16 | 0.6231 | 5.958 | 0.5639 | 285 | 2CZV_D | 7.088 | 0.7222 | 9.093 | 0.6163 |
| 234 | 1UGI_A | 11.45 | 0.408 | 8.133 | 0.4169 | 286 | 2D0P_B | 3.052 | 0.7624 | 4.638 | 0.6938 |
| 235 | 1UNG_D | 3.671 | 0.7202 | 4.523 | 0.6384 | 287 | 2D5R_B | 2.841 | 0.7832 | 5.129 | 0.6606 |
| 236 | 1USL_C | 6.33 | 0.75 | 5.469 | 0.6776 | 288 | 2DDS_A | 11.491 | 0.5075 | 16.711 | 0.3196 |
| 237 | 1V74_A | 4.039 | 0.6791 | 4.103 | 0.6789 | 289 | 2EWC_B | 5.241 | 0.7351 | 5.366 | 0.6709 |
| 238 | 1VCC_A | 6.121 | 0.5054 | 5.623 | 0.5087 | 290 | 2F22_B | 5.298 | 0.6627 | 6.314 | 0.5784 |
| 239 | 1VCY_A | 11.614 | 0.526 | 9.896 | 0.3594 | 291 | 2FA5_B | 6.275 | 0.6999 | 5.747 | 0.6486 |
| 240 | 1VD0_A | 13.427 | 0.4641 | 10.964 | 0.3634 | 292 | 2FKB_C | 5.411 | 0.6221 | 11.475 | 0.3281 |
| 241 | 1VDW_A | 6.691 | 0.6556 | 7.108 | 0.6128 | 293 | 2GBJ_B | 4.053 | 0.7273 | 4.571 | 0.719 |
| 242 | 1VHG_A | 10.015 | 0.5086 | 10.249 | 0.3916 | 294 | 2GJ3_A | 3.223 | 0.7419 | 3.773 | 0.6721 |
| 243 | 1VKE_E | 4.931 | 0.7088 | 4.261 | 0.7446 | 295 | 2GKC_A | 8.648 | 0.4838 | 10.804 | 0.3727 |
| 244 | 1VLL_A | 9.912 | 0.4707 | 12.551 | 0.506 | 296 | 2H30_A | 6.66 | 0.6385 | 6.956 | 0.5871 |
| 245 | 1VTK_A | 19.085 | 0.3826 | 23.596 | 0.3175 | 297 | 2H8E_A | 5.017 | 0.6014 | 7.413 | 0.4798 |
| 246 | 1VYI_A | 15.696 | 0.353 | 22.054 | 0.2443 | 298 | 2HI3_A | 5.455 | 0.6203 | 4.778 | 0.6403 |
| 247 | 1VYX_A | 5.07 | 0.4463 | 3.803 | 0.458 | 299 | 2HQ7_B | 5.577 | 0.6992 | 5.269 | 0.6193 |
| 248 | 1W1W_E | 2.503 | 0.7697 | 2.176 | 0.7634 | 300 | 2HYB_A | 5.401 | 0.6602 | 5.597 | 0.6149 |
| 249 | 1W53_A | 2.634 | 0.7639 | 2.583 | 0.7633 | 301 | 2ICT_A | 5.183 | 0.7478 | 5.498 | 0.7371 |
| 250 | 1WER_A | 3.426 | 0.8569 | 11.812 | 0.4429 | 302 | 2IEE_A | 12.886 | 0.5393 | 15.435 | 0.3694 |
| 251 | 1WJ8_A | 1.943 | 0.8464 | 1.993 | 0.8346 | 303 | 2IGS_B | 21.094 | 0.217 | 20.866 | 0.2428 |
| 252 | 1WKR_A | 9.207 | 0.5433 | 13.095 | 0.4156 | 304 | 2J4B_B | 2.984 | 0.7239 | 3.539 | 0.7182 |
| 253 | 1WLQ_C | 12.181 | 0.5635 | 11.376 | 0.4936 | 305 | 2J4H_B | 13.658 | 0.3872 | 8.551 | 0.4237 |
| 254 | 1WMH_B | 3.358 | 0.6736 | 2.738 | 0.6882 | 306 | 2J5S_A | 5.757 | 0.7619 | 10.79 | 0.5075 |
| 255 | 1XJA_C | 4.509 | 0.7208 | 8.356 | 0.5618 | 307 | 2J6Z_A | 2.74 | 0.7393 | 2.217 | 0.7957 |
| 256 | 1Y14_A | 11.264 | 0.4893 | 7.1 | 0.5238 | 308 | 2JC5_A | 6.611 | 0.5983 | 7.393 | 0.5396 |
| 257 | 1Y1X_A | 7.91 | 0.6094 | 5.512 | 0.6321 | 309 | 2JLP_D | 12.68 | 0.3046 | 14.941 | 0.2369 |
| 258 | 1YG2_A | 7.314 | 0.4919 | 5.31 | 0.6064 | 310 | 2JP3_A | 10.15 | 0.4223 | 11.299 | 0.4314 |
| 259 | 1YZH_B | 2.756 | 0.8693 | 4.571 | 0.7074 | 311 | 2K9X_A | 5.213 | 0.5726 | 7.324 | 0.4854 |
| 260 | 1Z8R_A | 15.191 | 0.353 | 13.704 | 0.3063 | 312 | 2KBW_A | 4.113 | 0.7436 | 4.824 | 0.6856 |

Continued on next page

Continue to last page

| No. | PDB | IddFold-AB | | IddFold-A | | No. | PDB | IddFold-AB | | IddFold-A | |
|---|---|---|---|---|---|---|---|---|---|---|---|
| | | RMSD | TM-score | RMSD | TM-score | | | RMSD | TM-score | RMSD | TM-score |
| 313 | 2L5P_A | 9.162 | 0.6322 | 5.705 | 0.5632 | 365 | 3I9V_7 | 7.879 | 0.5201 | 7.284 | 0.4654 |
| 314 | 2L74_A | 5.545 | 0.6612 | 6.151 | 0.6736 | 366 | 3IAM_4 | 14.073 | 0.448 | 15.745 | 0.3704 |
| 315 | 2LKP_A | 13.876 | 0.6259 | 11.357 | 0.6306 | 367 | 3JB9_L | 6.716 | 0.7261 | 7.055 | 0.5809 |
| 316 | 2LRB_A | 7.386 | 0.6236 | 7.323 | 0.5127 | 368 | 3JU7_A | 12.393 | 0.3922 | 10.022 | 0.4552 |
| 317 | 2LWP_A | 9.944 | 0.531 | 9.678 | 0.4783 | 369 | 3JZ4_A | 11.548 | 0.5522 | 12.167 | 0.4863 |
| 318 | 2NAZ_A | 4.909 | 0.6557 | 4.079 | 0.6859 | 370 | 3LQV_B | 8.145 | 0.6044 | 9.853 | 0.5623 |
| 319 | 2NCM_A | 2.99 | 0.774 | 5.101 | 0.6523 | 371 | 3M1N_B | 16.379 | 0.4277 | 19.94 | 0.3957 |
| 320 | 2NDP_A | 8.384 | 0.5233 | 4.611 | 0.5456 | 372 | 3MAT_A | 4.165 | 0.8304 | 9.22 | 0.5403 |
| 321 | 2NS9_B | 3.978 | 0.7254 | 6.931 | 0.596 | 373 | 3MQK_C | 2.093 | 0.7837 | 2.92 | 0.6582 |
| 322 | 2NZY_A | 23.372 | 0.2299 | 23.512 | 0.2112 | 374 | 3MX3_A | 13.371 | 0.5462 | 17.167 | 0.4379 |
| 323 | 2O2Y_C | 10.081 | 0.5859 | 10.112 | 0.515 | 375 | 3N0X_A | 12.068 | 0.5358 | 12.982 | 0.3926 |
| 324 | 2O70_F | 5.418 | 0.7517 | 6.689 | 0.621 | 376 | 3N1G_C | 4.096 | 0.6444 | 4.775 | 0.6433 |
| 325 | 2ODM_B | 2.095 | 0.8438 | 1.926 | 0.8613 | 377 | 3N9U_C | 7.207 | 0.6928 | 7.056 | 0.6602 |
| 326 | 2P7L_A | 4.801 | 0.6629 | 4.828 | 0.6385 | 378 | 3NGF_A | 5.546 | 0.6822 | 6.903 | 0.6303 |
| 327 | 2PI2_F | 5.787 | 0.6713 | 5.762 | 0.5654 | 379 | 3O61_A | 15.653 | 0.4732 | 12.309 | 0.4534 |
| 328 | 2PK3_A | 8.852 | 0.5761 | 9.796 | 0.5784 | 380 | 3OUJ_A | 8.274 | 0.7202 | 11.064 | 0.4505 |
| 329 | 2PPV_A | 10.346 | 0.5534 | 10.978 | 0.5059 | 381 | 3P8B_A | 2.407 | 0.6246 | 2.827 | 0.5865 |
| 330 | 2PYB_A | 4.056 | 0.7654 | 4.262 | 0.7465 | 382 | 3PD2_A | 5.865 | 0.6708 | 6.296 | 0.5666 |
| 331 | 2Q2H_A | 8.768 | 0.5121 | 9.676 | 0.4429 | 383 | 3QFM_A | 9.096 | 0.5839 | 11.665 | 0.4716 |
| 332 | 2QTT_A | 5.417 | 0.7172 | 11.516 | 0.4421 | 384 | 3QNS_A | 10.36 | 0.5068 | 14.731 | 0.3434 |
| 333 | 2QVG_A | 3.248 | 0.7844 | 3.178 | 0.7823 | 385 | 3QU3_A | 5.592 | 0.5775 | 6.603 | 0.512 |
| 334 | 2QZJ_A | 2.027 | 0.8565 | 2.27 | 0.8382 | 386 | 3R9T_A | 15.15 | 0.5697 | 16.721 | 0.3551 |
| 335 | 2R8W_B | 5.602 | 0.7139 | 7.074 | 0.6147 | 387 | 3ROT_A | 3.139 | 0.854 | 13.058 | 0.4106 |
| 336 | 2RCC_B | 1.872 | 0.9263 | 4.147 | 0.7358 | 388 | 3SDL_B | 25.133 | 0.2811 | 29.014 | 0.2829 |
| 337 | 2RD5_D | 10.905 | 0.6042 | 12.361 | 0.4996 | 389 | 3TEO_A | 15.008 | 0.5268 | 16.453 | 0.3958 |
| 338 | 2RLD_C | 3.856 | 0.8537 | 3.572 | 0.841 | 390 | 3UE6_E | 6.694 | 0.6774 | 8.328 | 0.5825 |
| 339 | 2UUX_A | 2.807 | 0.6152 | 4.764 | 0.4268 | 391 | 3V1O_A | 8.37 | 0.5361 | 10.64 | 0.4574 |
| 340 | 2VDX_A | 16.542 | 0.3234 | 14.829 | 0.3335 | 392 | 3W1Z_D | 9.752 | 0.6192 | 9.473 | 0.593 |
| 341 | 2VUL_A | 7.52 | 0.5483 | 13.183 | 0.4157 | 393 | 3X0G_A | 4.662 | 0.6015 | 8.998 | 0.469 |
| 342 | 2WCW_B | 3.567 | 0.7835 | 4.427 | 0.6044 | 394 | 3X15_A | 15.958 | 0.4478 | 17.586 | 0.5403 |
| 343 | 2WGP_A | 6.298 | 0.7357 | 6.926 | 0.6595 | 395 | 3ZGF_A | 13.726 | 0.6812 | 18.251 | 0.3605 |
| 344 | 2WZR_1 | 20.453 | 0.4588 | 17.663 | 0.2366 | 396 | 3ZO5_A | 7.884 | 0.5366 | 13.386 | 0.3884 |
| 345 | 2XGY_A | 6.562 | 0.6405 | 7.769 | 0.5878 | 397 | 4AIH_A | 2.885 | 0.8061 | 3.121 | 0.729 |
| 346 | 2XW2_A | 17.089 | 0.3923 | 12.932 | 0.3879 | 398 | 4ASW_C | 3.018 | 0.7218 | 3.073 | 0.625 |
| 347 | 2Z3B_A | 6.772 | 0.5797 | 7.919 | 0.5203 | 399 | 4AUB_C | 9.216 | 0.6234 | 8.67 | 0.5585 |
| 348 | 2ZMZ_B | 4.355 | 0.6547 | 3.881 | 0.6977 | 400 | 4B0M_A | 9.746 | 0.5762 | 8.098 | 0.4882 |
| 349 | 3A04_A | 7.978 | 0.597 | 12.095 | 0.443 | 401 | 4CSD_A | 4.767 | 0.723 | 17.497 | 0.3515 |
| 350 | 3ALU_A | 9.198 | 0.455 | 11.962 | 0.4189 | 402 | 4CXT_A | 4.993 | 0.6709 | 4.418 | 0.632 |
| 351 | 3BDB_A | 8.811 | 0.513 | 8.875 | 0.5406 | 403 | 4DF3_A | 3.953 | 0.7708 | 9.319 | 0.4908 |
| 352 | 3CAE_A | 24.912 | 0.48 | 24.878 | 0.4168 | 404 | 4DYW_A | 4.471 | 0.6611 | 8.035 | 0.4217 |
| 353 | 3CG4_A | 3.206 | 0.7933 | 3.124 | 0.7655 | 405 | 4ESB_A | 2.905 | 0.8028 | 3.2 | 0.7568 |
| 354 | 3CHB_D | 12.512 | 0.3232 | 12.826 | 0.3601 | 406 | 4F8X_A | 6.969 | 0.6541 | 11.507 | 0.4735 |
| 355 | 3CX5_F | 2.82 | 0.7292 | 2.844 | 0.7115 | 407 | 4GDK_A | 3.647 | 0.6595 | 4.898 | 0.5134 |
| 356 | 3E6M_E | 5.492 | 0.6596 | 7.5 | 0.591 | 408 | 4GF3_A | 8.175 | 0.6095 | 9.531 | 0.4578 |
| 357 | 3E9T_D | 4.159 | 0.6219 | 7.091 | 0.5013 | 409 | 4GQY_A | 7.013 | 0.6641 | 9.023 | 0.6657 |
| 358 | 3EOD_A | 3.463 | 0.8021 | 4.619 | 0.8037 | 410 | 4I60_A | 4.66 | 0.6747 | 7.24 | 0.4961 |
| 359 | 3F4W_A | 4.291 | 0.7794 | 5.666 | 0.64 | 411 | 4IMH_A | 8.443 | 0.5442 | 10.705 | 0.4358 |
| 360 | 3F8L_A | 9.428 | 0.632 | 6.539 | 0.5043 | 412 | 4IOS_A | 4.034 | 0.5997 | 5.608 | 0.4607 |
| 361 | 3G20_B | 5.546 | 0.6484 | 9.442 | 0.5026 | 413 | 4J20_A | 5.022 | 0.5721 | 4.302 | 0.5376 |
| 362 | 3GMX_A | 16.354 | 0.3453 | 18 | 0.3489 | 414 | 4JGX_B | 5.04 | 0.6499 | 4.382 | 0.6524 |
| 363 | 3H05_B | 6.362 | 0.622 | 6.858 | 0.5389 | 415 | 4K1F_A | 7.021 | 0.6 | 8.342 | 0.4936 |
| 364 | 3HGI_A | 11.849 | 0.5603 | 17.959 | 0.2901 | 416 | 4KA0_A | 3.174 | 0.7561 | 5.532 | 0.6347 |

Continued on next page

Continue to last page

| No. | PDB | IddFold-AB | | IddFold-A | | No. | PDB | IddFold-AB | | IddFold-A | |
|---|---|---|---|---|---|---|---|---|---|---|---|
| | | RMSD | TM-score | RMSD | TM-score | | | RMSD | TM-score | RMSD | TM-score |
| 417 | 4KCD_B | 14.728 | 0.3762 | 16.438 | 0.2407 | 440 | 4ZDO_A | 18.812 | 0.3392 | 15.678 | 0.3472 |
| 418 | 4KKY_X | 12.133 | 0.5065 | 15.868 | 0.4136 | 441 | 4ZF6_A | 13.061 | 0.4617 | 14.921 | 0.3907 |
| 419 | 4LE0_B | 1.565 | 0.8928 | 1.955 | 0.8495 | 442 | 5CJ3_B | 4.306 | 0.7145 | 4.397 | 0.6726 |
| 420 | 4LMS_A | 11.71 | 0.343 | 7.772 | 0.3428 | 443 | 5CZ8_Y | 8.875 | 0.5395 | 11.192 | 0.4881 |
| 421 | 4LZ6_A | 6.794 | 0.6598 | 17.904 | 0.3153 | 444 | 5E4E_A | 17.258 | 0.3465 | 15.54 | 0.3581 |
| 422 | 4M75_F | 4.442 | 0.713 | 5.599 | 0.7018 | 445 | 5EKT_A | 5.825 | 0.597 | 8.797 | 0.4801 |
| 423 | 4M7O_A | 4.871 | 0.7752 | 12.257 | 0.5159 | 446 | 5IAO_A | 5.869 | 0.6396 | 6.876 | 0.5546 |
| 424 | 4MLF_D | 14.775 | 0.2237 | 11.706 | 0.2486 | 447 | 5IZB_A | 17.036 | 0.5355 | 12.906 | 0.4963 |
| 425 | 4MMG_A | 2.873 | 0.7433 | 3.396 | 0.7186 | 448 | 5JTM_A | 10.193 | 0.568 | 11.581 | 0.5096 |
| 426 | 4MOU_A | 7.501 | 0.6533 | 6.789 | 0.6399 | 449 | 5L38_A | 2.309 | 0.8006 | 2.97 | 0.7455 |
| 427 | 4NBI_A | 8.236 | 0.6096 | 8.047 | 0.5042 | 450 | 5L8R_D | 12.351 | 0.4303 | 13.336 | 0.4332 |
| 428 | 4OW1_A | 2.52 | 0.7239 | 3.625 | 0.5968 | 451 | 5LAI_L | 6.082 | 0.7771 | 10.945 | 0.3958 |
| 429 | 4Q2O_A | 3.261 | 0.7504 | 3.365 | 0.6846 | 452 | 5LXE_A | 8.805 | 0.6403 | 8.686 | 0.5839 |
| 430 | 4Q2Q_A | 3.304 | 0.7555 | 5.886 | 0.6068 | 453 | 5O2V_A | 3.134 | 0.7421 | 5.382 | 0.6981 |
| 431 | 4Q75_A | 10.718 | 0.5191 | 13.345 | 0.3885 | 454 | 5O8G_A | 5.317 | 0.7038 | 5.963 | 0.6374 |
| 432 | 4R67_0 | 9.938 | 0.5369 | 8.598 | 0.501 | 455 | 5T17_A | 2.458 | 0.7369 | 2.646 | 0.7146 |
| 433 | 4R8D_A | 13.493 | 0.42 | 10.985 | 0.4767 | 456 | 5TMF_E | 7.621 | 0.5719 | 8.91 | 0.5091 |
| 434 | 4RUV_A | 2.286 | 0.7996 | 2.19 | 0.7881 | 457 | 5TUV_B | 12.54 | 0.3992 | 13.435 | 0.418 |
| 435 | 4UIJ_A | 3.054 | 0.7152 | 3.857 | 0.7166 | 458 | 5VAZ_B | 7.159 | 0.6711 | 6.199 | 0.6712 |
| 436 | 4V2O_A | 1.476 | 0.8471 | 1.988 | 0.7827 | 459 | 5W71_A | 11.136 | 0.46 | 14.955 | 0.366 |
| 437 | 4XEQ_B | 8.102 | 0.5943 | 15.959 | 0.4174 | 460 | 5WSE_A | 4.466 | 0.7119 | 6.224 | 0.6468 |
| 438 | 4Z6J_A | 6.784 | 0.5806 | 6.4 | 0.4863 | 461 | 6AQ3_B | 5.666 | 0.6394 | 5.283 | 0.5894 |
| 439 | 4ZBY_A | 5.073 | 0.6681 | 7.608 | 0.567 | 462 | 6BQE_A | 5.975 | 0.736 | 13.925 | 0.4422 |

Table S4: TM-score of the first model predicted by IddFold_relax1, IddFold, QUARK, RaptorX-Contact, BAKER-ROSETTASERVER (BAKER_R), and MULTICOM_cluster (M-COM_cluster) on 24 FM targets of CASP13.

| Target | Length | IddFold_relax1 | IddFold | QUARK | RaptorX-Contact | BAKER_R | M-COM_cluster |
|---|---|---|---|---|---|---|---|
| T0950-D1 | 342 | 0.62 | 0.65 | 0.44 | 0.56 | 0.46 | 0.22 |
| T0953s1-D1 | 67 | 0.45 | 0.44 | 0.4 | 0.28 | 0.19 | 0.38 |
| T0953s2-D1 | 44 | 0.33 | 0.32 | 0.36 | 0.18 | 0.33 | 0.2 |
| T0953s2-D2 | 111 | 0.61 | 0.59 | 0.48 | 0.69 | 0.47 | 0.22 |
| T0953s2-D3 | 93 | 0.19 | 0.14 | 0.35 | 0.29 | 0.19 | 0.14 |
| T0955-D1 | 41 | 0.56 | 0.47 | 0.73 | 0.59 | NA | 0.77 |
| T0957s1-D1 | 108 | 0.55 | 0.51 | 0.4 | 0.37 | 0.42 | 0.31 |
| T0957s2-D1 | 155 | 0.75 | 0.7 | 0.53 | 0.65 | 0.48 | 0.51 |
| T0958-D1 | 77 | 0.73 | 0.73 | 0.55 | 0.66 | 0.53 | 0.55 |
| T0960-D2 | 84 | 0.51 | 0.45 | 0.44 | 0.49 | 0.28 | 0.38 |
| T0963-D2 | 82 | 0.53 | 0.46 | 0.46 | 0.48 | 0.36 | 0.28 |
| T0968s1-D1 | 118 | 0.72 | 0.72 | 0.56 | 0.62 | 0.74 | 0.43 |
| T0968s2-D1 | 115 | 0.77 | 0.64 | 0.65 | 0.59 | 0.66 | 0.42 |
| T0969-D1 | 354 | 0.79 | 0.69 | 0.64 | 0.65 | 0.49 | 0.44 |
| T0970-D1 | 85 | 0.73 | 0.69 | 0.51 | 0.54 | 0.4 | 0.33 |
| T0980s1-D1 | 104 | 0.67 | 0.56 | 0.54 | 0.37 | 0.41 | 0.25 |
| T0990-D1 | 76 | 0.64 | 0.57 | 0.58 | 0.4 | 0.37 | 0.36 |
| T0990-D2 | 231 | 0.5 | 0.47 | 0.37 | 0.35 | 0.26 | 0.25 |
| T0990-D3 | 213 | 0.38 | 0.37 | 0.22 | 0.22 | 0.23 | 0.24 |
| T1005-D1 | 326 | 0.69 | 0.67 | 0.7 | 0.7 | 0.7 | 0.69 |
| T1008-D1 | 77 | 0.69 | 0.64 | 0.35 | 0.28 | 0.56 | 0.38 |
| T1021s3-D1 | 166 | 0.69 | 0.64 | 0.64 | 0.66 | 0.5 | 0.5 |
| T1021s3-D2 | 97 | 0.54 | 0.48 | 0.45 | 0.59 | 0.19 | 0.27 |
| T1022s1-D1 | 156 | 0.59 | 0.57 | 0.55 | 0.58 | 0.4 | 0.38 |
| Average | 138 | 0.59 | 0.55 | 0.50 | 0.49 | 0.42 | 0.37 |

Table S5: TM-score of the first model predicted by IddFold_relax1, IddFold, QUARK, RaptorX, BAKER-ROSETTASERVER (BAKER_R), and MULTICOM_cluster (M-COM_cluster) on 20 FM targets of CASP14.

| Target | Length | IddFold_relax1 | IddFold | QUARK | RaptorX | BAKER_R | M-COM_cluster |
|---|---|---|---|---|---|---|---|
| T1027-D1 | 99 | 0.39 | 0.42 | 0.47 | 0.38 | 0.34 | 0.37 |
| T1029-D1 | 125 | 0.45 | 0.46 | 0.47 | 0.45 | 0.49 | 0.46 |
| T1031-D1 | 95 | 0.29 | 0.25 | 0.72 | 0.35 | 0.23 | 0.35 |
| T1033-D1 | 100 | 0.26 | 0.27 | 0.33 | 0.26 | 0.44 | 0.26 |
| T1035-D1 | 102 | 0.59 | 0.42 | 0.83 | 0.4 | 0.26 | 0.77 |
| T1037-D1 | 404 | 0.67 | 0.41 | 0.78 | 0.41 | 0.29 | 0.73 |
| T1038-D1 | 114 | 0.36 | 0.33 | 0.32 | 0.35 | 0.25 | 0.34 |
| T1038-D2 | 76 | 0.66 | 0.66 | 0.6 | 0.47 | 0.59 | 0.56 |
| T1039-D1 | 161 | 0.31 | 0.32 | 0.31 | 0.31 | 0.48 | 0.23 |
| T1040-D1 | 130 | 0.25 | 0.22 | 0.41 | 0.25 | 0.21 | 0.24 |
| T1041-D1 | 242 | 0.71 | 0.62 | 0.76 | 0.69 | 0.4 | 0.65 |
| T1042-D1 | 276 | 0.19 | 0.18 | 0.72 | 0.23 | 0.2 | 0.56 |
| T1043-D1 | 148 | 0.19 | 0.15 | 0.2 | 0.2 | 0.27 | 0.16 |
| T1046s1-D1 | 72 | 0.72 | 0.7 | 0.69 | 0.73 | 0.74 | 0.69 |
| T1049-D1 | 134 | 0.73 | 0.71 | 0.68 | 0.65 | 0.74 | 0.63 |
| T1064-D1 | 92 | 0.22 | 0.18 | 0.25 | 0.22 | 0.25 | 0.25 |
| T1074-D1 | 132 | 0.52 | 0.29 | 0.48 | 0.4 | 0.54 | 0.43 |
| T1080-D1 | 133 | 0.51 | 0.51 | 0.37 | 0.34 | 0.27 | 0.39 |
| T1082-D1 | 75 | 0.56 | 0.49 | 0.7 | 0.52 | 0.63 | 0.55 |
| T1090-D1 | 189 | 0.72 | 0.6 | 0.65 | 0.68 | 0.57 | 0.56 |
| Average | 145 | 0.47 | 0.41 | 0.54 | 0.41 | 0.41 | 0.46 |
